# Molecular characterization of a cytochrome P450 involved in the skeletal diversification of tricyclic coumarins in *Angelica keiskei*

**DOI:** 10.64898/2026.10.02.756137

**Authors:** Nodoka Shinya, Junwen Han, Yoko Kamata, Masahiko Taniguchi, Kenji Miura, Kazufumi Yazaki, Akifumi Sugiyama, Ryosuke Munakata

## Abstract

Tricyclic coumarins (TCs) exhibit defensive properties against herbivorous insects and possess valuable pharmacological properties for humans. They are mainly distributed in the Apiaceae, Rutaceae, Moraceae, and Fabaceae families. The chemical structural diversity of TCs in these families primarily arises from differences in skeletal variety and attachment position of the third ring, as observed in unsaturated furanocoumarins (FCs) and saturated dihydrofuranocoumarins (DHFCs). In TC biosynthesis, the FC and DHFC pathways diverge through distinct oxidation patterns. The product specificity of the reported cytochrome P450 monooxygenase at the branching point suggests that these TC groups are produced in nearly identical ratios; however, some species of the genus *Angelica* (Apiaceae) accumulate DHFCs at higher levels than FCs. In this study, we investigated the enzymatic mechanism responsible for forming these two TC groups using *A. keiskei*, a Japanese native species used as a medicinal and edible herb. Organ-specific transcriptome analysis highlighted a P450 member *CYP71AJ62* that exhibited a close correlation with TC accumulation across organs. Enzymatic characterization demonstrated that CYP71AJ62 predominantly forms a hydroxylated DHFC, vaginidiol, which may promote DHFC over FC skeletal formation in TC biosynthesis. Phylogenetic analysis suggested that *CYP71AJ62* has been derived from a gene responsible for FC formation by functional shift without gene duplication. Subsequent three-dimensional structural modeling and site-directed mutagenesis revealed that two amino acid residues on substrate recognition sites cooperatively determine the product specificity of this P450 enzyme. This study provides new insights into the molecular evolutionary process underlying ring-structure diversification and, more broadly, the adaptation of plants to their environments.

## Introduction

Plants produce a variety of phenylpropanoid molecules, including flavonoids, lignins, and coumarins, many of which contribute to plant adaptation to environmental stresses and exhibit beneficial pharmacological activities for humans. Coumarins undergo diverse chemical modifications, and thus various naturally occurring structures have been reported, in which the coumarin skeleton is often fused to an oxygen-containing ring that differs in terms of both structure and position. Subgroups of coumarins with a five-membered ring (furan or dihydrofuran) fused at the 6,7- or 7,8-positions (Fig. S1) function in plant chemical defense against biotic stresses such as Lepidopteran insects (Berenbaum et al., 1981; Divekar et al., 2022) and phytopathogenic fungi (Afek et al. 1995; Song et al. 2017). These coumarins also exhibit valuable activities for humans, with potential applications in medical fields (Río et al., 2014), such as oncology (Sumorek-Wiadro et al., 2020), dermatology (Sakuntabhai et al., 1995), cardiology (Härmälä et al., 1992), and mammalian infectious diseases (Manderfeld et al., 1997; Kil et al., 2017). In this study, these bioactive compounds are referred to as tricyclic coumarins (TCs). Generally, both their physiological functions and pharmacological activities depend on their chemical structures (Nan et al., 2025). In plants, more than 300 structures have been reported, predominantly from Apiaceae, Rutaceae, Fabaceae, and Moraceae (Murray, 2002).

The biosynthesis of TCs commences with the introduction of an isoprene unit into a simple coumarin skeleton, that is, prenylation of umbelliferone by the UbiA prenyltransferase family (Karamat et al., 2014; Munakata et al., 2016, 2021)(Fig. 1). In this reaction, the prenylation position determines whether a linear or angular product is formed. Umbelliferone 6-dimethylallyltransferase (U6DT) synthesizes demethylsuberosin, an intermediate of linear TCs, whereas U8DT produces osthenol, an angular TCs intermediate. In both linear and angular pathways, demethylsuberosin and osthenol are converted into marmesin and columbianetin, respectively, via oxidative cyclization catalyzed by distinct cytochrome P450s (Zhao et al., 2024). Subsequent side-chain cleavage reactions by cytochrome P450s yield psoralen or angelicin as linear or angular furanocoumarins (FCs)(Larbat et al., 2007, 2009). In the angular pathway, columbianetin and its derivatives retaining the saturated dihydrofuran ring are collectively known as angular dihydrofuranocoumarins (DHFCs). A hydroxylated DHFC is formed as a by-product of the side-chain cleavage from columbianetin to angelicin in apiaceous plants. Vaginidiol, (9R)-hydroxycolumbianetin, has been predicted to be the by-product of angular FC formation (Larbat et al. 2009). Focusing on the hydroxylation of columbianetin, we classified angular DHFCs in apiaceous species (Murray, 2002) into two types based on the presence of a C-9 modification: type A (unmodified at C-9) and type B (hydroxylated at C-9 with additional modifications).

**Figure 1.**
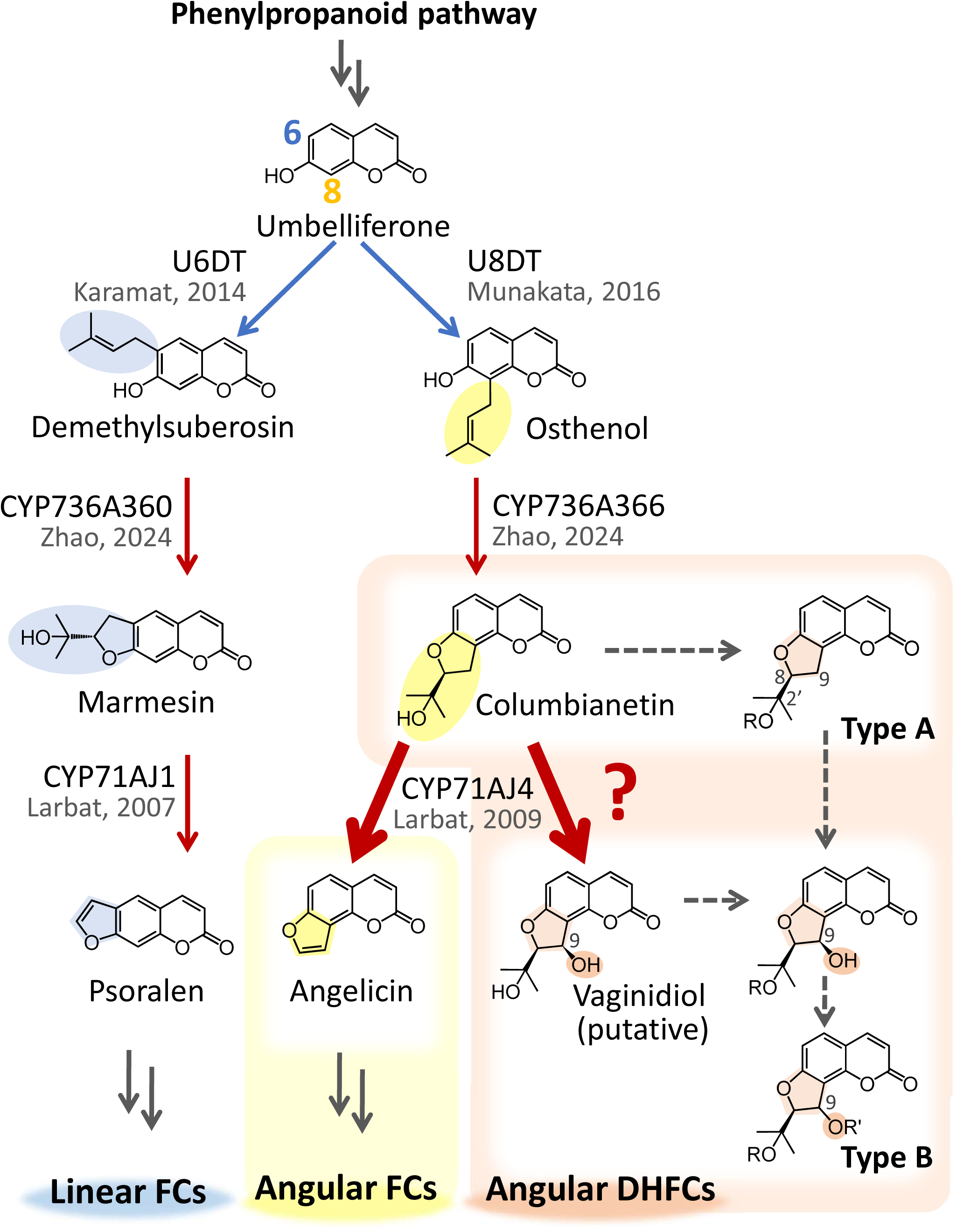
The biosynthetic pathway of the major TCs in Apiaceae. Single arrows represent a single enzymatic reaction, whereas double arrows represent multiple steps. Solid lines indicate known genes; dashed lines indicate unknown genes. Biosynthetic steps catalyzed by cytochrome P450s and PTs are shown in red and blue, respectively. The branching point between angular FCs and DHFCs is shown as a bold arrow. Classification of angular DHFCs in apiaceous species into type A and B, depending on the presence of C-9 modification. R and R’ mean chemical modifications, such as the methyl moiety and the angeloyl moiety. FC, furanocoumarin; DHFC, dihydrofuranocoumarin; U6DT, umbelliferone 6-dimethylallyltransferase; U8DT, umbelliferone 8-dimethylallyltransferase.

The flux from columbianetin may branch into three pathways: conversion to angelicin as an FC via side-chain cleavage, transformation from type A to type B precursor through 9-hydroxylation, and different chemical modifications within type A (Figure 1). The first two reactions are simultaneously catalyzed by a single cytochrome P450, such as CYP71AJ4 in parsnip (*Pastinaca sativa*), and the ratio of their products is almost identical (Larbat et al., 2009). In contrast, metabolic profiling has shown that angular DHFCs can accumulate at higher levels than angular FCs in certain species of the genus *Angelica* (Patyra et al., 2024; Ji et al., 2025). This contradicts previous observations that these biosynthetic enzymes produce both compounds in nearly equal amounts, as mentioned above, and that no precursor-product relationship exists between angular hydroxy DHFC and FC (Larbat et al., 2009). This suggests the existence of unexplored oxidative mechanisms regulating the metabolic flux between FCs and DHFCs (Fig. 1). Further diversification of TC structures involves downstream oxidative modifications, several of which remain enzymatically uncharacterized (Fig. S2).

To address these uncharacterized reaction steps, we selected *Angelica keiskei* as our experimental species. This species is an apiaceous species native to Japan and is used as both a medicinal and edible plant. This is suitable for this study because of its aforementioned metabolite profile, with the accumulation of both linear FCs and exceptionally high levels of angular DHFCs, while angular FCs remain barely detectable (Baba, 2013; Kim et al., 2014). To identify P450 genes involved in TC biosynthesis in *A. keiskei*, transcriptome libraries and metabolite profiles of this species have been generated from different organs. Co-expression analysis and enzymatic characterization were performed to elucidate the unknown oxidative mechanism underlying the formation of angular FCs and DHFCs. By combining phylogenetic analysis and three-dimensional modeling, we also explored amino acid residues that contribute to the catalytic activity in the oxidative mechanism of TC skeletal formation.

## Results

### Organ-specific coumarin accumulation in A. keiskei

To confirm the chemical profile of our *A. keiskei* samples, we quantified TCs (linear FCs, linear DHFCs, angular DHFCs, and angular FCs) and simple coumarins in different *A. keiskei* organs: petiole, main root cortex, main root stele, and lateral root, using an LC-MS/MS system (Fig. S3–S6). Several TCs not previously reported in *A. keiskei* were detected, including heraclenin and heraclenol (linear FCs), apterin (angular DHFC type B), and sphondinol and sphondin (angular FCs). In addition, pimpinellin (angular DHFCs) was detected, which has not been reported in the genus *Angelica* according to SciFinder (https://scifinder-n.cas.org/). Importantly, we observed organ-specific accumulation patterns of TCs. Across all analyzed organs, large amounts of linear FCs and angular DHFCs accumulated at relatively high levels, whereas angular FCs were less abundant. Consistent with previous reports (Sugamoto et al., 2009; Baba, 2013), these three TC groups and simple coumarins accumulated at lower levels in the main root stele than in other organs.

### Co-expression analysis-based selection of candidate cytochrome P450s for TC biosynthesis

We performed *de novo* transcriptome assembly, generating 101,282 contigs from the same samples used for metabolite analysis (Table S1, Fig. S7). To screen for cytochrome P450 genes involved in TC metabolism, co-expression analysis was conducted using three *prenyltransferases (PTs)* as bait genes: the *A. keiskei* homologs of parsnip *U6DT* and *U8DT* (Karamat et al., 2014) at the first reaction step (Fig. 1), and *B5ODT* from *A. keiskei* at a downstream step of TC skeletal formation (Munakata et al., 2016, 2021)(Fig. S2). These bait genes showed lower expression levels in the main root stele than in other organs, which was consistent with the distribution patterns of TCs. Among TC groups, their expression patterns showed higher correlations with the accumulation patterns of linear FC and angular DHFC type B (Fig. 2a). We also observed similar expression patterns between these bait genes and other known TC biosynthetic genes, such as *demethylsuberosin cyclase (DC)* and *O-methyltransferase 1 (OMT1)* (Fig. 2b), suggesting that new TC biosynthetic genes can be screened by co-expression analysis.

**Figure 2.**
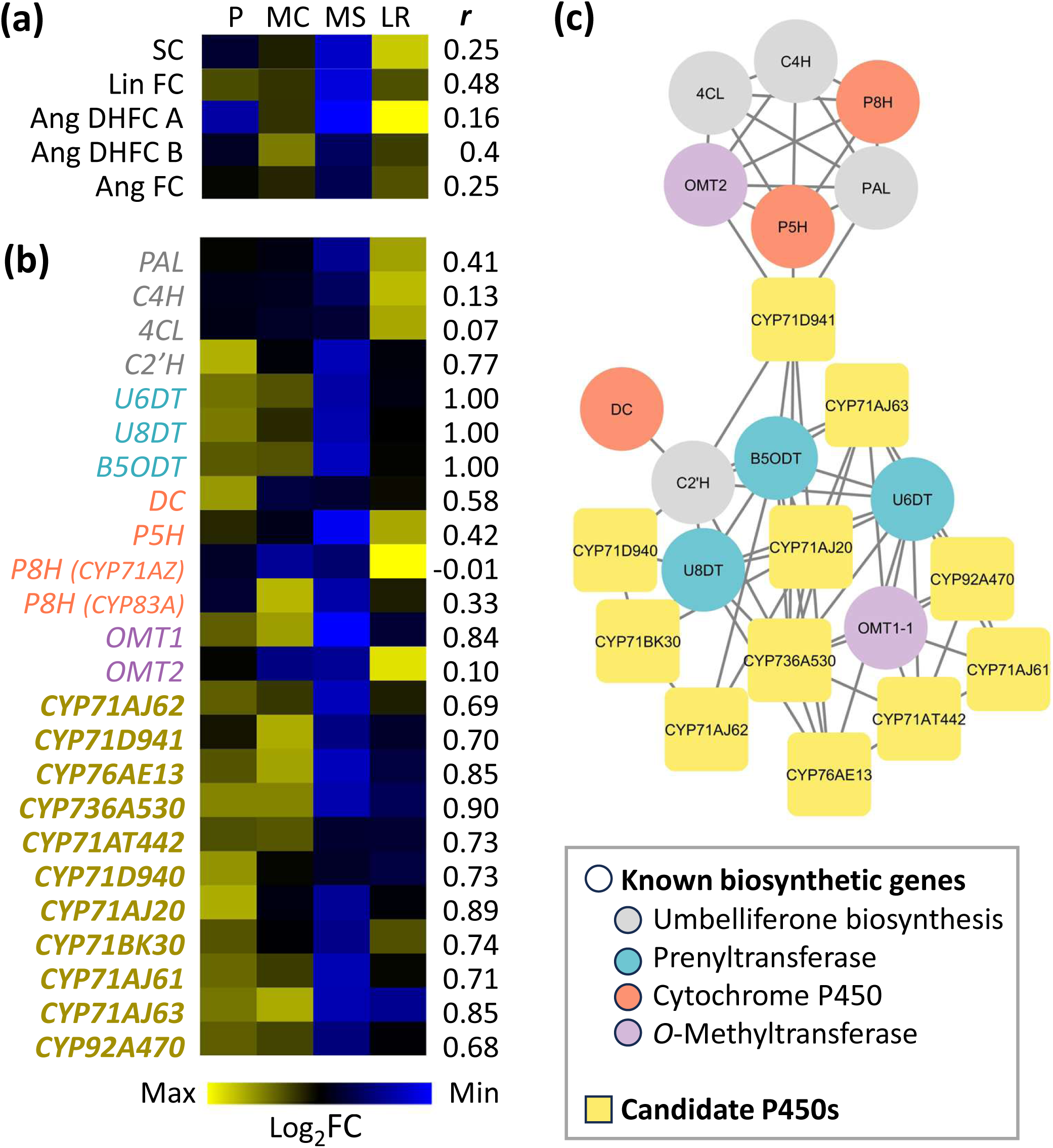
Co-expression analysis of candidate cytochrome P450s and TC biosynthetic genes. Heatmaps of TC accumulation patterns (a) and gene expression patterns (b). The Pearson correlation coefficient with PTs was calculated, and the highest is displayed as *r*. The TPM values of all 24 samples were first normalized using the overall average. For each organ, the mean of the six biological replicates (n = 6) was then calculated, and the resulting values were log₂-transformed to obtain the expression levels. The maximum and minimum values across all cells are represented in yellow and blue, respectively. *PAL, phenylalanine ammonia-lyase; C4H, cinnamate 4-hydroxylase; 4CL, 4-coumaroyl CoA ligase; C2’H, p-coumaroyl CoA 2’-hydroxylase; U6DT, umbelliferone 6-dimethylallyltransferase; U8DT, umbelliferone 8-dimethylallyltransferase; B5ODT, bergaptol 5-O-dimethylallyltransferase; DC, demethylsuberosin cyclase; P5H, psoralen 5-hydroxylase; P8H, psoralen 8-hydroxylase*. (c) Co-expression network analysis in *A. keiskei*. Edges were depicted if Pearson’s correlation was ≥ 0.65. In this network, P8H represents *P8H (CYP71AZ)*, and *P8H (CYP83A)* did not show a correlation above the threshold with any other genes.

The contigs in *the de novo* assembled library were filtered based on the expression and co-expression levels with any of the three *PTs* (Fig. S8), highlighting 376 contigs. These contigs were functionally annotated based on the Enzyme Commission (EC) classification (McDonald et al., 2023) using Blast2GO 6.0.3 (Conesa et al., 2005). Forty-eight contigs were assigned to EC1 (oxidoreductases), 18 of which belong to cytochrome P450s (Table S2, Fig. S9). After excluding four sequences with open reading frames (ORFs) shorter than 400 amino acids and three TC-unrelated genes, 11 candidate genes were retained. P450 family IDs were assigned to the candidates according to the classification procedure (P450 nomenclature committee, https://drnelson.uthsc.edu/CytochromeP450.html), and these IDs were used hereafter (Table S3). These 11 candidates were classified into CYP71 (eight candidates), CYP76 (one candidate), CYP92 (one candidate), and CYP736 (one candidate) subfamilies. All of these enzymes are part of the CYP71 clan, which is known to be the main clan involved in the phenylpropanoid pathway (Hamberger et al., 2013; Hansen et al., 2021).

To investigate the relationships between the candidate genes and known TC biosynthetic genes, a co-expression network analysis was performed. All candidates formed a network with genes responsible for the formation of coumarin and TC skeletons (Tables S4–S6, Fig. 2c), including *C2’H*, *PTs*, and *P450s*. In contrast, genes in the upstream phenylpropanoid pathway, such as *PAL*, *C4H*, and *4CL*, formed another cluster. As this study focused on the skeletal formation and structural diversity of TCs, all candidates were considered strong targets for further functional analyses.

### Cloning and sequence verification of candidate cytochrome P450s

The full coding sequences (CDSs) of the 11 candidate genes were isolated from the petiole by RT-PCR, except for *CYP71D941* and *CYP71AT442*. As these two genes were not amplified by PCR, their CDSs were artificially synthesized. To verify the conservation of cytochrome P450 motifs, the predicted ORFs of the 11 candidate genes were translated and aligned with the amino acid sequence of CYP71AJ1 (Fig. S10). Motifs essential for cytochrome P450 function, *e.g.*, proline-rich region, C-terminal helix, I-helix motif, EXXXR motif, PERF motif, and heme-binding motif (Hasemann et al. 1995; Fang et al. 2024), were conserved in all candidates. In addition, six substrate recognition sites (SRSs) were confirmed in all 11 candidates (Gotoh 1992; Dueholm et al. 2015). We also isolated *cytochrome P450 reductase* from *A. keiskei* (*AkCPR*), which shows 73.2 % nucleotide identity with *AtCPR2* (Harada et al. 2011). Accession numbers for the *P450s* and *AkCPR* sequences analyzed in this study are provided in the Data Availability section.

### In vitro screening of candidate cytochrome P450s

Each candidate was individually introduced into budding yeast, together with *AkCPR*, and their proteins were produced by galactose induction. Microsomal fractions were prepared from the cells and used as crude enzymes for *in vitro* assays. When microsomes were incubated with columbianetin and other coumarin molecules selected based on endogenous metabolites in *A. keiskei* (Table S7) under NADPH-saturated conditions, enzymatic products were detected for five clones out of 11 candidates, *i. e.*, CYP71AJ62, CYP71AJ63, CYP92A470, CYP71AJ20, and CYP736A530, by UPLC-PDA analysis (Table S8).

The first clone, CYP71AJ62, exhibited hydroxylation activity toward columbianetin, and the products were assigned as angelicin and hydroxylated columbianetin based on their retention times matching those of the products generated by the known angelicin synthase CYP71AJ4 from parsnip (Fig. 3a). Although the products were identical between the two enzymes, CYP71AJ62 predominantly produced one product. To identify the chemical structure of the main product of CYP71AJ62, we prepared vaginidiol (9R-hydroxylated columbianetin) via stereoretentive hydrolysis of apterin (vaginidiol 1’-glucoside) using a commercially available β-glucosidase (Steck et al., 1974). From 10.47 mg of apterin, 4.16 mg of its aglycone was obtained as white powder, and two-dimensional NMR analysis confirmed the position of the hydroxy moiety and stereochemistry at C-9 (Table S9 and 10, Fig. S11–15). LC-MS/MS analysis using this chemical standard showed that the fragmentation patterns in tandem mass spectrometry spectra were consistent between the CYP71AJ62 product and vaginidiol, confirming their identity (Fig. 3b–f). We also confirmed that the by-product of the parsnip CYP71AJ4 was this molecule.

**Figure 3.**
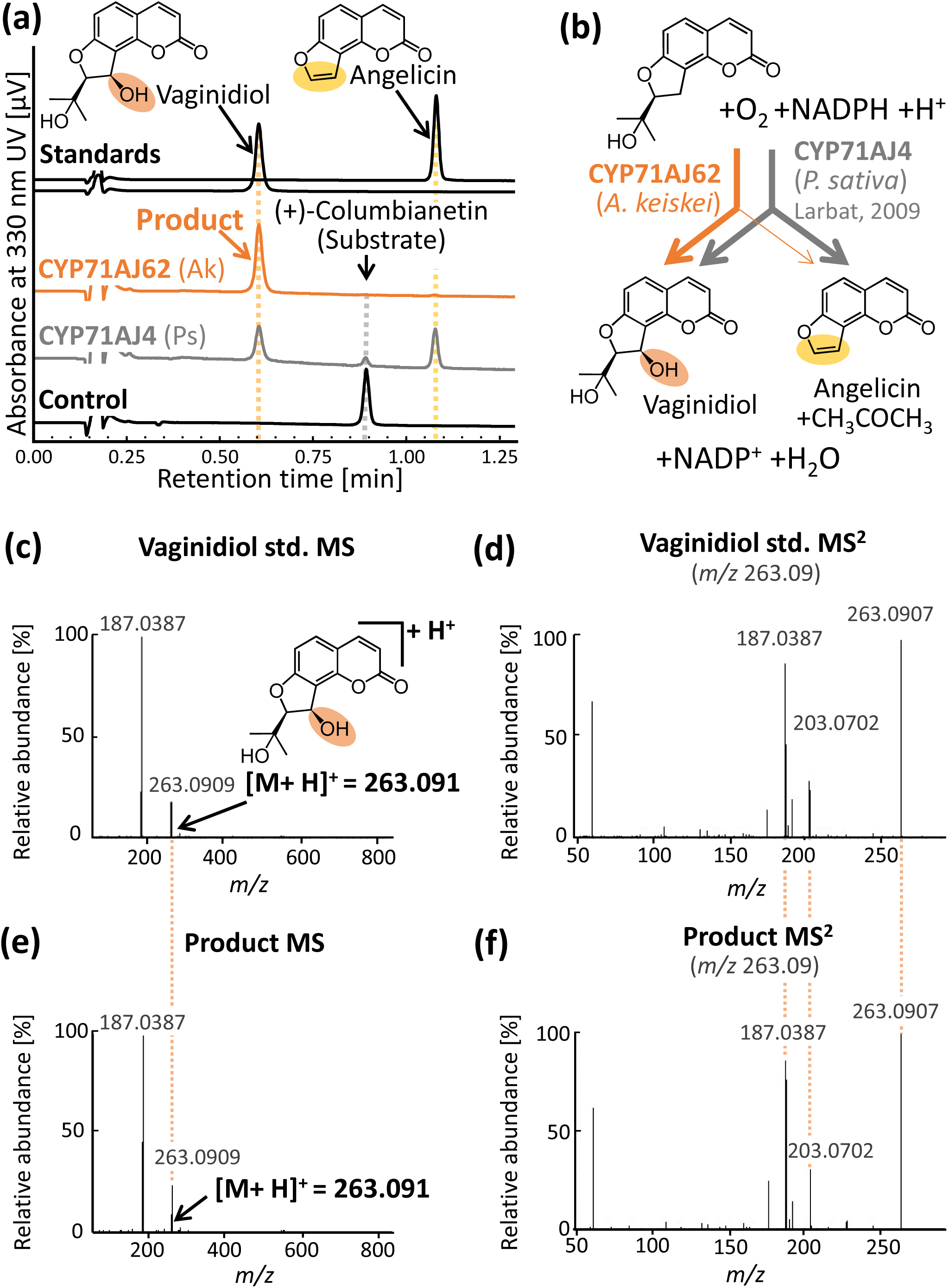
Vaginidiol synthase activity of CYP71AJ62. (a) UV chromatograms of the incubation mixture of CYP71AJ62 and CYP71AJ4 (Larbat et al., 2009), and negative control (AkCPR) using columbianetin as a substrate. Ak, *Angelica keiskei*; Ps, *Pastinaca sativa*. (b) Reactions catalyzed by CYP71AJ62 (orange) and CYP71AJ4 (gray). CYP71AJ4 produced both vaginidiol and angelicin, consistent with previous studies (Larbat et al., 2009), whereas CYP71AJ62 predominantly produced vaginidiol. (c, e) MS fragmentation of vaginidiol standard (c) and CYP71AJ62 product (e). (d, f) MS^2^ fragmentation of vaginidiol standard (d) and the product (f) from *m/z* 263.09.

The second clone, CYP71AJ63, epoxidized the isoimperatorin side chain, yielding oxypeucedanin (Fig. S16a, b). The CYP71AJ63 product was confirmed by comparison with authentic standards, which showed identical retention times in liquid chromatography and fragment patterns in tandem mass spectrometry (Fig. S16c–f). As these activities have not been reported in plants, we performed further biochemical analysis of CYP71AJ63 together with CYP71AJ62 in the following sections.

The third clone, CYP92A470, exhibited 5-hydroxylation activity for psoralen to produce bergaptol (Fig. S17). As this activity was minimal, we also attempted to characterize this enzyme in the *Nicotiana benthamiana* transient expression system (Fig. S18). When psoralen was infiltrated into *N. benthamiana* leaves expressing *CYP92A470*, the bergaptol content was increased in both expression lines. A similar result was obtained in the infiltration of *CYP71AZ20*, the homolog of the reported psoralen 5-hydroxylase (Ji et al., 2025). Despite this functional similarity, CYP92A470 exhibited only 35.1 % amino acid identity with CYP71AZ20. The remaining two clones were characterized as *A. keiskei* homologs of TC biosynthetic enzymes previously reported in other apiaceous species. Psoralen synthase activity was detected from *CYP71AJ20* (Fig. S19), with 89.2 % nucleotide homology with known *CYP71AJ1* from *Ammi majus* (Larbat et al. 2007). Similarly, *CYP736A530* exhibited columbianetin synthase activity (Fig. S20) and 95.7 % homology with *CYP736A366* (Zhao et al., 2024).

Based on the enzymatic activities, the five P450s were mapped onto the TC biosynthetic pathway, along with organ-specific metabolite accumulation and gene expression data (Fig. S21). Genes involved in the general phenylpropanoid pathway tended to be highly expressed in lateral roots, whereas those involved in TC biosynthesis showed higher expression in the petiole and main root cortex. Moreover, both the expression level of TC biosynthetic genes and the TC content were generally low in the root core. Among the identified P450s, CYP71AJ62 is positioned at the branch point between angular FC and DHFC biosynthesis, with type-B angular DHFCs preferentially accumulating over FCs in accordance with the product specificity of this enzyme. CYP71AJ63 is involved in the conversion of isoimperatorin to oxypeucedanin, which has not been reported in plants. Thus, we conducted further biochemical analysis of CYP71AJ62 and CYP71AJ63.

### In vitro enzymatic characterization of CYP71AJ62

To investigate the enzymatic properties of CYP71AJ62, we examined its substrate specificity, product specificity, and apparent *K*_m_ value. Five TCs were examined as substrates, and the product was compared with the parsnip enzyme CYP71AJ4 (Fig. 4a, b) under the optimal pH of CYP71AJ62 (Fig. S22). When CYP71AJ62 was incubated with lomatin, a structural analog of columbianetin, in which the dihydrofuran ring is replaced with a dihydropyran ring, an enzymatic product was observed (Fig. S23). The product was identified as khellactone, *i. e.*, 10-hydroxylomatin. In contrast, this enzyme showed no oxidative activity toward columbianadin (the 1’-angeloyl ester of columbianetin), angelicin (the core structure of angular FCs), or marmesin (a linear DHFC). This preference suggests that CYP71AJ62 accepts angular DHFC without ester modification. Among the substrates examined, the only difference between CYP71AJ62 and AJ4 was observed in the product profile from columbianetin.

**Figure 4.**
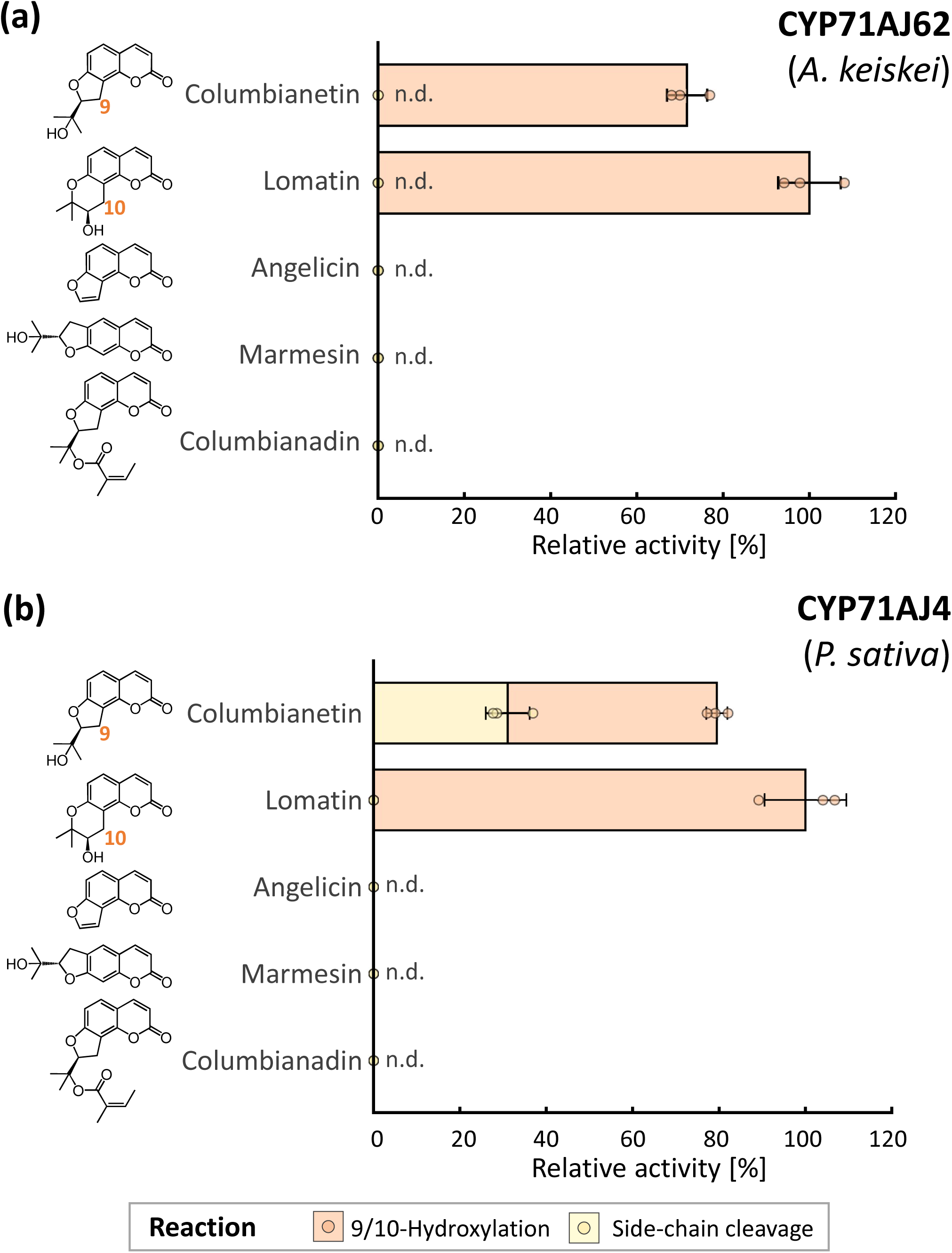
Substrate preferences of CYP71AJ62 and CYP71AJ4 Substrate preferences of CYP71AJ62 (a) and CYP71AJ4 (b) are presented. Hydroxylation at C9 in columbianetin or C10 in lomatin activities are shown in orange, and side-chain cleavage is shown in yellow. The relative activity of lomatin 10-hydroxylation was set to 100 %. Data are shown as mean ± standard deviation from three independent experiments. n.d., not detected.

We also compared the apparent *K*_m_ values between CYP71AJ62 and CYP71AJ4 for columbianetin in the formation of vaginidiol or angelicin (Table 1, Fig. S24 and 25). The apparent *K*_m_ value of CYP71AJ62 was approximately twice that of CYP71AJ4, indicating lower affinity for columbianetin in CYP71AJ62. The ratio of the apparent *K*_m_ values between angelicin and vaginidiol formation was nearly identical. In addition, CYP71AJ62 showed comparable apparent *K*_m_ values for columbianetin and lomatin (330 ± 40 nM), indicating that the type of heterocyclic moiety at the angular position has little influence on substrate affinity.

**Table 1.**
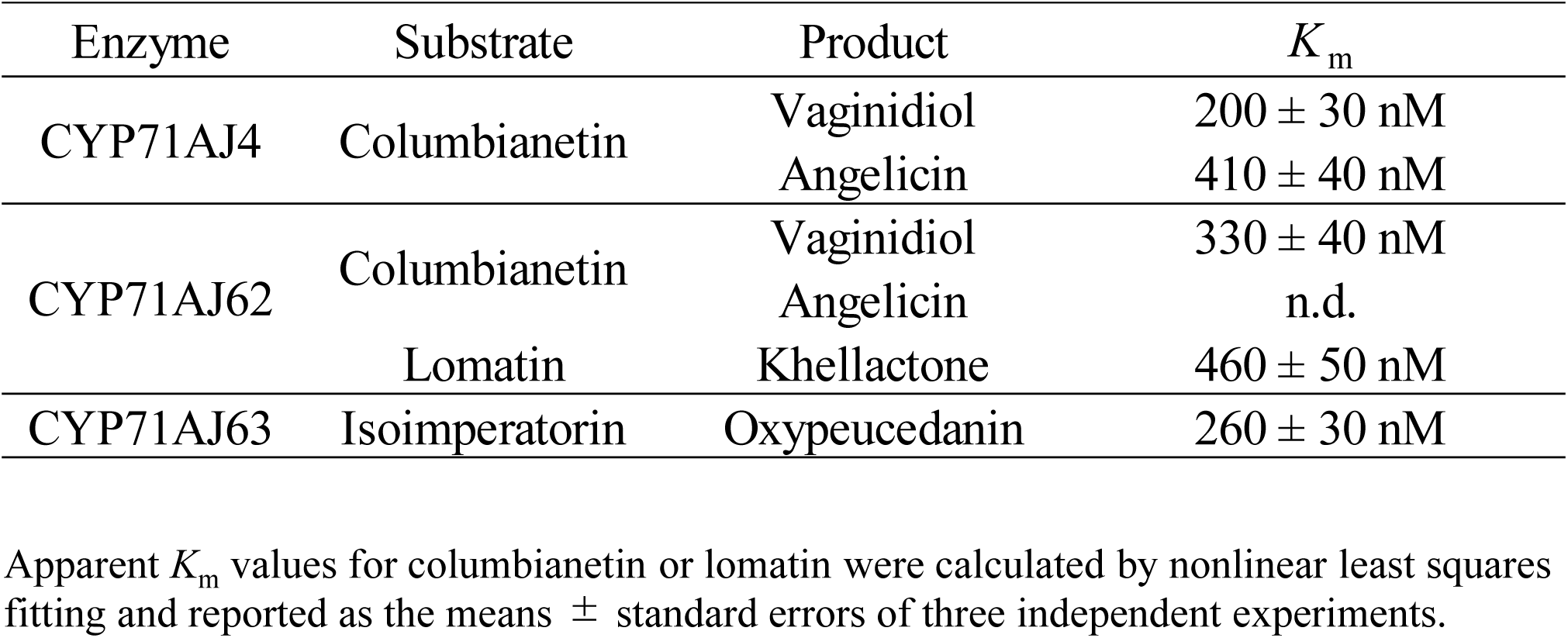
Apparent *K*_m_ values of CYP71AJ4, CYP71AJ62, and CYP71AJ63.

| Enzyme | Substrate | Product | $K_m$ |
| --- | --- | --- | --- |
| CYP71AJ4 | Columbianetin | Vaginidiol | $200 \pm 30$ nM |
| | | Angelicin | $410 \pm 40$ nM |
| CYP71AJ62 | Columbianetin | Vaginidiol | $330 \pm 40$ nM |
|  |  | Angelicin | n.d. |
| | Lomatin | Khellactone | $460 \pm 50$ nM |
| CYP71AJ63 | Isoimperatorin | Oxypeucedanin | $260 \pm 30$ nM |

Apparent $K_m$ values for columbianetin or lomatin were calculated by nonlinear least squares fitting and reported as the means ± standard errors of three independent experiments.

### In vitro enzymatic characterization of CYP71AJ63

The apparent *K*_m_ value for the conversion of isoimperatorin to oxypeucedanin by CYP71AJ63 was determined to be 260 ± 30 nM (Table 1, Fig. S26 and S27). The substrate specificity of CYP71AJ63 was examined using structural analogs of isoimperatorin (with 5-*O*-dimethylallyl moiety), *i. e.*, imperatorin (with 8-*O*-dimethylalllyl moiety), and bergamottin (with 5-*O*-geranyl moiety). These results indicate that this enzyme recognized only isoimperatorin as its substrate (Fig. S28), showing its high specificity toward the position and length of the prenyl side chain.

### Phylogenetic and syntenic analysis of the CYP71AJ subfamily underlying the diversification of enzymatic properties

To obtain evolutionary insights into the emergence of CYP71AJ62 and CYP71AJ63, we conducted a phylogenetic analysis of Apiaceae cytochrome P450s. The protein sequence dataset consisted of functionally characterized CYP71AJs from Apiaceae (Larbat et al., 2007; Larbat et al., 2009; Dueholm et al., 2015; Jian et al., 2020), and their 268 orthologous sequences (>400 amino acids) extracted from fourteen public genomes of Apiales using OrthoFinder (Emms et al., 2015). All functionally characterized CYP71AJs formed a clade with 33 sequences. These sequences and three outgroups from the orthogroups were used to generate the phylogeny shown in Fig. 5. The phylogenetic tree was subdivided into clades according to their TC substrates: marmesin, isoimperatorin, and columbianetin. The topology within the marmesin clade largely mirrors the taxonomic tree (Liao et al. 2013; Huang et al. 2024), suggesting an association with vertical inheritance during plant speciation (Han et al., 2025). CYP71AJ63 formed a clade with several sequences from other Apiaceae species, positioned close to the columbianetin clade. As these species have been reported to produce oxypeucedanin, this clustering may reflect conservation of the same function. CYP71AJ62 was located in the columbianetin clade, together with the angelicin synthase, CYP71AJ4 and AJ51. The positions of these enzymes suggest that angelicin synthase has evolved into vaginidiol synthase. In the recently released *A. keiskei* genome (Han et al., 2026), CYP71AJ62 showed the highest sequence similarity to the known angelicin synthase of parsnip, CYP71AJ4, and no other amino acid sequence corresponding to an angelicin synthase was identified.

**Figure 5.**
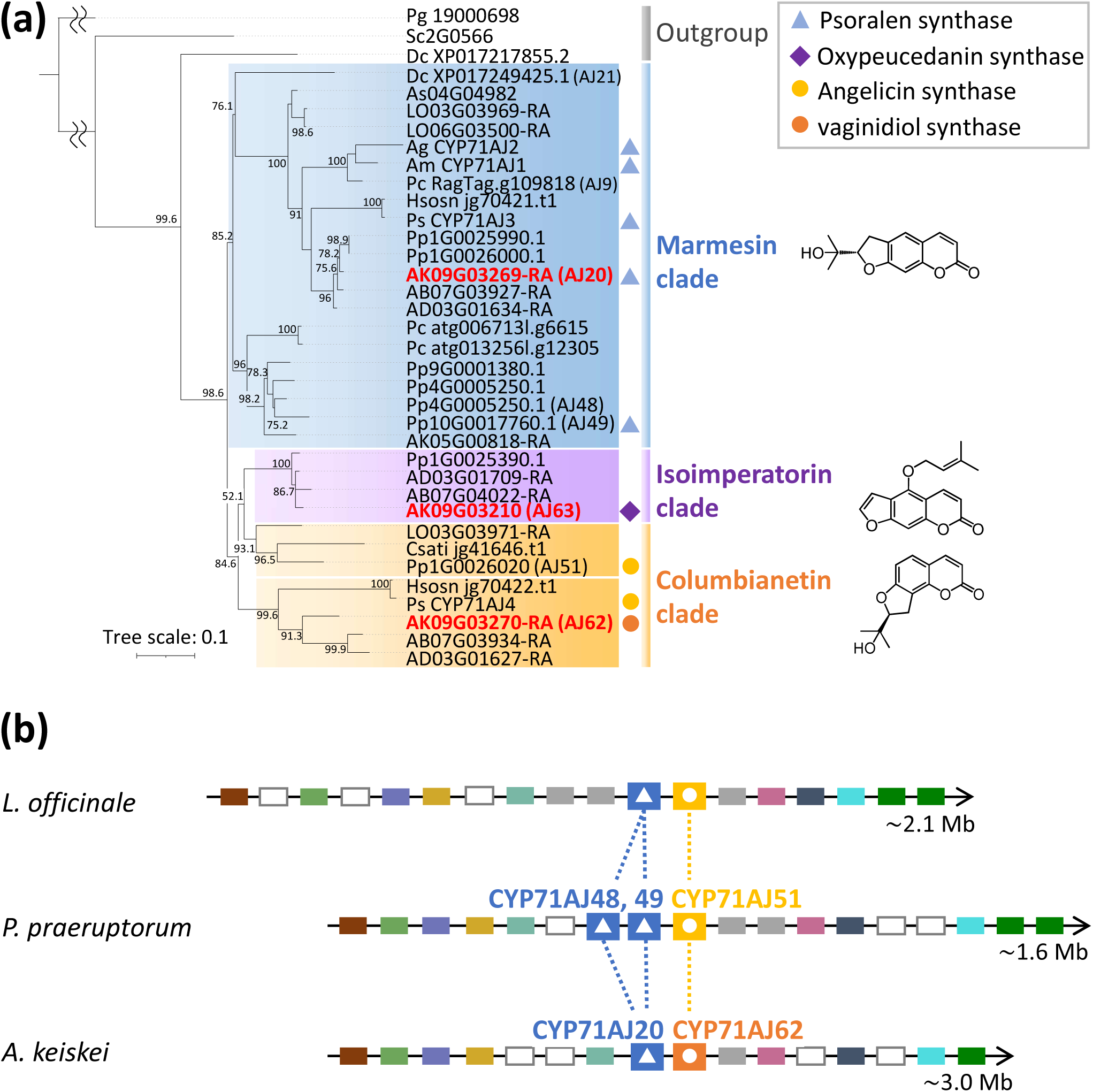
Phylogenetic relationships and genomic synteny of CYP71AJ enzymes (a) Phylogenetic tree of CYP71AJ polypeptide sequences obtained from the genomes of Apiales and functionally characterized CYP71AJs. The tree was constructed using 1,000 bootstrap replicates. Sequences were clustered into clades based on their substrates: marmesin, blue; isoimperatorin, purple; columbianetin, yellow. Symbols indicate functionally characterized enzymes: blue triangles represent psoralen synthase; a purple diamond represents oxypeucedanin synthase; yellow circles represent angelicin synthase; and orange circles represent vaginidiol synthase. AB, *Angelica biserrata*; AD, *Angelica dahurica*; Ag, *Apium graveolens*; Ak, *Angelica keiskei*; Am, *Ammi majus*; As, *Angelica sinensis*; Csati, *Coriandrum sativum*; Dc, *Daucus carota*; LO, *Levisticum officinale*; Pc, *Petroselinum crispum*; Pg, *Panax ginseng*; Pp, *Peucedanum praeruptorum*; Ps, *Pastinaca sativa*. (b) Syntenic region containing CYP71AJ62-orthologs in the genomes of *L. officinale*, *P. praeruptorum*, and *A. keiskei*. Colored boxes indicate syntenic genes, with the same colors representing putative orthologous genes among species. White boxes indicate genes not conserved among the three species. Blue boxes with triangles indicate orthologs of CYP71AJ20, CYP71AJ48, and CYP71AJ49, whereas yellow or orange boxes with circles indicate orthologs of CYP71AJ51 and CYP71AJ62.

To investigate the molecular evolutionary process in more detail, we examined the syntenic regions surrounding these genes using three Apiaceae genome assemblies with publicly available gene annotations: those of *Levisticum officinale*, *Peucedanum praeruptorum*, and *A. keiskei* (Fig. 6b; Table S12). Comparison of the genomic regions containing CYP71AJ62 orthologs in *L. officinale*, the most evolutionarily basal species in the columbianetin clade, *P. praeruptorum*, which harbors the functionally characterized angelicin synthase CYP71AJ51, and *A. keiskei* revealed conserved synteny in the surrounding regions. A blastn search of the region in *A. keiskei* detected no evidence of gene duplication, including pseudogene remnants. Conservation of the surrounding gene order indicated that angelicin synthase and CYP71AJ62 occupy orthologous genomic positions, strongly supporting the hypothesis that the functional transition occurred within a single-copy gene lineage rather than through gene duplication.

**Figure 6.**
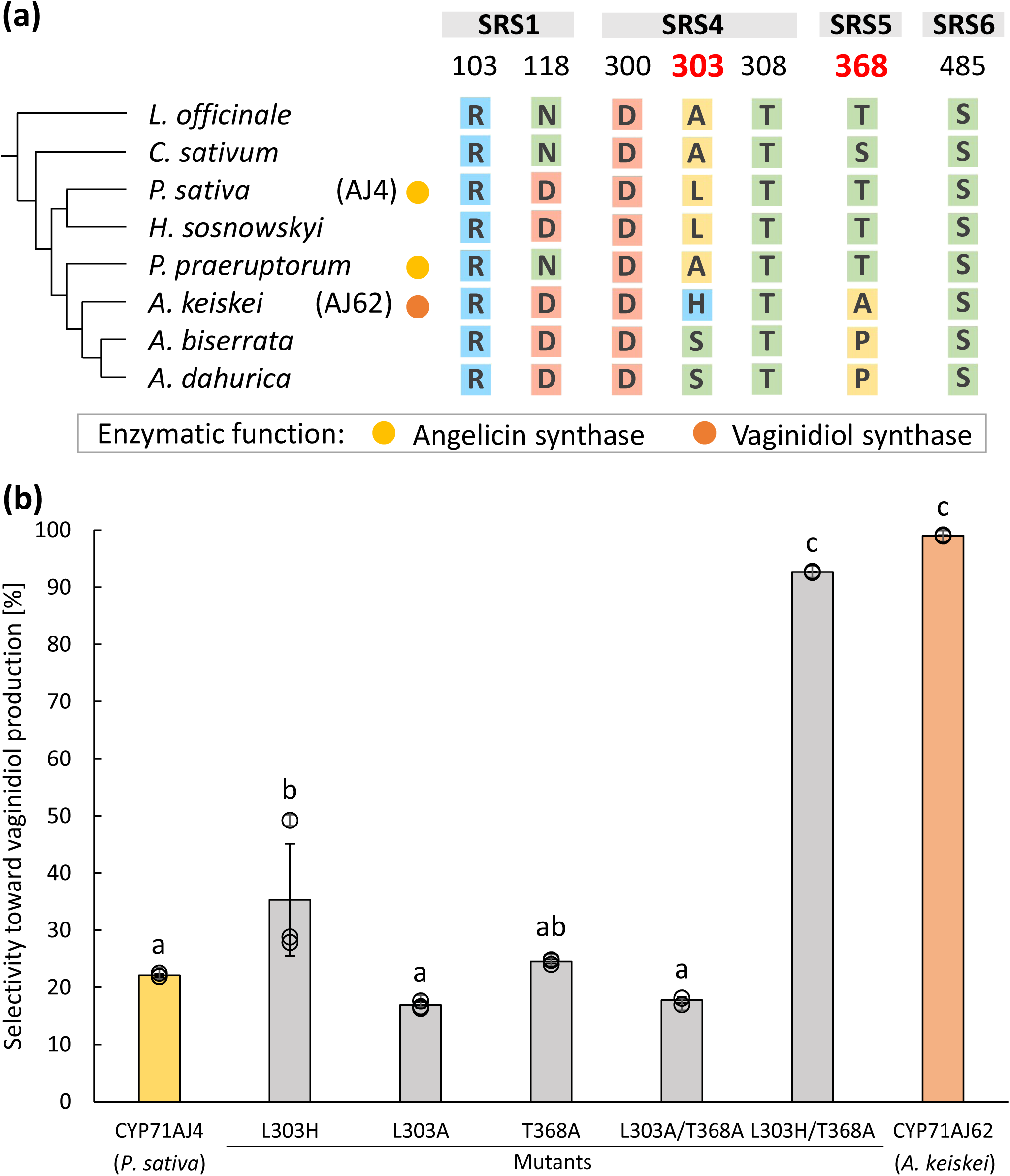
Identification of amino acid residues determining vaginidiol synthase activity (a) Comparison of the polar residues located within 5 Å of columbianetin across the columbianetin clade in Fig. 5a. The phylogeny was simplified from Fig. 5a. Yellow and orange circles indicate angelicin synthase and vaginidiol synthase activities, respectively. Residue numbering follows CYP71AJ4. Nonpolar amino acids, including proline, uncharged polar amino acids, acidic amino acids, and basic amino acids, are shown in yellow, green, salmon, and cyan, respectively. (b) Effect of site-directed mutagenesis on product specificity (*n* = 3). Data are shown as mean ± standard deviation from three independent experiments. Different letters indicate significant differences as assessed by Tukey’s HSD test (*p* < 0.05).

### In silico prediction of residues crucial for different columbianetin oxidation patterns between CYP71AJ4 and CYP71AJ62 active sites

To explore the factors underlying the differences in columbianetin oxidation activity between CYP71AJ4 and CYP71AJ62, we modeled the active-site docking of both CYP71AJ4 and CYP71AJ62 with the transmembrane domain removed (Fig. S29). Three-dimensional homology models of CYP71AJ4 and 62 without transmembrane regions were generated using ColabFold (Mirdita et al., 2022) to predict their putative structures (Fig. S29a). A columbianetin molecule was virtually docked into the heme-containing active sites of both enzymes using AutoDock Vina (Trott et al., 2010).

In the proposed angelicin formation mechanism, hydrogen atom abstraction was suggested to occur at C9 of columbianetin (Larbat et al., 2009). Based on this, we hypothesized that hydrogen atom abstraction at C9 is also the first step in vaginidiol formation (Fig. S30). We selected the model with the lowest binding affinity among those in which the distance between C9 and the heme iron-coordinated oxygen atom was within 5 Å (Fig. S29b, c). As polar residues may contribute to interactions between the enzyme and substrate, polar residues located within 5 Å of the substrate were examined, and seven residues were found for each enzyme. Among these residues, Leu303/His304 and Thr368/Ala369 differed between CYP71AJ4 and AJ62. Of the two residues, the nitrogen atom of His304 was located 2.7 Å from the hydroxyl oxygen on the isopropyl side chain of columbianetin, a distance compatible with hydrogen-bond formation.

### Site-directed mutagenesis of CYP71AJ4

The seven amino acid sites described above were compared among sequences belonging to the columbianetin clade (Fig. 5a, Fig. 6a, Fig. S31). All of these residues are located in SRS1, 4, 5, and 6, which supports previous studies showing that SRSs contribute to substrate specificity (Gotoh 1992; Dueholm et al. 2015). Positions 303 and 368, which were highlighted by three-dimensional modeling, differed in physicochemical properties between angelicin synthases (CYP71AJ51 and AJ4) and vaginidiol synthase (AJ62). Based on these differences, we designed site-directed mutagenesis in CYP71AJ4 following the substitution patterns observed along species divergence leading to *A. keiskei* (Fig. 6b). In these experiments, L303 of CYP71AJ4 was substituted with histidine (L303H), the corresponding residue in AJ62, or alanine, a small nonpolar amino acid (L303A). None of the single mutants reproduced the product specificity of CYP71AJ62. Similarly, the double alanine mutant (L303A/T368A) showed a product profile comparable to that of wild-type CYP71AJ4. Only the double mutant carrying the CYP71AJ62-type substitutions (L303H/T368A) exhibited a level comparable to that of wild-type CYP71AJ62. These results indicate that substitutions at both L303H and T368A are required for the functional conversion from angelicin synthase activity to vaginidiol synthase activity.

## Discussion

The cytochrome P450 superfamily comprises a large gene group that accounts for approximately 1 % of plant genomes. CYP71s form the largest family within this group and catalyze diverse oxidative reactions involved in specialized metabolism (Mizutani et al., 2010; Nelson et al., 2011). Within this CYP71 family, the CYP71AJ subfamily is involved in FC skeletal formation. CYP71AJ1 was the first enzyme in this subfamily identified as a psoralen synthase that catalyzes the final step of linear FC skeletal formation (Larbat et al., 2007). This reaction represents an unusual oxidative transformation involving side-chain C–C bond cleavage in a DHFC precursor and concomitant desaturation to form a furan ring. Subsequent characterization of CYP71AJ4 as an angelicin synthase demonstrated that the same P450 subfamily also participates in angular FC biosynthesis through a related oxidative cleavage reaction (Larbat et al., 2009). CYP71AJ enzymes involved in FC biosynthesis have been functionally characterized in diverse apiaceous species (Dueholm et al., 2015); however, the functions of CYP71AJ enzymes beyond FC biosynthesis remain unknown. In this study, we identified CYP71AJ62 in *A. keiskei* as a vaginidiol synthase involved in angular DHFC biosynthesis, expanding the known metabolic role of this subfamily to a distinct TC pathway.

The acquisition of novel functions in plant specialized metabolism is often facilitated by enzyme promiscuity, whereby a single enzyme can exhibit different activities. Promiscuous activities can provide a starting point for the evolution of new catalytic functions (Kreis et al., 2019). Gene duplication further promotes this process by relaxing functional constraints on one gene copy and allowing subsequent functional divergence (Maeda et al., 2021); this process also drives the specialization of P450 catalytic activities (Hansen et al., 2021; Werck-Reichhart et al., 2023). As an alternative route to neofunctionalization in plant genes, functional shifts can also occur without gene duplication through the evolutionary accumulation of sequence substitutions (Leong et al., 2017). For example, in tomato acylsugar biosynthesis, small numbers of amino acid substitutions in orthologous BAHD acyltransferases altered substrate preferences and contributed to changes in pathway reaction order (Fan et al., 2017). Our phylogenetic and syntenic results are consistent with this scenario for CYP71AJ62: functional specialization appears to have occurred in a single-copy lineage of *angelicin synthase* through only two amino acid substitutions. Although neofunctionalization is more constrained in single-copy genes (Panchy et al., 2016), changes in their catalytic specificity can alter pathway output because no duplicated copy retains the ancestral function. The functional shift of CYP71AJ62-lineage may consequently have redirected TC metabolic flux toward angular DHFCs at the expense of angular FCs in *Angelica* species including *A. keiskei*.

From an enzymological perspective, we found that two residues in SRS4 and SRS5 are sufficient to convert angelicin synthase to vaginidiol synthase. Among the six SRSs of cytochrome P450s, SRS1, 4, 5, and 6 play central roles in determining substrate-binding properties (Gotoh et al., 1992). SRS4 is located near the I-helix above the catalytic center, where it can directly interact with the substrate. SRS5 lies at the N-terminal region of the β1–4 strand facing the substrate-binding pocket and helps define the shape of the pocket. The residues in both regions contribute to substrate orientation within the active site (Schuler et al., 2006; Dueholm et al., 2015). Across plants, mammals, and bacteria, residues in SRS4 or SRS5 have been shown to influence product specificity, either individually or in combination with residues in SRS1. For example, single-residue substitutions in SRS1, SRS4, or SRS5 can alter product regioselectivity, whereas combined substitutions across all three sites have been shown to confer an additional product (Yoshimoto et al., 2014; Kardam et al., 2022; Subedi et al., 2022; Williams et al., 2022). This study provides an additional example of how amino acid residues in SRS4 and SRS5 can cooperate to determine product specificity. Further structural analysis will elucidate how the 303rd residue in SRS4 and the 368th residue in SRS5 cooperatively regulate the product specificity underlying angelicin and vaginidiol synthase activities.

The molecular evolution observed in this study may affect the chemical defense strategy of Apiaceae plants. Angular FCs are toxic to Apiaceae-specialist herbivores (Berenbaum et al., 1981), whereas some DHFCs show higher antioxidant activity (Dincel et al., 2013) and broader and stronger fungicidal activity against several pathogens (Song et al., 2017) than angular FCs. These differences in biological activity suggest that P450-mediated hydroxylation, such as CYP71AJ62-catalyzed vaginidiol formation, may help shape the balance among defense metabolites with different biological functions. In a recent report on soybean defense against the oomycete pathogen, the P450 GmIF6H1 regulates the metabolic flux between constitutive and infection-induced isoflavone pathways (Yang et al. 2025). Such P450-mediated flux regulation is also being explored for application as a breeding tool. In potato, knockdown of *CYP88B1* redirected metabolism toward resistance-related metabolites while reducing the accumulation of steroidal glycoalkaloids toxic to humans (Yasumoto et al., 2024). In the pharmacological context of the TCs, some angular DHFCs exhibit beneficial effects such as antitumor activity (Okuyama et al., 1991) and calcium-antagonistic activity with potential antihypertensive effects (Härmälä et al., 1992), while angular FCs can cause phototoxicity and interfere with xenobiotic metabolism in humans (Pohl et al., 1980; Mahendra et al., 2024). Our findings on the branching point between angular FCs and DHFCs would lead to metabolic engineering that improves stress tolerance and redesigns pharmacological activity with less toxicity to humans in commercially important Apiaceae species.

This study clarified important oxidation steps in TC biosynthesis using an integrated transcriptomic and biochemical approach, which is a powerful framework for understanding the molecular basis of plant specialized metabolism (Nett et al., 2020; Nakanishi et al., 2024). As *A. keiskei* produces a wide array of bioactive metabolites, including TCs, prenylchalcones, and diacetylenes (Akihisa et al., 2006; Kil et al., 2017), further research utilizing co-expression gene libraries may help elucidate the biosynthetic capacity of these metabolites and the underlying transcriptional regulatory network (Eshaghi et al., 2024). Given that lipophilic metabolites accumulate in the oil duct, an extracellular compartment, of apiaceous plants (Bosabalidis, 1996), such datasets will be useful to elucidate transcription factors involved in oil duct formation (Wang et al., 2024; Zhang et al., 2025), as well as transporters mediating export to the extracellular space (Nogia et al., 2021) in the future studies.

In conclusion, this study revealed the diverse cytochrome P450 functions underlying TC biosynthesis of Apiaceae, especially the metabolic divergence from angular FCs to DHFCs. These findings will lead to further elucidate the physiological functions of TCs, facilitate the breeding of Apiaceae species, and promote the development of useful bioactive agents for human health.

## Materials and methods

### Plant materials and chemical reagents

Petiole, main root cortex, stele, and lateral root of *A. keiskei* plants (Oshima variety) were maintained under the same conditions as previously reported (Munakata et al. 2021). Coumarin standards were purchased from suppliers or kindly offered as shown in Table S13, except for vaginidiol. To prepare vaginidiol, apterin (10.47 mg, Taniguchi et al., 2005) was dissolved in 22.5 mL of 50 mM citrate-phosphate buffer (pH 5.0) containing 10 % (v/v) glycerol to a final concentration of approximately 1.0 mM. β-Glucosidase from almonds (Sigma-Aldrich, St. Louis, MO, USA) was dissolved at 50 mg mL^-1^ (≥ 200 U/mL). Glucosidase solution (2.5 mL) was added at 0, 24, and 48 h into the incubation system. The reaction mixture was incubated at 50 °C without agitation (Patchett et al. 1987; Bi et al. 2019). After 72 h, an equal volume of chloroform was added, and the mixture was vortexed for 10 s and then centrifuged (5,000 × *g*, 5 min). Chloroform extraction was repeated twice. The organic phase was collected and dried over anhydrous sodium sulfate. Evaporation and drying of the organic layer yielded 4.16 mg. Approximately 1 mg of the dried residue was dissolved in ethanol and subjected to LC-MS/MS analysis. The remaining material was dissolved in approximately 0.5 mL of CDCl₃, and the ¹H NMR spectra were recorded at 600 MHz and ¹³C spectra at 150 MHz using an Agilent NMR system 600-DD2 (Agilent Technology, Santa Clara, CA, USA). Chemical shifts were referenced to tetramethylsilane (δ = 0 ppm). In addition, 2D NMR experiments, including HSQC, HMBC, and NOESY, were performed essentially as described Taniguchi et al., 2005. Data acquisition and processing were accomplished using VnmrJ 3.2 (Agilent).

### LC-MS/MS for accurate mass analysis of plant samples and enzymatic products

The sample solutions (5 μL) were analyzed using a Vanquish system (Thermo Fisher Scientific, Waltham, MA, USA) coupled to a Q Exactive Focus hybrid quadrupole-orbitrap mass spectrometer with a heated electrospray ionization (HESI II) source (Thermo Fisher Scientific). The analytical column was a C18 reverse-phase column (ACQUITY UPLC BEH C_18_ Column, 130 Å, 1.7μm, 2.1 mm × 150 mm; Waters, Milford, MA, USA) with a UPLC BEH C18 VanGuard pre-column (1.7 μm, 2.1 × 5 mm), and the column oven was set at 40 °C. The compounds were separated at a flow rate of 0.3 mL min^-1^, consisting of 0.1 % (v/v) formic acid as solvent A and acetonitrile containing 0.1 % (v/v) formic acid as solvent B. For plant metabolites, a linear gradient comprised of 5–95 % B for 15 min, followed by a 5 min wash with 95 % B. For enzymatic products, a linear gradient comprised of 5–95 % B for 15 min, followed by a 5 min wash with 95 % B. The separated compounds were detected using a mass spectrometer operated in full-scan mode over an *m/z* range of 66.7–1,000 at a resolution of 70,000, followed by higher-energy collisional dissociation (HCD) with a stepped normalized collision energy of 20, 30, 50 %. The electrospray ionization voltage was set at 2.0 kV. The scan mode settings are listed in Table S14. Data were recorded and analyzed using the Xcalibur software 1.5 (Thermo Fisher Scientific) and visualized using the FreeStyle software 1.5 (Thermo). The coumarin contents of the peak areas were calculated using calibration curves constructed with authentic compounds.

### Construction of transcriptome datasets

Total RNA was purified from frozen tissues using the RNeasy Plant Mini Kit (Qiagen, Hilden, Germany) and treated with DNase (Ambion DNA-free DNase Treatment and Removal, Thermo Fisher Scientific). The library was paired-end sequenced using the Illumina NovaSeq X Plus system (Illumina, San Diego, CA, USA). The acquired sequences were assembled using rnaSPAdes v3.13.0 after trimming with fastp 0.23.4, and transcript expression was quantified as TPM values with Salmon v1.5.0.

### Transcriptomic analysis of A. keiskei

Co-expression analysis was performed based on the Pearson correlation coefficient with three bait *PT*s. Contigs corresponding to *U6DT* and *U8DT* were annotated based on blastn scores with reported counterparts from *P. sativa* (Munakata et al. 2016), and *B5ODT* was already described in *A. keiskei* (Munakata et al. 2021). Contigs correlated with any of the query *PT*s with a coefficient value of 0.65 or more were functionally annotated using Blast2GO 6.0.3 (Conesa et al. 2005). Contigs annotated as *cytochrome P450* with either unknown functions or known involvement in TC biosynthesis were subjected to functional analysis. Contigs lacking ORFs that were longer than 1,000 bp were excluded. Multiple alignment by MAFFT v7.490 and its visualization were performed in Geneious Prime 2024.2.1 (Dotmatics, Boston, MA, USA).

### Isolation of P450 genes and construction of yeast expression plasmids

Total RNA prepared for transcriptome library construction served as the template. Reverse transcription to cDNA was performed from total RNA extracted from the petiole using SuperScript IV Reverse Transcriptase (Thermo Fisher Scientific). Synthesized cDNA was diluted and used as a template for PCR reactions. The nucleotide sequences containing the full CDSs of candidate genes were amplified by PCR using KOD One PCR Master Mix (Toyobo, Osaka, Japan) and primer pairs (Table S15). The amplicons were inserted into the pGEM T-easy vector system (Promega, Madison, WI, USA) for sequencing. The CDSs of *CYP71D941* and *CYP71AT442* were artificially synthesized and codon-optimized for *Saccharomyces cerevisiae* by GeneArt (Thermo Fisher Scientific). The CDSs of candidate genes and *AkCPR* were amplified by PCR using PrimeSTAR Max DNA Polymerase (Takara, Kusatsu, Japan). The PCR product of *AkCPR* was introduced into the NotI/SacI site of the pESC-Ura vector (Agilent), and each candidate gene was subsequently introduced into the BamHI/SalI site of pESC-Ura-*AkCPR* by In-fusion Snap Assembly cloning (Takara).

### Heterologous expression of candidate cytochrome P450s in budding yeast and microsome preparation

Recombinant pESC-Ura-*AkCPR* plasmids containing candidate *P450s* were introduced into the *S. cerevisiae* strain INVSc1 (Thermo Fisher Scientific) using the lithium acetate method (Ito et al. 1983). A single colony of *S. cerevisiae* harboring the pESC-Ura construct was precultured in 3 mL of synthetic dextrose (SD) medium lacking uracil and containing 2 % glucose overnight at 30 °C with shaking at 200 rpm. A 1 mL aliquot of the preculture was inoculated into 100 mL of the same medium and further cultured overnight. The cells were collected and resuspended in 100 mL of SD medium lacking uracil and containing 2 % galactose. Protein expression was induced at 30 °C for 24 h with shaking at 200 rpm. Unless otherwise stated, subsequent procedures were performed at 4 °C or on ice. The induced cells were incubated in 100 mM Tris-HCl (pH 9.6) containing 10 mM dithiothreitol (DTT) for 30 min at 30 °C, washed with sorbitol-KPi buffer (20 mM potassium phosphate (pH 7.5), 1.2 M sorbitol), and treated with Zymolyase-20T (Nacalai Tesque, Kyoto, Japan) at 5 mg g⁻¹ wet cells for 1 h at 30 °C. The resulting spheroplasts were suspended in breaking buffer (20 mM Tris-HCl (pH 7.4), 0.6 M sorbitol, 1 mM DTT, and 100 μM phenylmethylsulfonyl fluoride (PMSF)), homogenized, and centrifuged at 10,000 × *g* for 30 min. The supernatant was further ultracentrifuged at 100,000 × *g* for 30 min. The microsomal pellet was resuspended in reaction buffer (100 mM Tris-HCl (pH 7.5), 100 μM PMSF) for immediate enzyme assays or in TEG buffer (50 mM Tris-HCl, pH 7.5, 1 mM EDTA, and 30 % glycerol) for storage at -30 °C. Protein concentrations were determined using the Qubit Protein Assay Kit (Thermo Fisher Scientific).

### In vitro P450 screenings and assays for estimating the catalytic and kinetic features

Microsomal fractions were suspended in 0.1 M KPi buffer (pH 7.5) containing 100 μM of substrate and 200 μM NADPH in a final volume of 100 μL, except for the activity comparison between CYP71AJ4 and AJ62 (Fig. 3a), for which pH was set to 7.0 to reproduce the previous results (Larbat et al., 2009). For substrate specificity analysis and kinetic analysis, the microsomal fraction was suspended in Tris–EDTA–Glycerol buffer and stored at -30 °C. In the assays to examine substrate specificity, microsomal fractions were incubated in the presence of 1.0 μM substrate standards and 200 μM NADPH for 30 min. The kinetic parameters of CYP71AJ62 and CYP71AJ63 were determined by incubating the functional P450s with each substrate in 0.1 M KPi buffer, pH 7.5, containing 200 μM NADPH. The concentrations of substrates ranged from 62.5–2000 nM. Incubations were kept at 27 °C, under an agitation of 600 rpm, and stopped by adding 250 μL of ethyl acetate, shaken thoroughly, and centrifuged (4400 × *g*, room temparature, 5 min). A 200 μL aliquot of the ethyl acetate phase was collected and evaporated to dryness by vacuum centrifugation. The residue was dissolved in 25 μL of methanol, mixed with an equal volume of ultrapure water, and centrifuged. The resulting supernatant was subjected to UPLC-PDA analyses. For any combination that showed peak detection in the initial screening, three independent reactions were subsequently conducted twice.

### Detection and quantification of enzymatic reaction products

Reaction products were analyzed using UPLC-PDA, as described by Han et al. (2025). Samples exhibiting peaks suspected to correspond to the enzymatic product were further analyzed by LC-MS/MS using the same conditions as described above. To estimate the structures of the enzyme reaction products, retention time and tandem mass spectrometric fragmentation patterns were compared with those of standard molecules.

### Transient expression of recombinant protein in N. benthamiana for comparison of enzymatic activity

The CDSs of *AkCPR* and *CYP92A470* were subcloned into the SalI site of the pTKB3 vector (Nosaki et al. 2021). The plasmids were individually introduced into *Agrobacterium tumefacien*s GV3101. Cultivation and infiltration of *N. benthamiana* were performed as described by Koita et al. (2025). Three experimental groups with biological triplicates were prepared: pTKB3-*AkCPR* with pTKB3-*EGFP* as control, pTKB3-*AkCPR* with pTKB3-*CYP92A470* and pTKB3-*AkCPR* with pTKB3-*CYP71AZ18-like*. Three days after agroinfiltration, 100 μM psoralen in 0.1 % DMSO was fed using a 1-mL syringe (Terumo, Tokyo, Japan) and incubated for 24 h in a growth room. The treated leaves were frozen in liquid nitrogen and immediately crushed. 100 mg of fresh leaf powder was extracted with 500 μL of ethyl acetate. The supernatant was evaporated and dissolved in 200 μL of 50 % methanol and analyzed using LC-MS/MS.

### Phylogenetic analysis of the CYP71AJ subfamily in Apiaceae

The protein sequence dataset consisted of known CYP71AJs from Apiaceae (Larbat et al., 2007; Larbat et al., 2009; Dueholm et al., 2015; Jian et al., 2020), and genome sequences obtained from public databases (Table S11). All sequences were classified using OrthoFinder 3.0.1b1 (Emms et al., 2015). Sequences longer than 400 amino acids from the orthogroup were used for phylogenetic analysis using iQTree 2.4.0 (Minh et al., 2020) with Q.plant +I+G4 substitution models. Model selection was performed using Model Finder Plus, with bootstrap with 1,000 replicates and 1,000 Shimodaira-Hasegawa approximate Likelihood Ratio test (SH-aLRT) replicates. From the phylogenetic tree, 33 sequences that formed the cluster, including all known CYP71AJs, and three outgroup sequences were extracted from the same orthogroups. The 36 sequences were aligned using MAFFT v7.525 (Katoh et al., 2013) and used for phylogenetic tree construction using iQTree with Q.plant + G4 substitution models. The resulting trees were visualized using iTOL v7 (Letunic et al., 2024).

### Synteny analysis of CYP71AJ62

Three plant species with sufficiently annotated genomes to provide assigned gene IDs were selected for the analysis. The genome data of the three plant species were obtained from the public (Huang et al., 2024; Han et al., 2026). Orthology IDs were annotated using Hayai-Annotation Plants v2.0 (Ghelfi et al., 2024). Genome synteny was detected using orthology IDs located in the genomic region around CYP71AJ62 orthologs.

### Homology modeling and docking experiments

Three-dimensional homology models of CYP71s were generated using ColabFold v1.5.5 (Mirdita et al., 2022). Three-dimensional models of heme (CID: 4971) and columbianetin (CID: 92201) were downloaded from PubChem (https://pubchem.ncbi.nlm.nih.gov/). The PDB files were analyzed using the PyMOL Molecular Graphics System v.3.1.0 (Schrödinger, LLC), and the first 30 residues of the enzyme proteins were excluded as the N-terminal transmembrane domain. After hydrogen atoms were added using AutoDock Tools 4.1 (Morris et al., 2009), heme was docked to the heme-binding site as a rigid ligand with a 16 Å × 16 Å × 16 Å grid box using AutoDock VINA v1.2.7 (Trott et al., 2010). Unless otherwise specified, standard settings were used. Columbianetin was then docked into the CYP71AJ models containing docked heme. For substrate docking, the search space was defined as a 16 Å × 16 Å × 16 Å cube centered on heme iron. The resulting 39 (AJ4) or 31 (AJ62) binding poses were ranked according to their predicted binding affinities. Among the poses in which the distance between C9 of columbianetin and the modeled Compound I oxygen was within 5 Å, the pose with the lowest predicted binding affinity was selected for further analysis. PyMOL was subsequently used to visualize and compare the selected models, measure interatomic distances, merge the receptor and ligand structures, and prepare the figures.

### Site-directed mutagenesis of the CYP71AJ4 enzyme

The CDSs of CYP71AJ4 with the mutations L303A, T368A, and L303A/T368A were artificially synthesized and codon-optimized for Saccharomyces cerevisiae by GeneArt (Thermo Fisher Scientific). All enzymes with the L303H point mutation were constructed by mutagenic PCR using the primers and PrimeSTAR Max DNA Polymerase (Takara). The primers are listed in Table S15. The pMA-RQ plasmid containing the artificially synthesized *P450* was used as the reaction substrate. After constructing the point mutation *P450s* on the pMA-RQ plasmid, we introduced them into the pESC-Ura-*AkCPR* vector via gap-repair cloning (Eckert-Boule et al., 2012). In short, the CDS was amplified using the primer set with the tag overlapping the sequence of the multiple cloning site 2 (MCS2) of pESC-Ura. The linearized pESC-Ura-*AkCPR* plasmid and amplified insert were transformed in the same way as the functional expression of the candidate cytochrome P450s in budding yeast cells. Colony PCR was performed using KOD FX Neo (Toyobo) and primers designed to flank the MCS2 site of pESC-Ura. The resulting PCR products were subjected to Sanger sequencing to confirm the insertion of the target sequence into the MCS2 site.

### Statistics and reproducibility

Enzymatic reactions were carried out independently in triplicate. Apparent *K*_m_ values were calculated by nonlinear least squares fitting in SigmaPlot 14.5 (Systat Software Inc., CA, USA). The psoralen 5-hydroxylation activity in *N. benthamiana* was examined using Welch’s *t*-test in Microsoft Excel 2019, as the homogeneity of variances could not be assumed. Vaginidiol production selectivity in site-directed mutant enzymes was analyzed using Tukey’s HSD test in the multcomp package on R (v4.2.3) (Hothorn et al., 2008).

## Acknowledgements

The authors thank Dr. Alain Hehn and Dr. Frédéric Bourgaud (Lorraine University–INRAe) for kindly providing the pCR8-CYP71AJ4 (Larbat et al., 2009) and chemical standards. We thank Dr. David Nelson (University of Tennessee) for naming the P450 genes isolated from A. *keiskei*. We are grateful to Dr. Takao Yamaura (Nippon Shinyaku Co., Ltd.) for providing plant materials. Dr. Katsuhiko Minoura (Osaka Medical and Pharmaceutical University) is gratefully acknowledged for his expertise in NMR measurements. Technical assistance from Ms. Kaori Kanazawa is sincerely appreciated. Part of this study was conducted using the facilities of DASH/FBAS (Research Institute for Sustainable Humanosphere (RISH), Kyoto University). This work was supported in part by the Japan Society for the Promotion of Science (JSPS), Grants-in-Aid for Scientific Research (KAKENHI) (grant no. 25K09039 to R.M.) and Transformative Research Areas (A) (grant no. 23H04967 to K.Y. and R.M.); by JSPS Overseas Research Fellowships (to R.M.); by the Precursory Research for Embryonic Science and Technology (PRESTO) program of the Japan Science and Technology Agency (grant no. JPMJPR20D7 to R.M.); and by the GteX Program (JPMJGX23B2 to A.S.). Additional support was provided by the RIKEN Cluster for Science, Technology, and Innovation Hub (to K.Y.), RISH, Kyoto University (Mission Research 5), and the Iwatani Research Scholarship for Graduate Students (to N.S.) from the Iwatani Naoji Foundation.

## Author contributions

N.S. and R.M. conceived the project. N.S., J.H., and R.M. performed the transcriptome analysis. N.S., M.T., and R.M. performed the metabolic analysis. N.S., Y.K., K.M., and R.M. performed the enzymatic analysis. A.S., K.Y., and R.M. supervised the study. N.S. and R.M. wrote the manuscript with the contributions of the co-authors.

## Data Availability Statement

Nucleotide sequences of *cytochrome P450s* and *AkCPR* are available in NCBI (Table S19). The datasets are publicly available in the DNA Data Bank of Japan (DDBJ) under BioProject accession PRJDB35810. The associated accession numbers include BioSample SAMD01605014-SAMD01605037, Experiment DRX730296-DRX730319, and Run DRR750693-DRR750716.

## Supporting information

Additional Supporting Information may be found in the online version of this article.

**Table S1.** Summary statistics of the *de novo* assembly of *Angelica keiskei* transcriptome data.

**Table S2.** Pearson correlation and gene annotations of EC1-annotated contigs correlated with bait PTs. The largest correlation coefficient with *U6DT*, *U8DT*, or *B5ODT* is highlighted in red in this table and shown in Fig. 2b. These results were used for screening candidates (Table S8). Candidates were defined as sequences that met the ORF criteria, excluding those with previously characterized biosynthetic functions unrelated to coumarins.

**Table S3.** Correspondence between contigs and CYP IDs.

**Table S4.** Co-expressed biosynthetic genes in *A. keiskei.* The genes are shown in Figure 3b.

**Table S5.** Data table for co-expression network analysis. The Pearson correlation coefficient (*r*) was calculated for each pair of genes. To avoid redundancy, gene pairs were listed only once, with one gene assigned as the node and the other as the source.

**Table S6.** Organ-specific expression of TC biosynthetic genes based on TPM values.

**Table S7.** Substrates used for the screening.

**Table S8.** Functional screening of eleven candidates. Each trial was performed once, and in the trials marked with “+”, additional peaks, except for the substrate, were detected. DMS, demethylsuberosin; Ost, osthenol, Mar, marmesin; Cnt, columbianetin; Cnd, columbianadin; Pso, psoralen; Ber, bergapten; Isoimp, isoimperatorin.

**Table S9.** ^1^H-NMR spectrum of prepared vaginidiol

**Table S10.** ^13^C-NMR spectrum of prepared vaginidiol.

**Table S11.** Genome datasets for phylogenetic analysis.

**Table S12.** Synteny analysis of CYP71AJ62 orthologues.

**Table S13.** List of chemical standards of metabolites

**Table S14.** Metabolites detection by selected ion recording mode.

**Table S15.** List of primers used in this study

**Table S16.** The accession numbers of *cytochrome P450s* and *AkCPR* are available on DDBJ.

**Figure S1.** The core structures of tricyclic coumarins in this study.

**Figure S2.** Overview of the modification pathways of linear furanocoumarins in angiosperms.

All biosynthetic enzymes of linear FCs described are shown in solid arrows (Hehmann et al., 2004; Kruse et al., 2008; Limones-Mendez et al., 2020; Zhang et al., 2022; Bouillé et al., 2025; Ji et al., 2025). Cytochrome P450s are labelled with their IDs and color-coded according to the plant family. B8H, bergapten 8-hydroxylase; B5ODT, bergaptol 5-O-dimethyallyltransferase; B5OMT, bergaptol 5-O-methyltransferase; 5HB8OMT, 5-hydroxybergapten 8-O-methyltransferase; 8HX5OMT, 8-hydroxyxanthotoxin 5-O-methyltransferase; IE, isoimperatorin epoxidase; OH, oxypeucedanin hydrolase; P5H, psoralen 5-hydroxylase; P8H, psoralen 8-hydroxylase; X5H, xanthotoxin 5-hydroxylase; X8ODT, xanthotoxol 8-O-dimethyallyltransferase; X8OMT, xanthotoxol 8-O-methyltransferase.

**Figure S3.** The petiole, main root cortex, main root stele, and lateral root of *A. keiskei* used for analysis.

**Figure S4.** Quantified coumarin molecules in *Angelica keiskei* organs. The two types of dihydrofuranocoumarins (DHFCs), type-A and type-B, are shown as (A) and (B). In SciFinder registration, a ^†^ indicates the metabolites that have not been reported in *Angelica keiskei*, and ^‡^ indicates the metabolite that has not been reported in the genus *Angelica*.

**Figure S5.** Tissue-specific accumulation of coumarins.

Data represent the contents of individual metabolites in simple coumarins (a), linear dihydrofuranocoumarin (b), linear furanocoumarins (c), angular dihydrofuranocoumarins type A (d), type B (e), and angular furanocoumarins (f) in the petiole (P), main root cortex (MRC), main root stele (MRS), and lateral root (LR). The total content of each coumarin group (a–f) is also shown. Data are presented as mean ± standard error from six individuals. The LC-MS detection parameters for the analyzed metabolites are listed in Table S14.

**Figure S6.** Coumarin accumulation patterns in *A. keiskei* organs.

TC accumulation in the petiole, main root cortex, main root stele, and lateral root. The accumulation amounts of simple coumarins, linear furanocoumarins, angular dihydrofuranocoumarins type A, type B, and angular furanocoumarins are shown in grey, blue, brown, orange, and yellow, respectively. Data are shown as mean ± standard error from six individuals. The LC-MS detection parameters for the metabolites analyzed are listed in Table S14.

**Figure S7.** Histogram of contigs by sequence length.

**Figure S8**. Stepwise filtering of candidate cytochrome P450 genes.

**Figure S9.** Functional annotations of contigs co-expressed with TC-biosynthetic *PTs* using Blast2GO. Annotation of each contig was performed using Blast2GO.

**Figure S10**. A multiple alignment of the polypeptide sequences of candidate P450s and CYP71AJ1.

Amino acid sequences deduced from the eleven candidates and CYP71AJ1 were aligned using MAFFT v7.490 and visualized using Geneious Prime 2024. SRSs and motifs were annotated according to Dueholm et al. (2015) and Fang et al. (2024). The transmembrane region was estimated using TargetP-2.0 (Armenteros et al., 2019).

**Figure S11.** ^1^H-NMR spectrum of vaginidiol obtained by hydrolysis.

**Figure S12**. ^13^C-NMR spectrum of vaginidiol obtained by hydrolysis.

**Figure S13.** HMBC correlations of vaginidiol obtained by hydrolysis. An unassigned methylene peak was detected in NMR spectra (indicated by a yellow arrow). This peak was regarded as an impurity due to no coupling with any proton or carbon of vaginidiol and no UV absorption.

**Figure S14**. HSQC correlation of vaginidiol obtained by hydrolysis.

**Figure S15**. NOESY correlations of vaginidiol obtained by hydrolysis. The strong correlation between H-8 and H-9 was observed (shown in red arrows on the structure and NOESY plane), which demonstrates the *cis*-configuration.

**Figure S16.** Oxypeucedanin synthase activity of CYP71AJ63.

(a) *In vitro* assay of CYP71AJ63 and control (AkCPR) using isoimperatorin as a substrate

(b, d) MS fragmentation of oxypeucedanin standard (b) and CYP71AJ63 product (d).

(c, e) MS^2^ fragmentation of oxypeucedanin standard (c) and the product (e) at *m/z* 287.09.

**Figure S17**. Psoralen 5-hydroxy activity of CYP92A470.

a. UV chromatogram of *in vitro* yeast microsome assay of CYP92A470 and control (AkCPR) using psoralen as a substrate.

(b, d) MS fragmentation of bergaptol standard (b) and CYP92A470 product (d).

(c, e) MS^2^ fragmentation of bergaptol standard (c) and the product (e) at *m/z* 203.03. The structure of the *m/z* 147 fragment was assigned based on Du et al. (2025).

**Figure S18.** Psoralen 5-hydroxy activity of CYP92A470 and AkCYP71AZ18-like in *Nicotiana benthamiana* leaves.

Bergaptol content of *N. benthamiana* leaves expressing EGFP (control), CYP92A470 (a), and AkCYP71AZ18-like (b). Data are shown as mean ± standard deviation with three biological replicates (*n* = 3). Statistical significance was evaluated using Welch’s *t* test.

**Figure S19**. Psoralen synthase activity of CYP71AJ20.

(a) UV chromatogram of *in vitro* yeast microsome assay of CYP71AJ20 and control using marmesin as the substrate.

(b, d) MS fragmentation of psoralen standard (b) and CYP71AJ20 product (d).

(c, e) MS^2^ fragmentation of psoralen standard (c) and the product (e) at *m/z* 187.04.

**Figure S20.** Columbianetin synthase activity of CYP736A530.

(a) UV chromatogram of *in vitro* yeast microsome assay of CYP736A530 and control using osthenol as the substrate.

(b, d) MS fragmentation of columbianetin standard (b) and CYP736A530 product (d).

(c, e) MS^2^ fragmentation of columbianetin standard (c) and the product (e) at *m/z* 247.10.

**Figure S21.** Biosynthetic pathway of TCs with transcriptome and metabolite data in *A. keiskei*.

The transcriptome and metabolite data were overlaid onto the TC biosynthetic pathway of Apiaceae. Each arrow represents a single enzymatic reaction. Cytochrome P450 reactions identified in this study are shown in red. Solid lines indicate known genes, and dashed lines indicate unknown genes. Gene expression heatmaps are based on mean-standardized log₂ fold changes. Metabolite accumulation levels are shown as mean-standardized log₂ values across all samples. Organs and metabolites that were not detected are indicated in gray (n.d.). PAL, phenylalanine ammonia-lyase; C4H, cinnamate 4-hydroxylase; 4CL, 4-coumaroyl: CoA ligase; C2’H, p-coumaroyl CoA 2’-hydroxylase; U6DT, umbelliferone 6-dimethylallyltransferase; U8DT, umbelliferone 8-dimethylallyltransferase; DC, demethylsuberosin cyclase; PS, psoralen synthase; VS, vaginidiol synthase; AS, angelicin synthase; P5H, psoralen 5-hydroxylase; P8H, psoralen 8-hydroxylase; OMT, O-methyltransferase; B5ODT, bergaptol 5-O-dimethylallyltransferase; X8ODT, xanthotoxol 8-O-dimethylallyltransferase; OS, oxypeucedanin synthase, ImE, imperatorin epoxidase; B8H, bergapten 8-hydroxylase; X5H, xanthotoxin 5-hydroxylase.

**Figure S22**. A preliminary analysis of the optimal pH for the vaginidiol synthesis activity of CYP71AJ62. Single reactions were performed at pH 5.0−9.0. The peak areas corresponding to vaginidiol formation were measured at 330 nm.

**Figure S23.** Lomatin 10-hydroxylation activity of CYP71AJ62.

(a) UV chromatogram of in vitro microsome assay of CYP71AJ62 and control using lomatin as a substrate. (b, c) MS fragmentation of khellactone standard (b) and CYP71AJ62 product (c).

(d, e) MS^2^ fragmentation of khellactone standard (d) and the product (e).

**Figure S24**. Kinetic analysis of CYP71AJ62 with columbianetin and lomatin as substrates.

Apparent *K*_m_ values for columbianetin (a) and lomatin (b) were calculated using nonlinear least squares fitting. Kinetic assays were independently repeated three times.

**Figure S25.** Kinetic analysis of CYP71AJ4 with columbianetin as a substrate.

Apparent *K*_m_ values for angelicin (a) and vaginidiol (b) production were calculated using nonlinear least squares fitting. Kinetic assays were independently repeated three times.

**Figure S26.** A preliminary analysis of the optimal pH for the oxypeucedanin synthesis activity of CYP71AJ63. Single reactions were performed at pH 5.0−9.0. The peak areas corresponding to the formation of oxypeucedanin were measured at 310 nm.

**Figure S27**. Kinetic analysis of CYP71AJ63 with isoimperatorin as a substrate.

The apparent *K*_m_ value for oxypeucedanin production was calculated using nonlinear least squares fitting. Kinetic assays were independently repeated three times.

**Figure S28**. Substrate preference of CYP71AJ63.

The isoimperatorin epoxidase activity was set to 100 %. Data are shown as mean ± standard deviation from three independent experiments. n.d., not detected.

**Figure S29**. Three-dimensional homology models of CYP71AJ4 and CYP71AJ62.

Three-dimensional homology models of CYP71AJ4 and CYP71AJ62 are shown in khaki and pale orange, respectively.

(a) Columbianetin docking with CYP71AJ4 and 62 is shown in yellow and orange, respectively. The heme and heme iron are shown in gray and lime, respectively. Oxygen and nitrogen atoms of the heme are shown in red and blue.

(b,c) Orientation of columbianetin toward the heme iron in CYP71AJ4 (b) and 62 (c). The heme-binding oxygen atom (red sphere) is set 1.7 Å from the heme iron (Rupasonghe et al., 2003). Distances from the heme-binding oxygen to C9 are shown.

**Figure S30.** Presumed reaction mechanisms catalyzed by CYP71AJ4 (yellow) and CYP71AJ62 (orange). The angelicin biosynthetic pathway was inferred based on Stanjek et al. (1999) and Larbat et al. (2009).

**Figure S31.** Sequence alignment of candidate P450s in the columbianetin clade of Fig. 5a.

Amino acid sequences deduced from the 11 candidates were aligned using MAFFT v7.490. All analyses were performed using Geneious Prime 2024.2.1 (Dotmatics). Known angelicin synthases are indicated by yellow circles, while the vaginidiol synthase of *A. keiskei* is indicated by orange circles. The SRSs and motifs were annotated according to Dueholm et al. (2015) and Fang et al. (2024). The transmembrane region was estimated using TargetP-2.0 (Armenteros et al., 2019). Residues located within 5 Å of the substrate in the three-dimensional structural analysis are highlighted with cyan boxes.

**Supplementary Data S1.** Newick data for the 286 sequences selected based on OrthoFinder results

