## Supplementary Figure S1 to S29 for "Molecular characterization of a cytochrome P450 involved in the skeletal diversification of tricyclic coumarins in *Angelica keiskei*"

**This PDF file includes:**
Figures S1 to S31

**Other supplementary materials for this manuscript include the following:**
Tables S1 to S19
Suppramentary Data S1

| Third ring | Position where the third ring is substituted | |
|---|---|---|
| | 6,7-fused | 7,8-fused |
| furan | 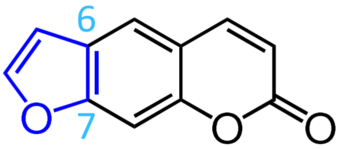 Psoralen | 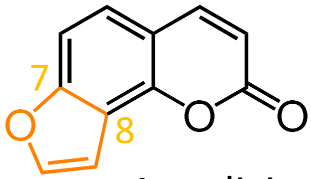 Angelicin |
| dihydrofuran | 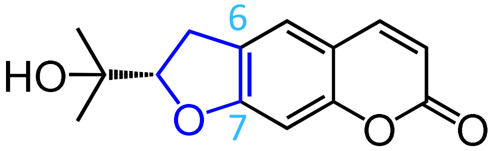 Marmesin | 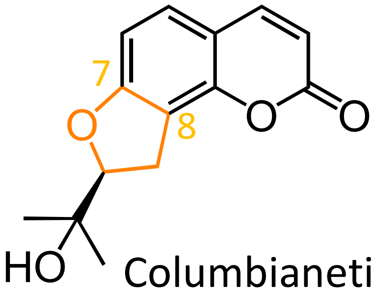 Columbianetin |

**Figure S1.** The core structures of tricyclic coumarins in this study.

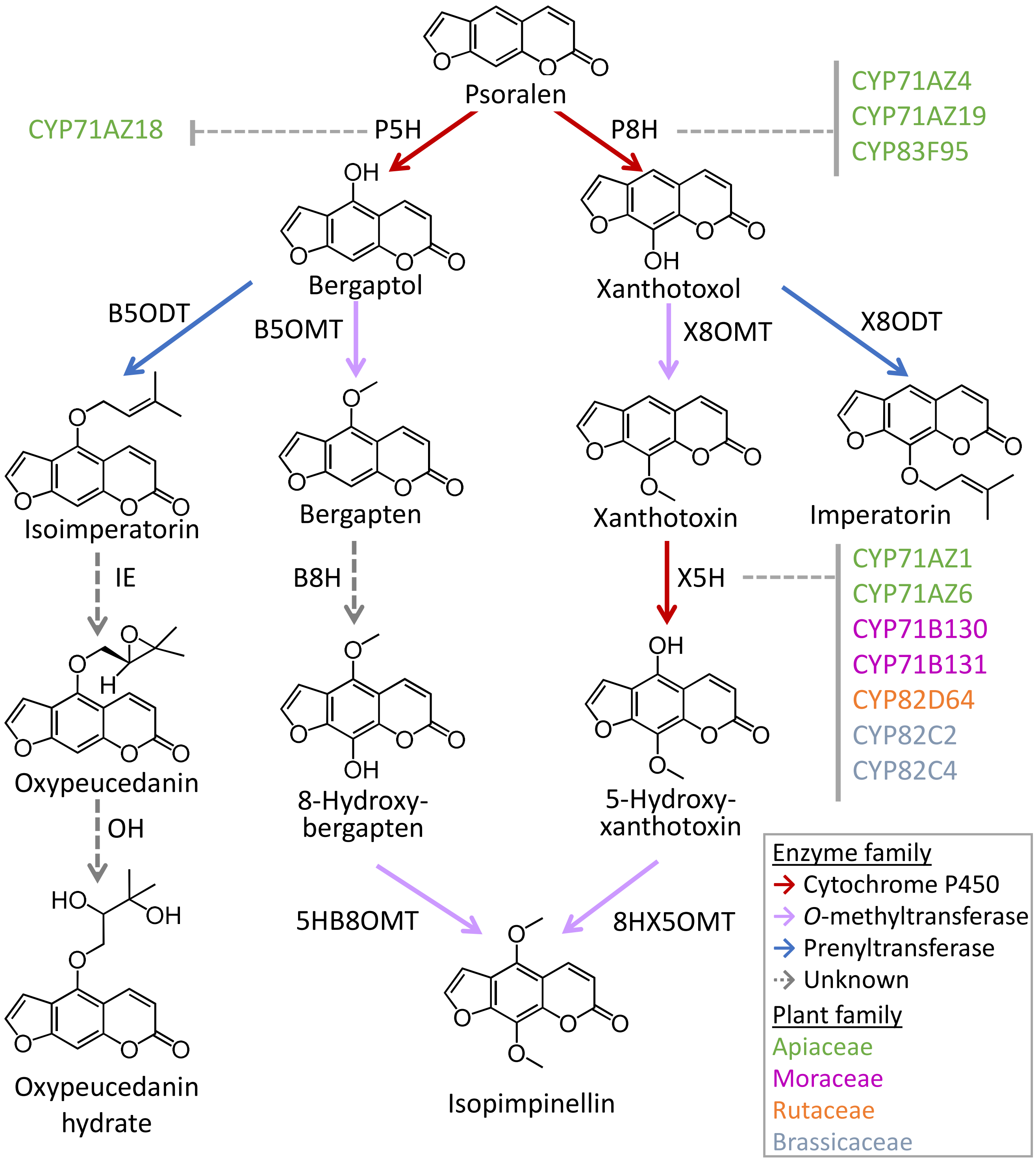


**Figure S2.** Overview of the modification pathways of linear furanocoumarins in angiosperms.
All biosynthetic enzymes of linear FCs described are shown in solid arrows (Hehmann et al., 2004; Kruse et al., 2008; Limones-Mendez et al., 2020; Zhang et al., 2022; Bouillé et al., 2025; Ji et al., 2025). Cytochrome P450s are labelled with their IDs and color-coded according to the plant family. Abbreviations: *B8H, bergapten 8-hydroxylase; B5ODT, bergaptol 5-O-dimethyallyltransferase; B5OMT, bergaptol 5-O-methyltransferase; 5HB8OMT, 5-hydroxybergapten 8-O-methyltransferase; 8HX5OMT, 8-hydroxyxanthotoxin 5-O-methyltransferase; IE, isoimperatorin epoxidase; OH, oxypeucedanin hydralase; P5H, psoralen 5-hydroxylase; P8H, psoralen 8-hydroxylase; X5H, xanthotoxin 5-hydroxylase; X8ODT, xanthotoxol 8-O-dimethyallyltransferase; X8OMT, xanthotoxol 8-O-methyltransferase.*

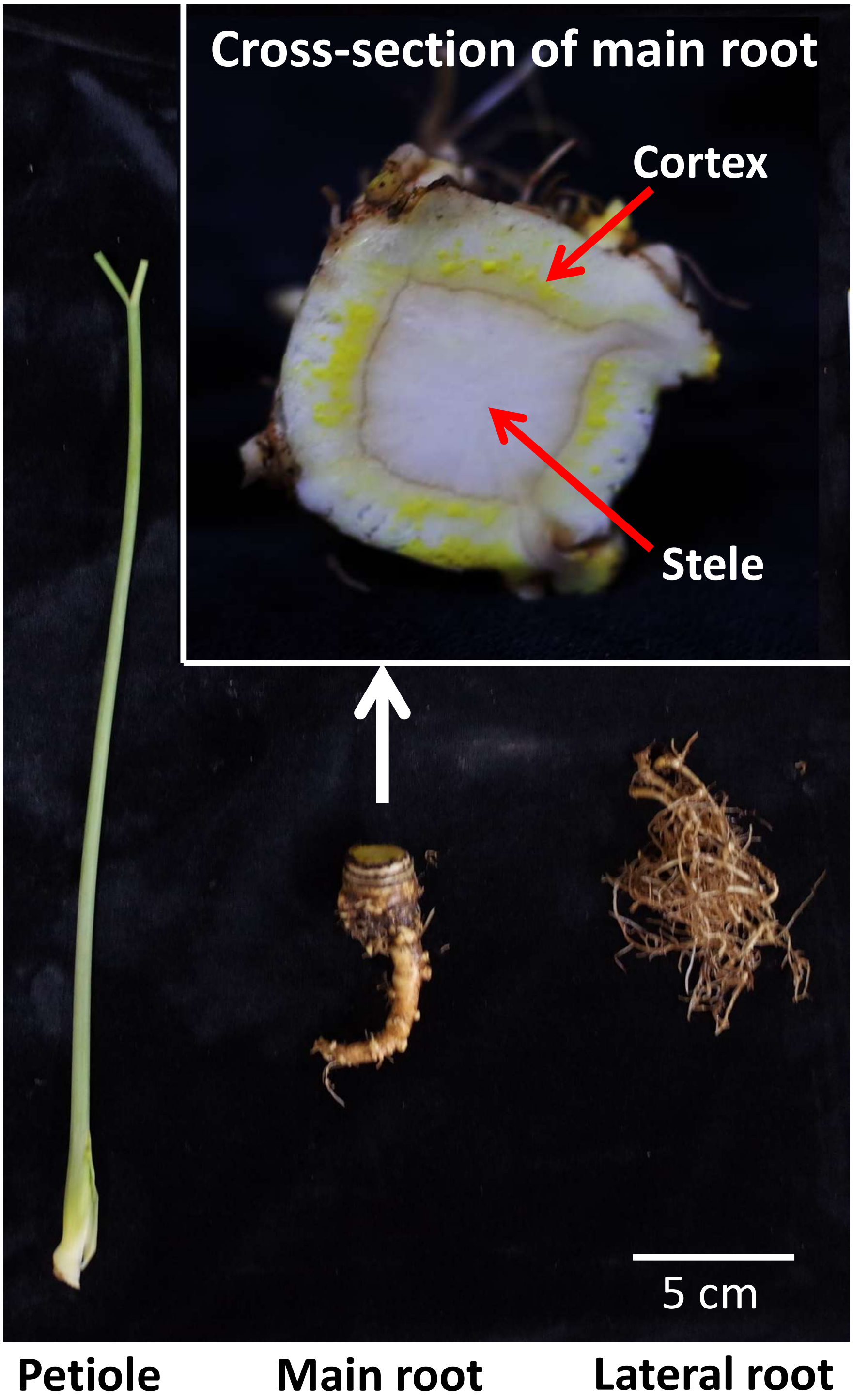


**Figure S3.** The petiole, main root cortex, main root stele, and lateral root of *A. keiskei* used for analysis.

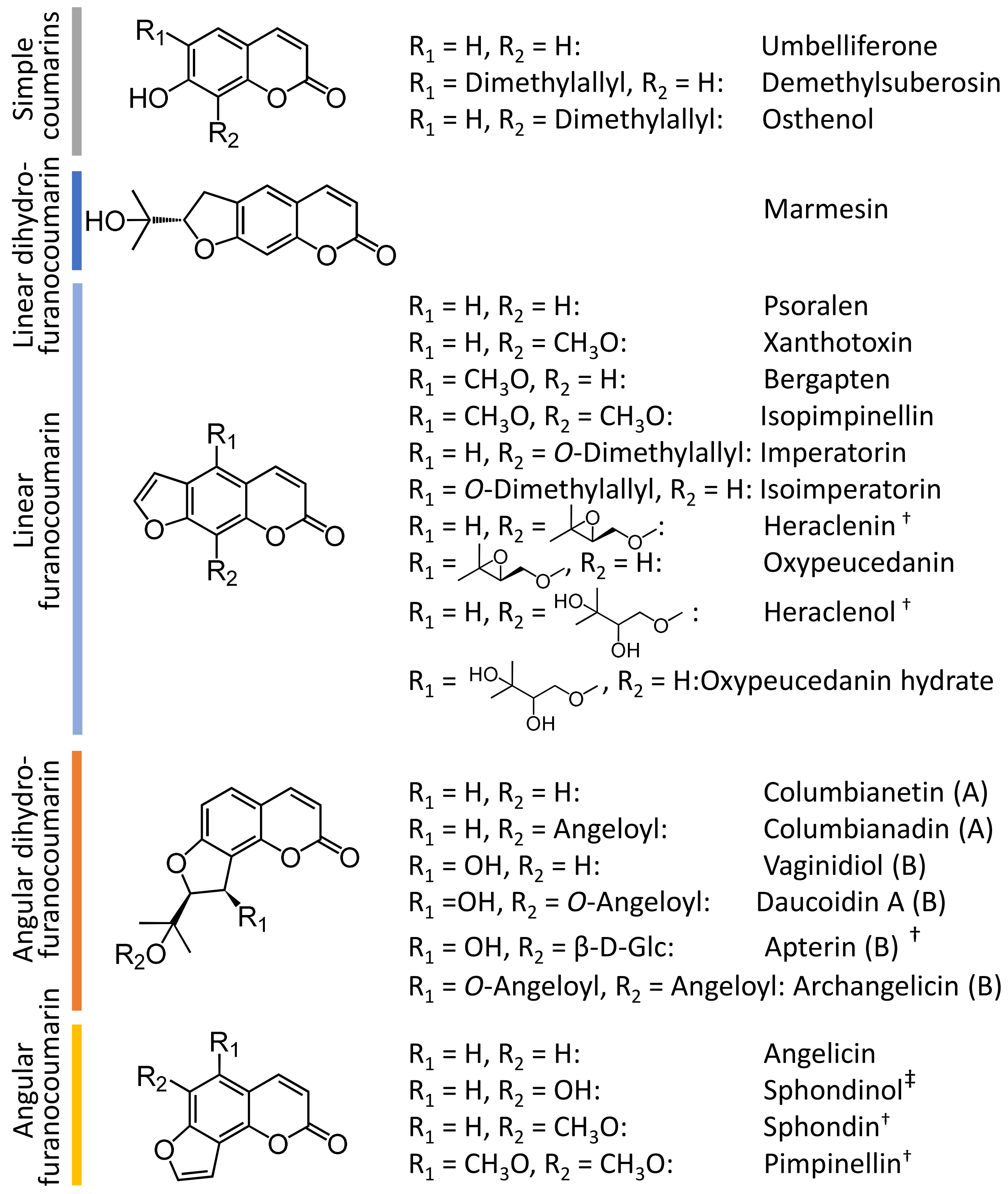


**Figure S4.** Quantified coumarin molecules in Angelica keiskei organs. The two types of dihydrofuranocoumarins (DHFCs), type-A and type-B, are shown as (A) and (B). In SciFinder registration, a † indicates the metabolites that have not been reported in *Angelica keiskei*, and ‡ indicates the metabolite that has not been reported in the genus *Angelica*.

### (a) Simple coumarins

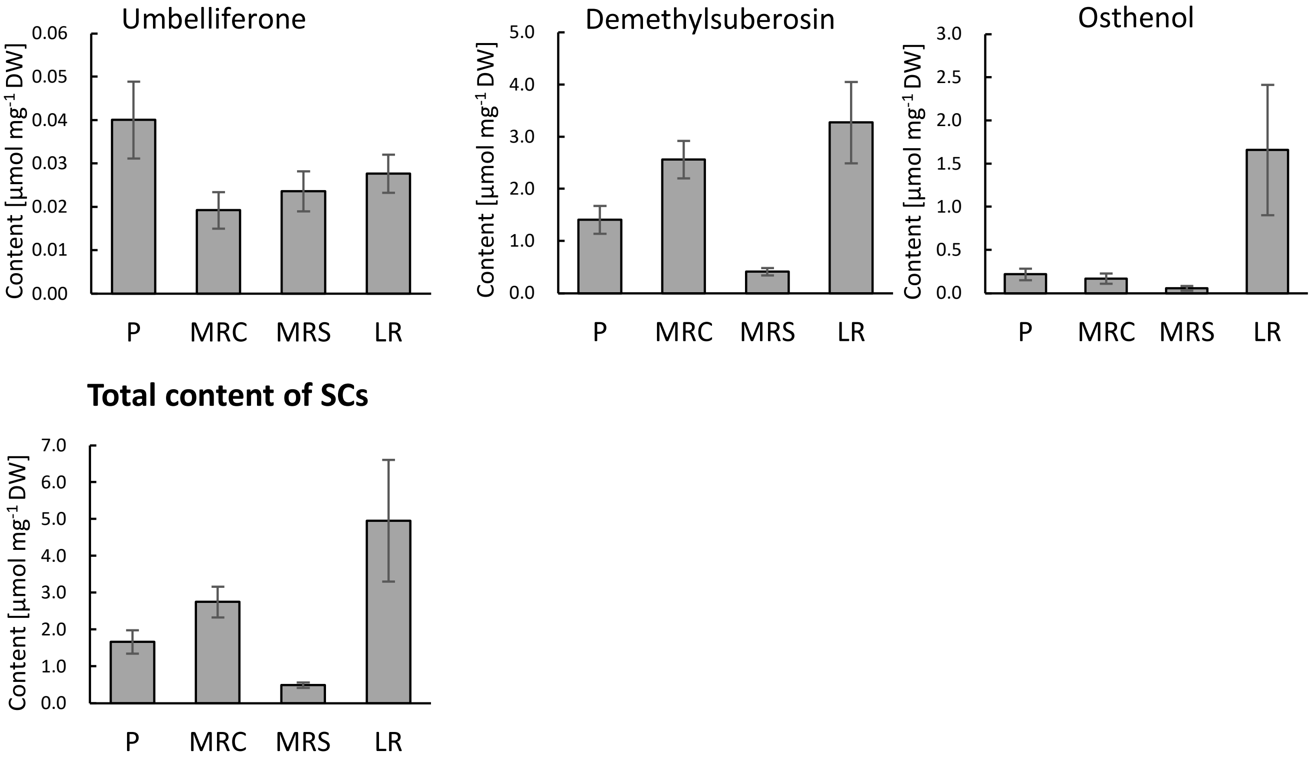


### (b) linear dihydrofuranocoumarin

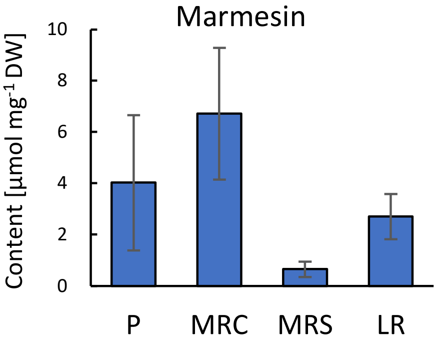


**Figure S5.** Tissue-specific accumulation of coumarins.

Shinya *et al.*

### (c) Linear furanocoumarins

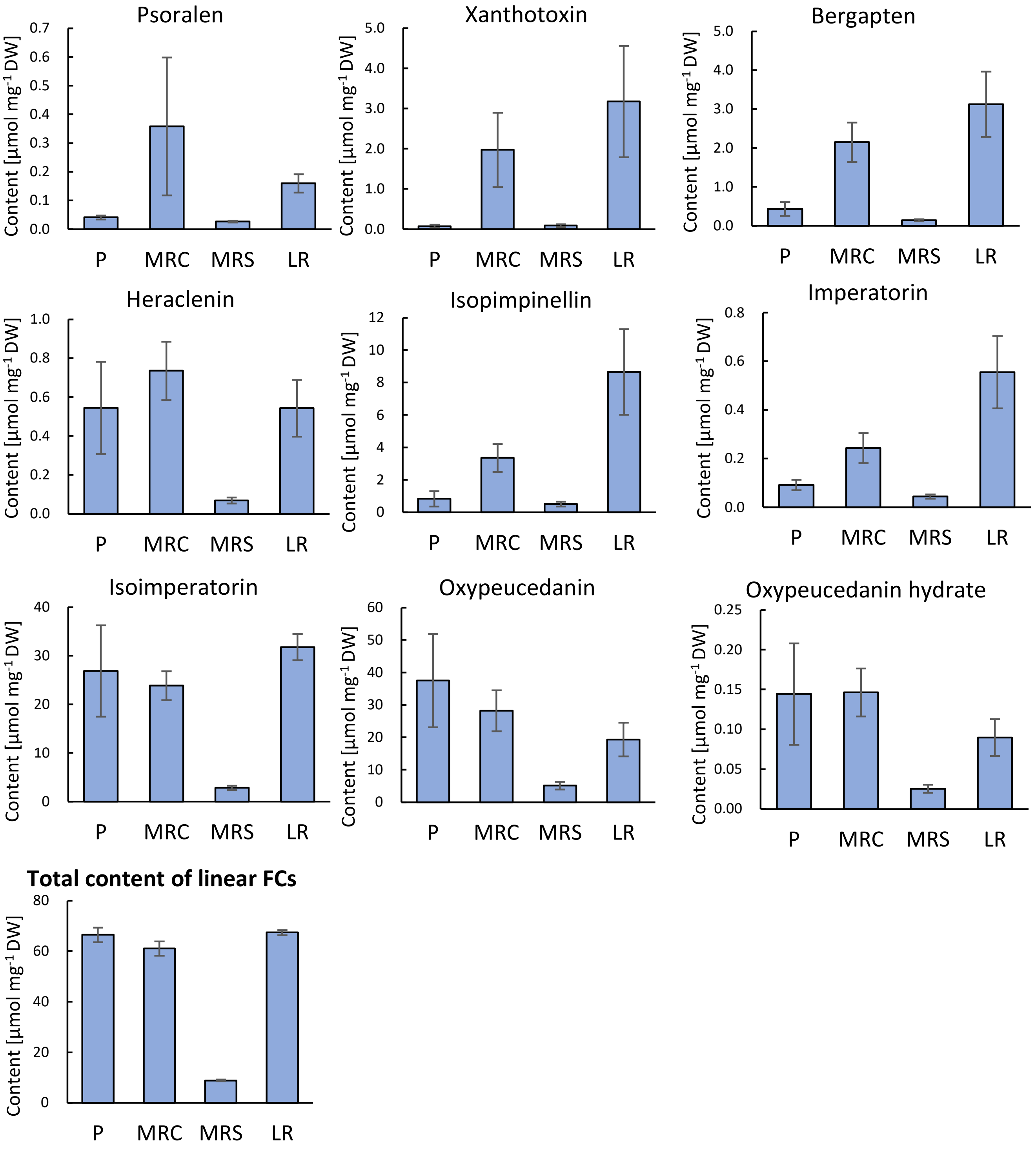


**Figure S5.** Tissue-specific accumulation of coumarins. *–continued*

### (d) Angular dihydrofuranocoumarins type A

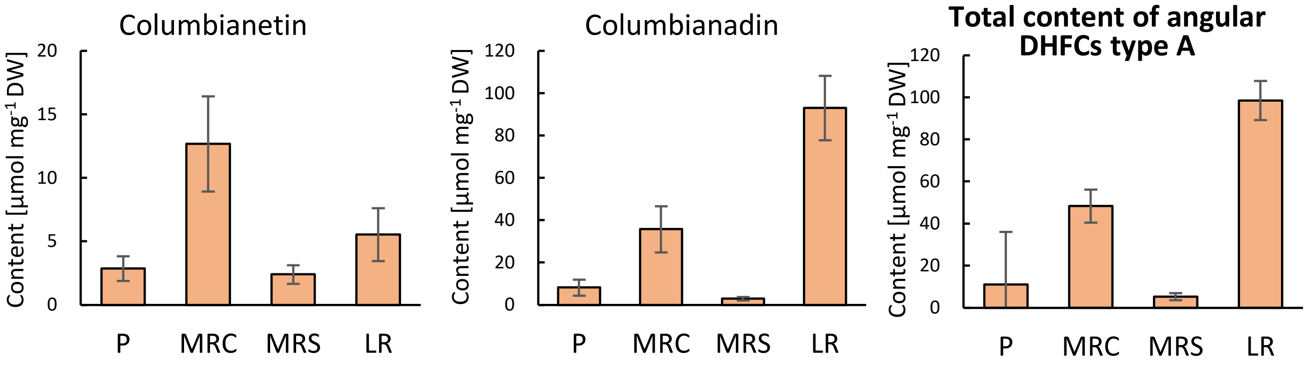


### (e) Angular dihydrofuranocoumarins type B

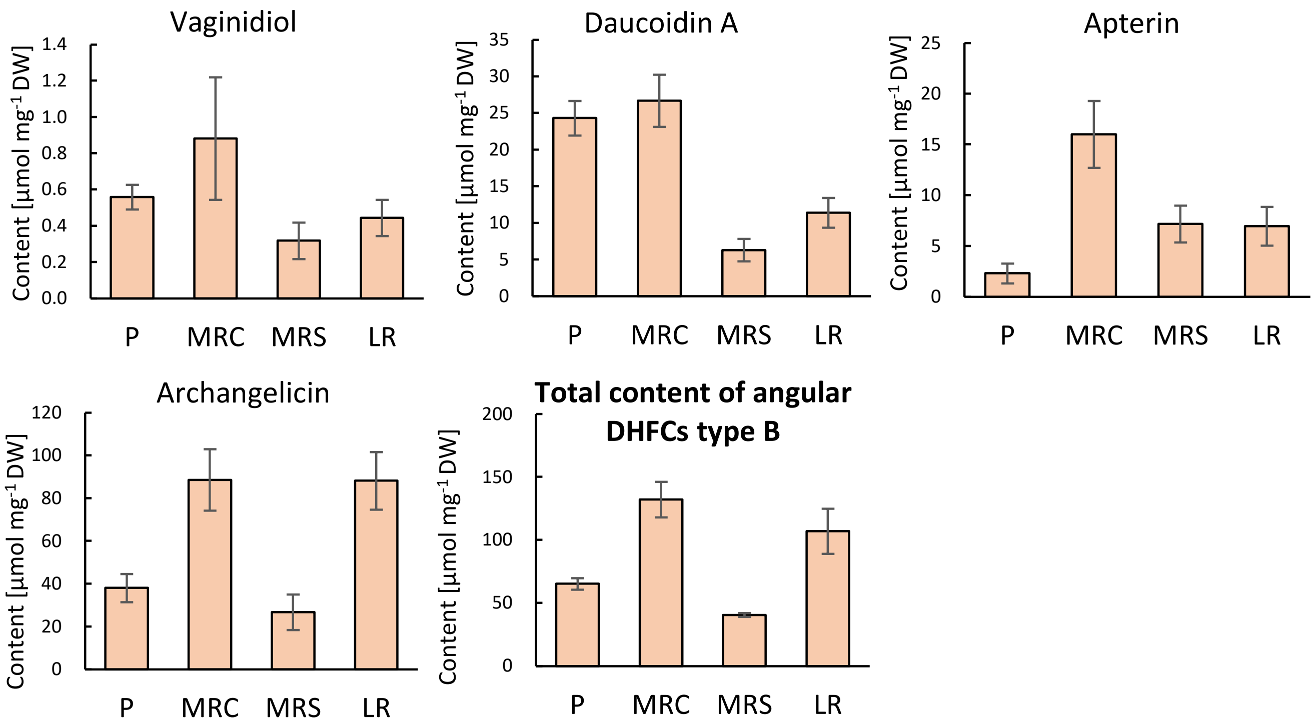


**Figure S5.** Tissue-specific accumulation of coumarins. *–continued*

### (f) Angular furanocoumarins

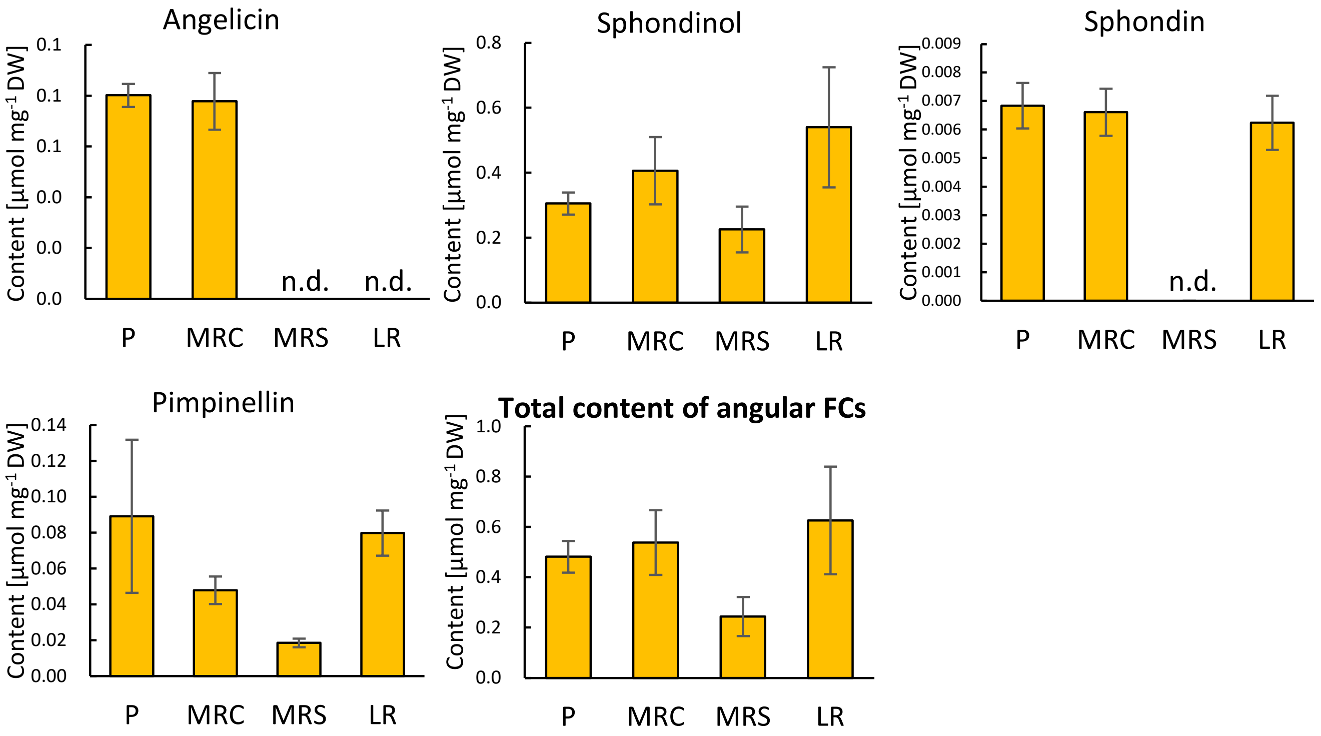


**Figure S5.** Tissue-specific accumulation of coumarins.
Data represent the contents of individual metabolites in simple coumarins (a), linear dihydrofuranocoumarin (b), linear furanocoumarins (c), angular dihydrofuranocoumarins type A (d), type B (e), and angular furanocoumarins (f) in the petiole (P), main root cortex (MRC), main root stele (MRS), and lateral root (LR). The total content of each coumarin group (a–f) is also shown. Data are presented as mean ± standard error from six individuals. The LC-MS detection parameters for the analyzed metabolites are listed in Table S14.

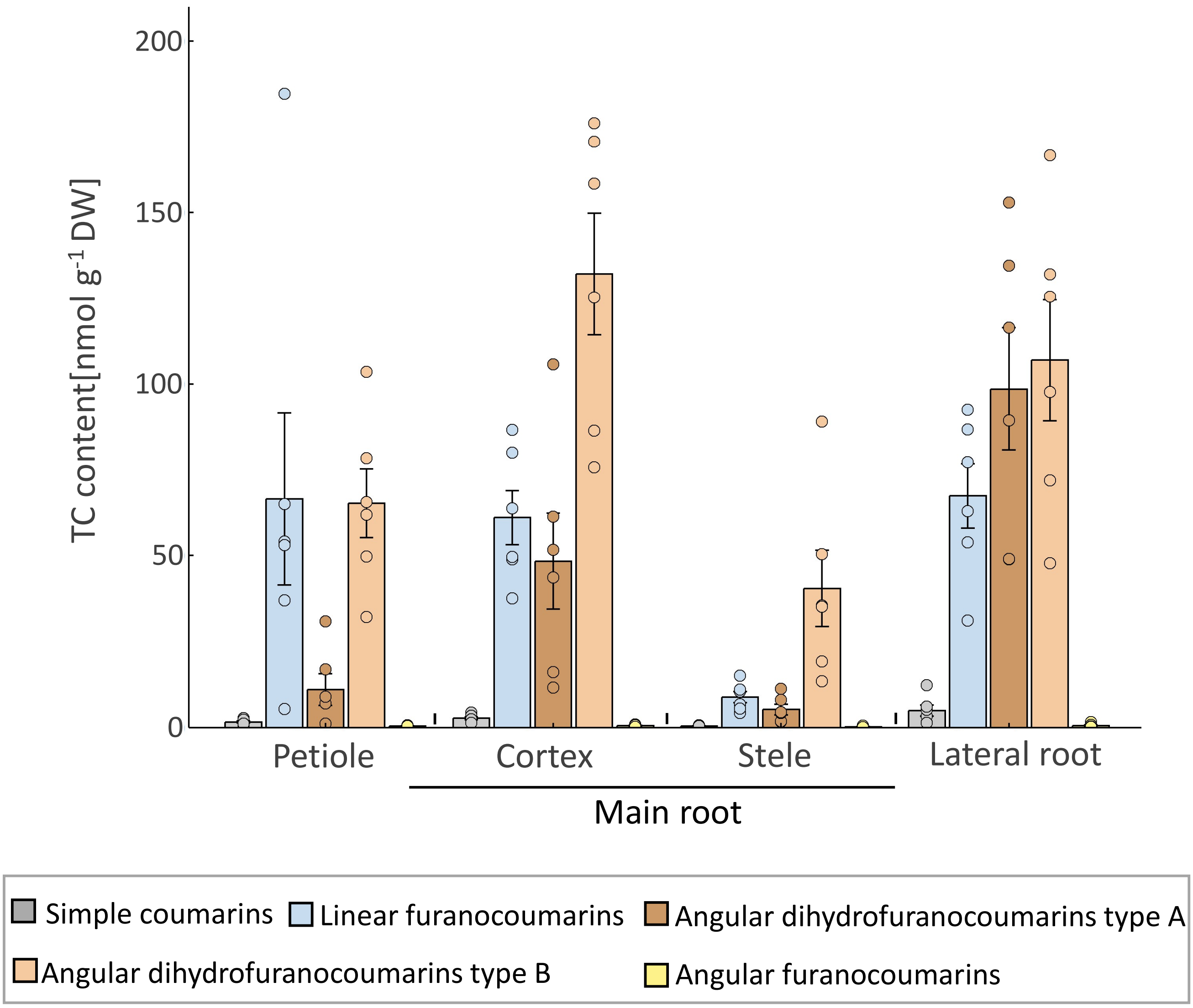


**Figure S6.** Coumarin accumulation patterns in *A. keiskei* organs.
TC accumulation in the petiole, main root cortex, main root stele, and lateral root. The accumulation amounts of simple coumarins, linear furanocoumarins, angular dihydrofuranocoumarins type A, type B, and angular furanocoumarins are shown in grey, blue, brown, orange, and yellow, respectively. Data are shown as mean ± standard error from six individuals. The LC-MS detection parameters for the metabolites analyzed are listed in Table S14.

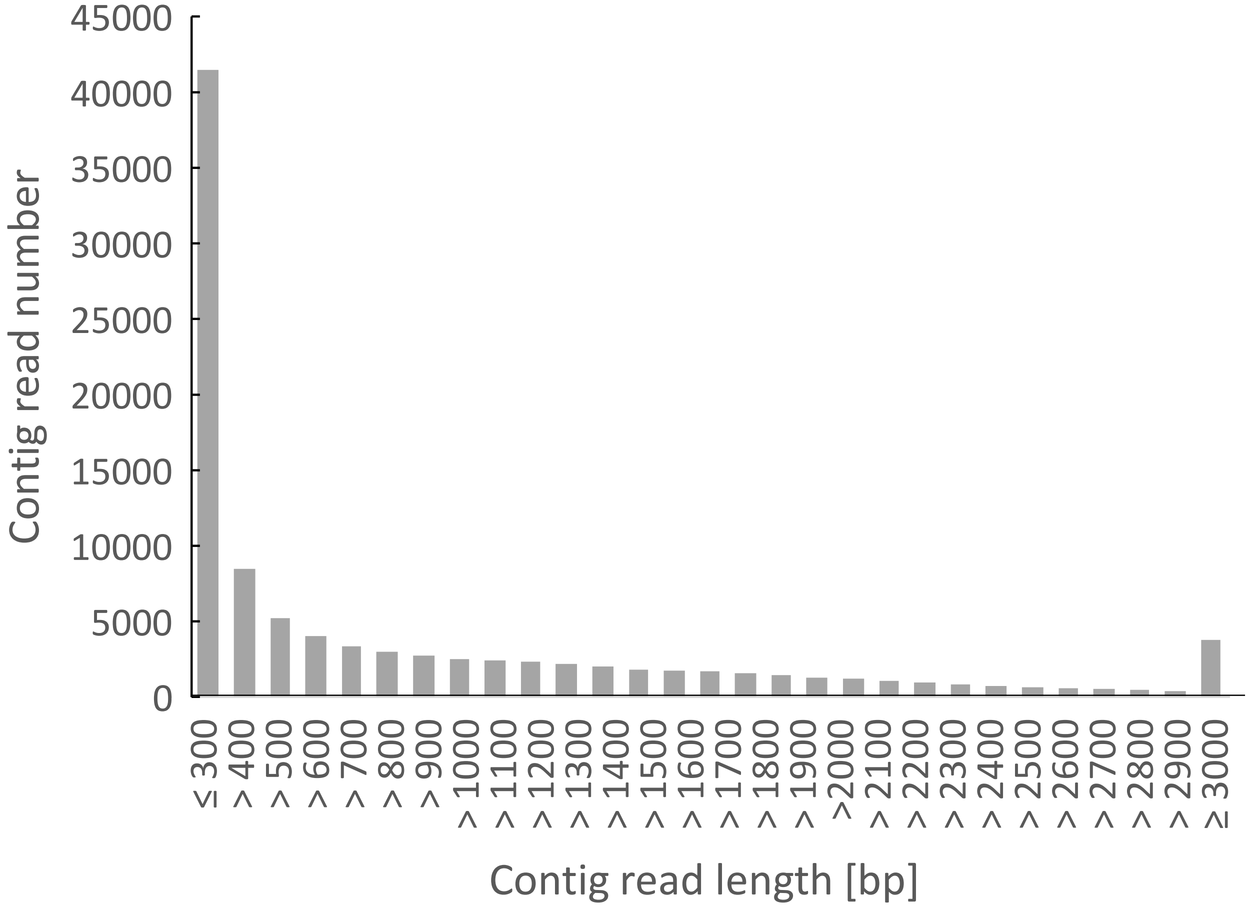


**Figure S7.** Histogram of contigs by sequence length.

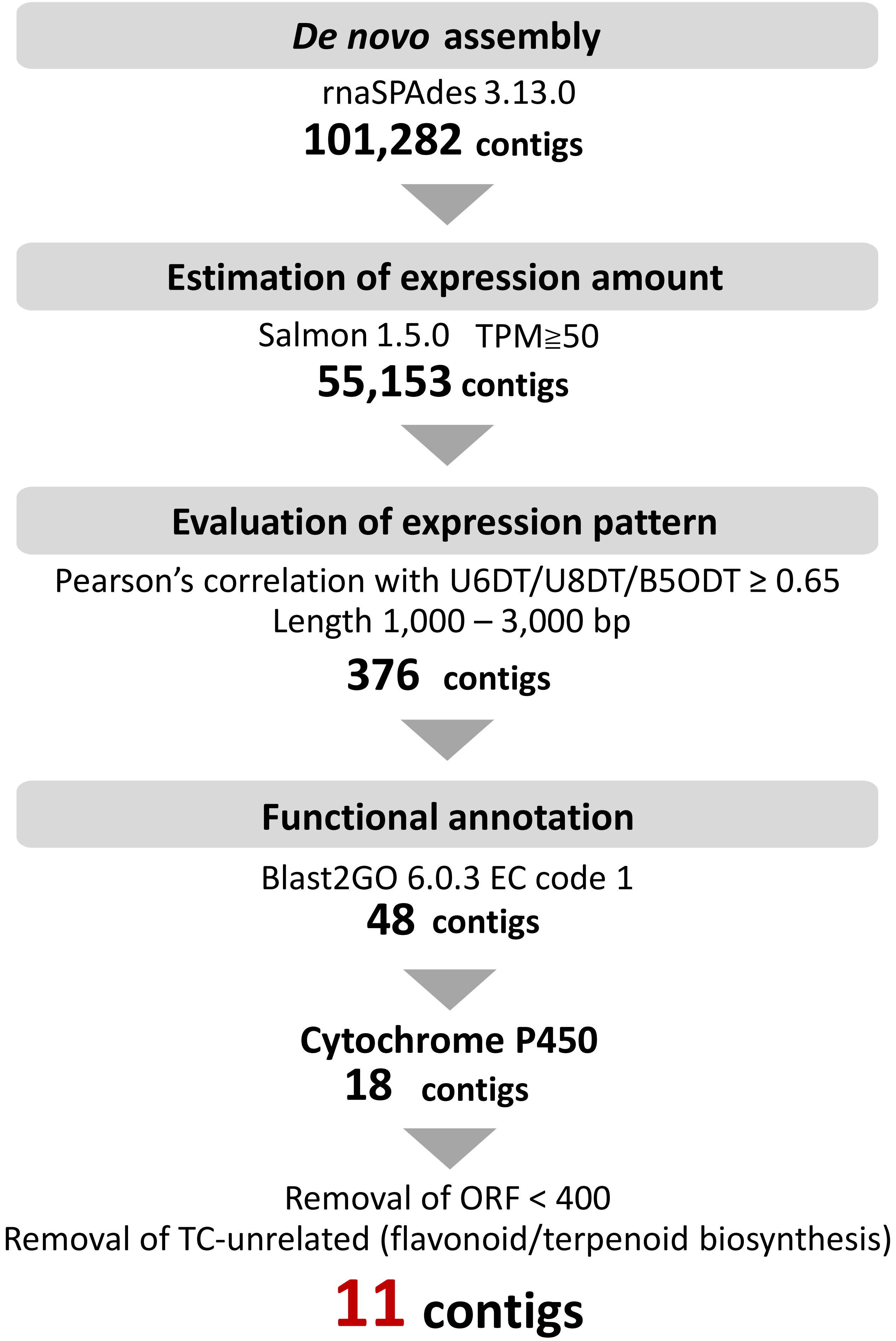


**Figure S8.** Stepwise filtering of candidate cytochrome P450 genes.

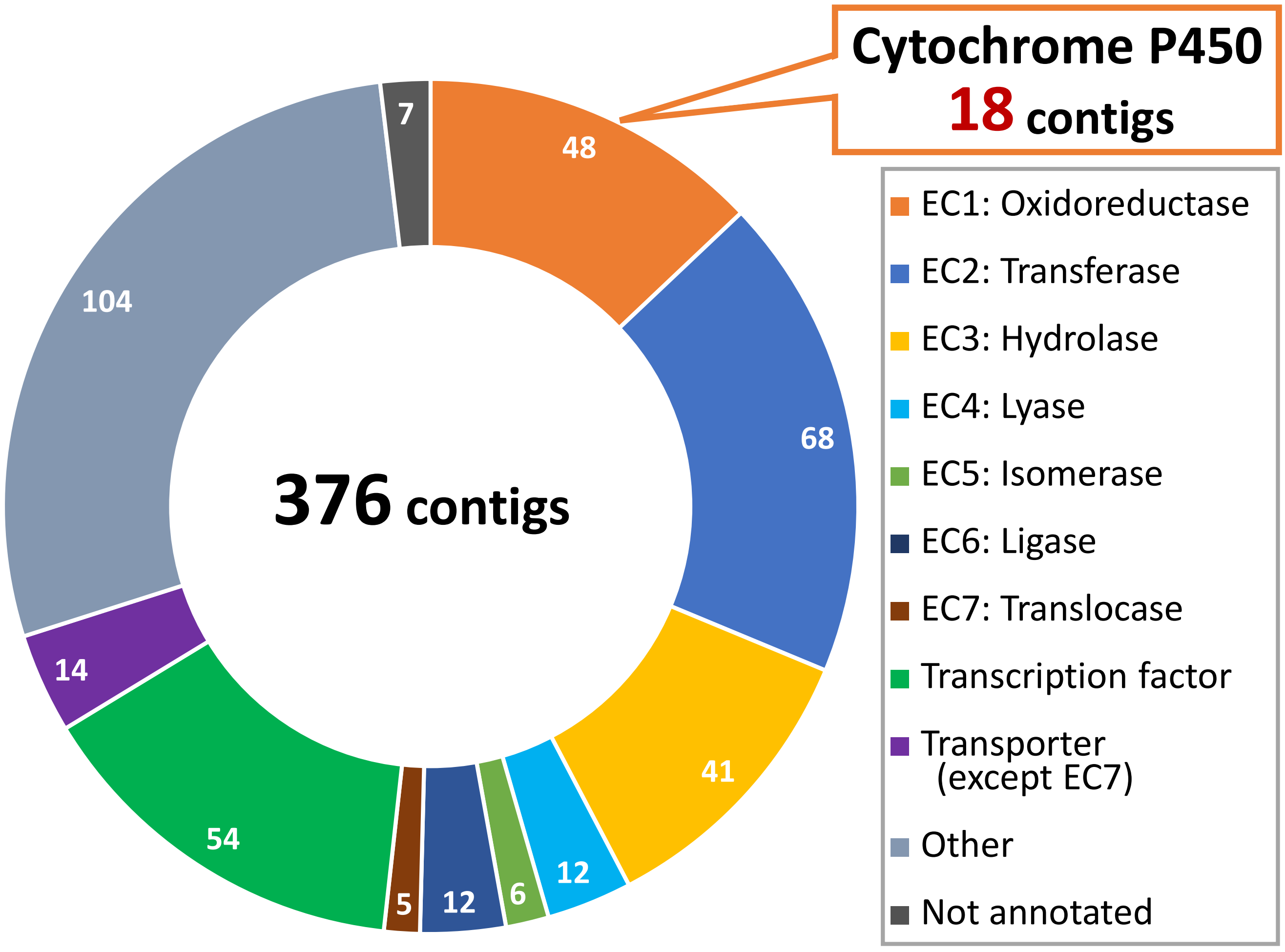


**Figure S9.** Functional annotations of contigs co-expressed with TC-biosynthetic PTs using Blast2GO.
Annotation of each contig was performed using Blast2GO.

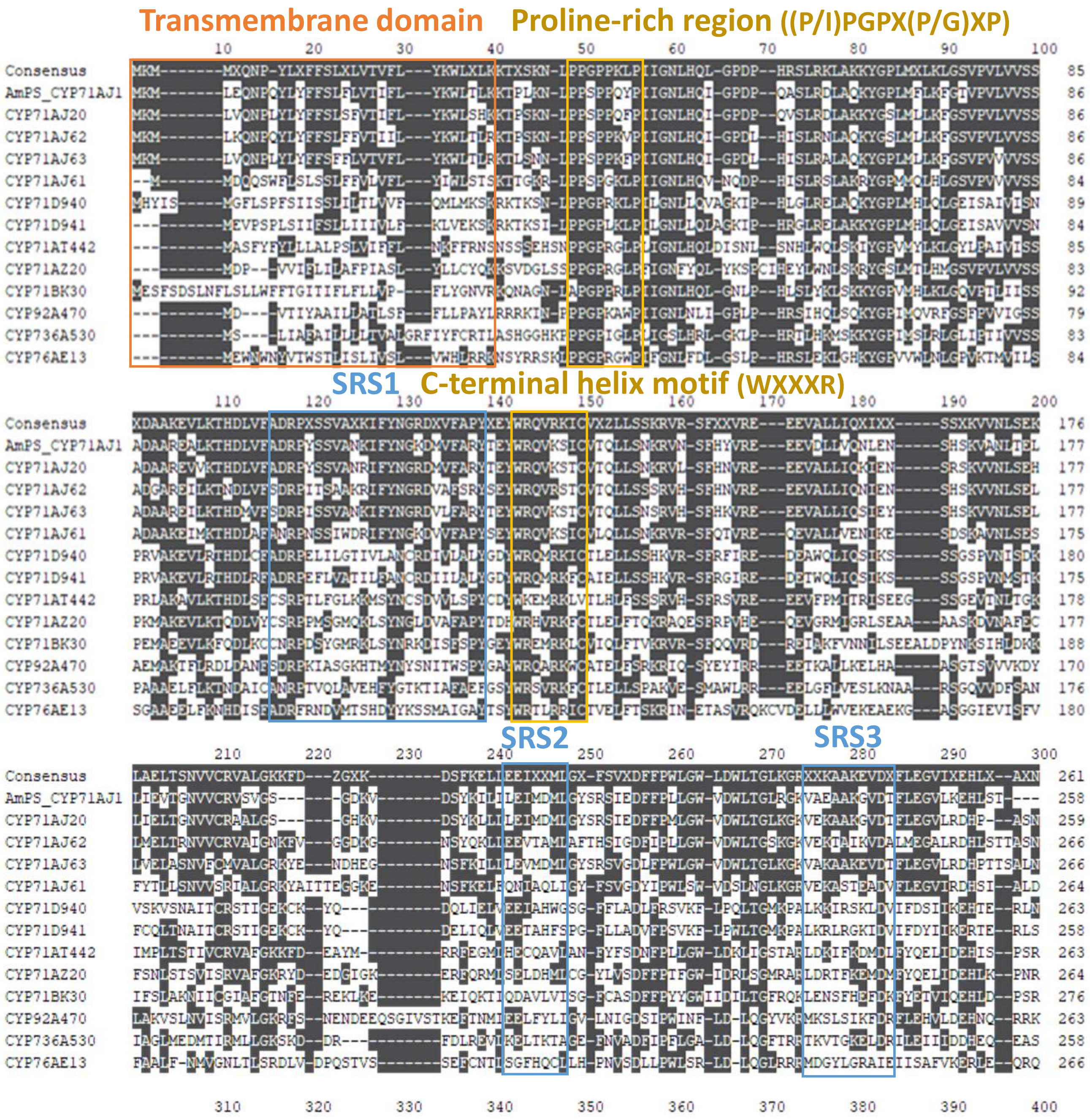


**Proline-rich region:** forms the proline hinge for conformational changes
**Substrate recognition sites (SRSs):** responsible for substrate recognition
**C-terminal helix:** related to the interaction with redox partners

**Figure S10.** Sequence alignment of candidate P450s and CYP71AJ1.

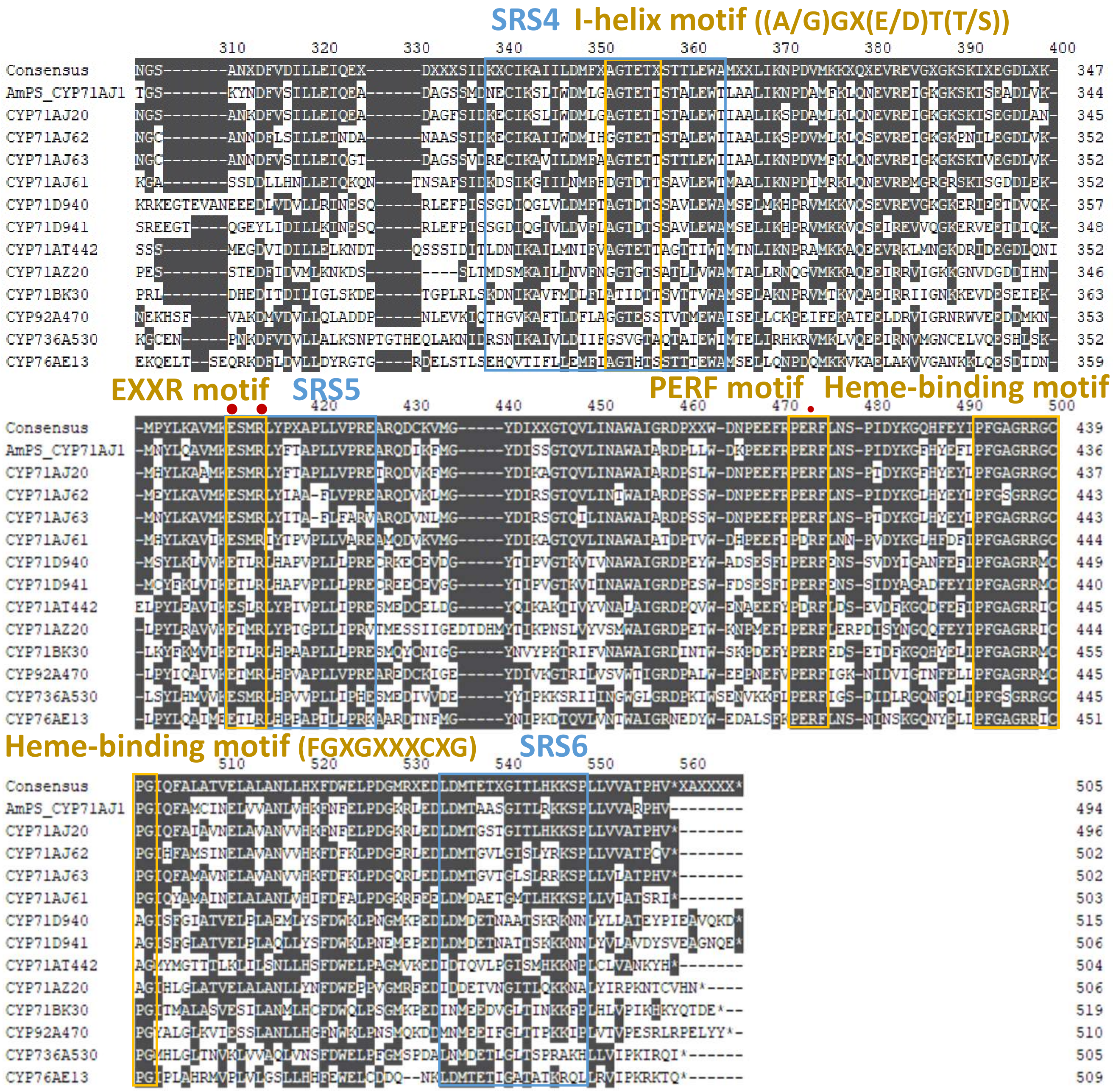


**I-helix motif:** associated with oxygen activation

**EXXR motif:** involved in proton transfer and structural stabilization

**PERF motif:** contributes to stabilization of the catalytic core

**E-R-R triad •:** contributes to heme fixation and stabilization

**Heme-binding motif:** serves as the axial ligand and coordinates the heme iron

**Figure S10.** A multiple alignment of the polypeptide sequences of candidate P450s and CYP71AJ1.
Amino acid sequences deduced from the eleven candidates and CYP71AJ1 were aligned using MAFFT v7.490 and visualized using Geneious Prime 2024. SRSs and motifs were annotated according to Dueholm et al. (2015) and Fang et al. (2024). The transmembrane region was estimated using TargetP-2.0 (Armenteros et al., 2019).

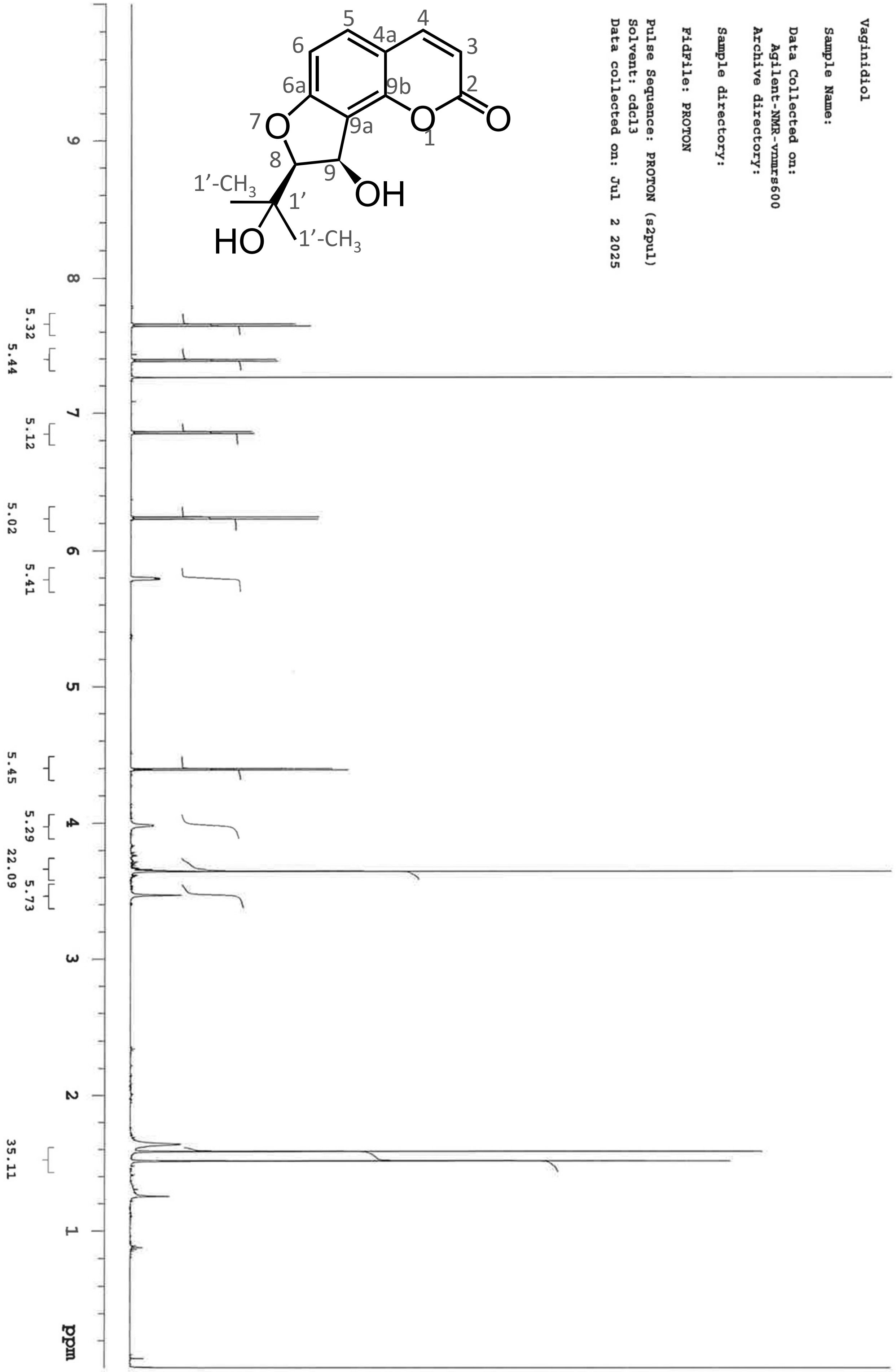


**Figure S11.** $^{1}$H-NMR spectrum of vaginidiol obtained by hydrolysis.

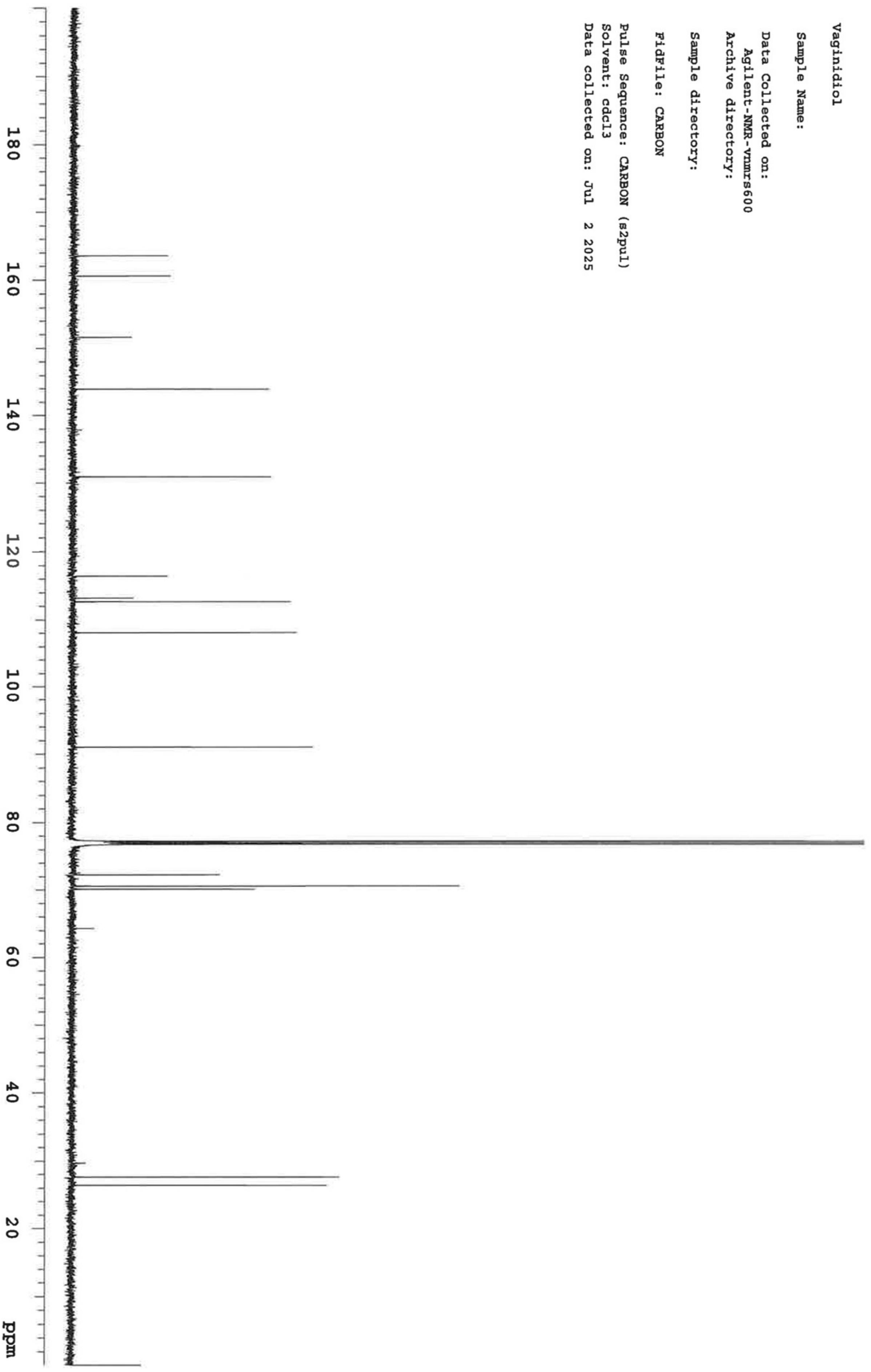


**Figure S12.** $^{13}$C-NMR spectrum of vaginidiol obtained by hydrolysis.

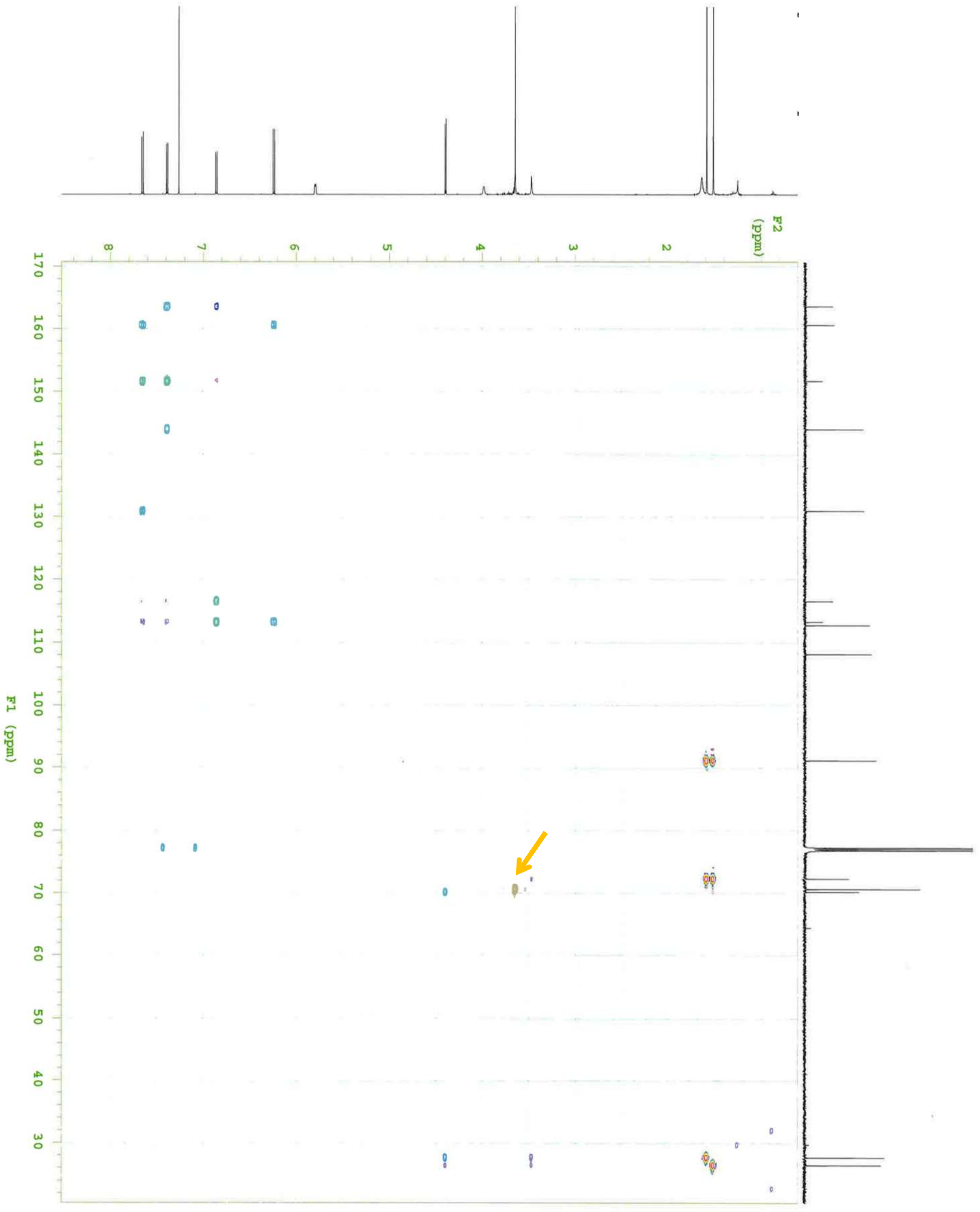


**Figure S13.** HMBC correlations of vaginidiol obtained by hydrolysis.
An unassigned methylene peak was detected in NMR spectra (indicated by a yellow arrow). This peak was regarded as an impurity due to no coupling with any proton or carbon of vaginidiol and no UV absorption.

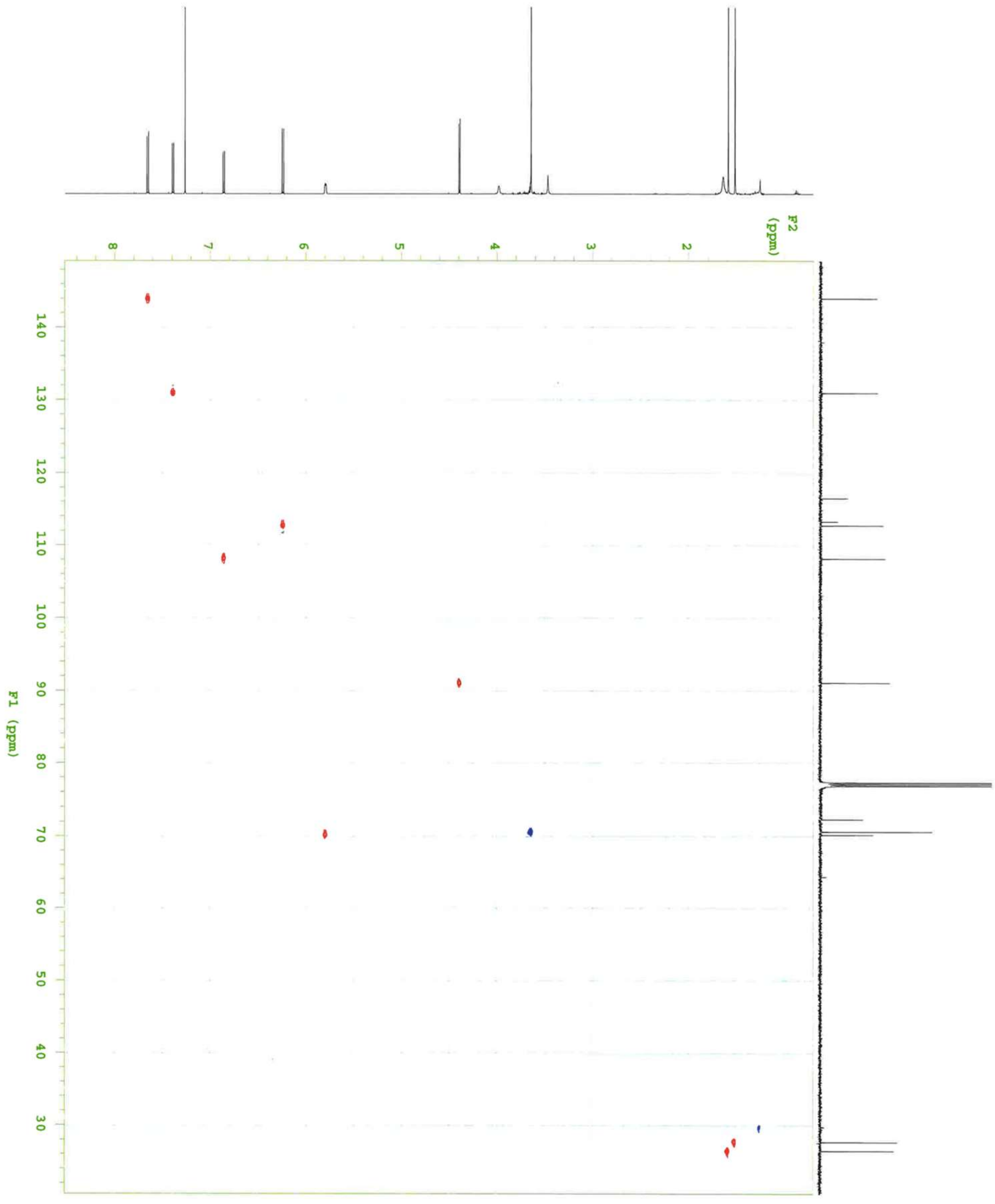


**Figure S14.** HSQC correlation of vaginidiol obtained by hydrolysis.

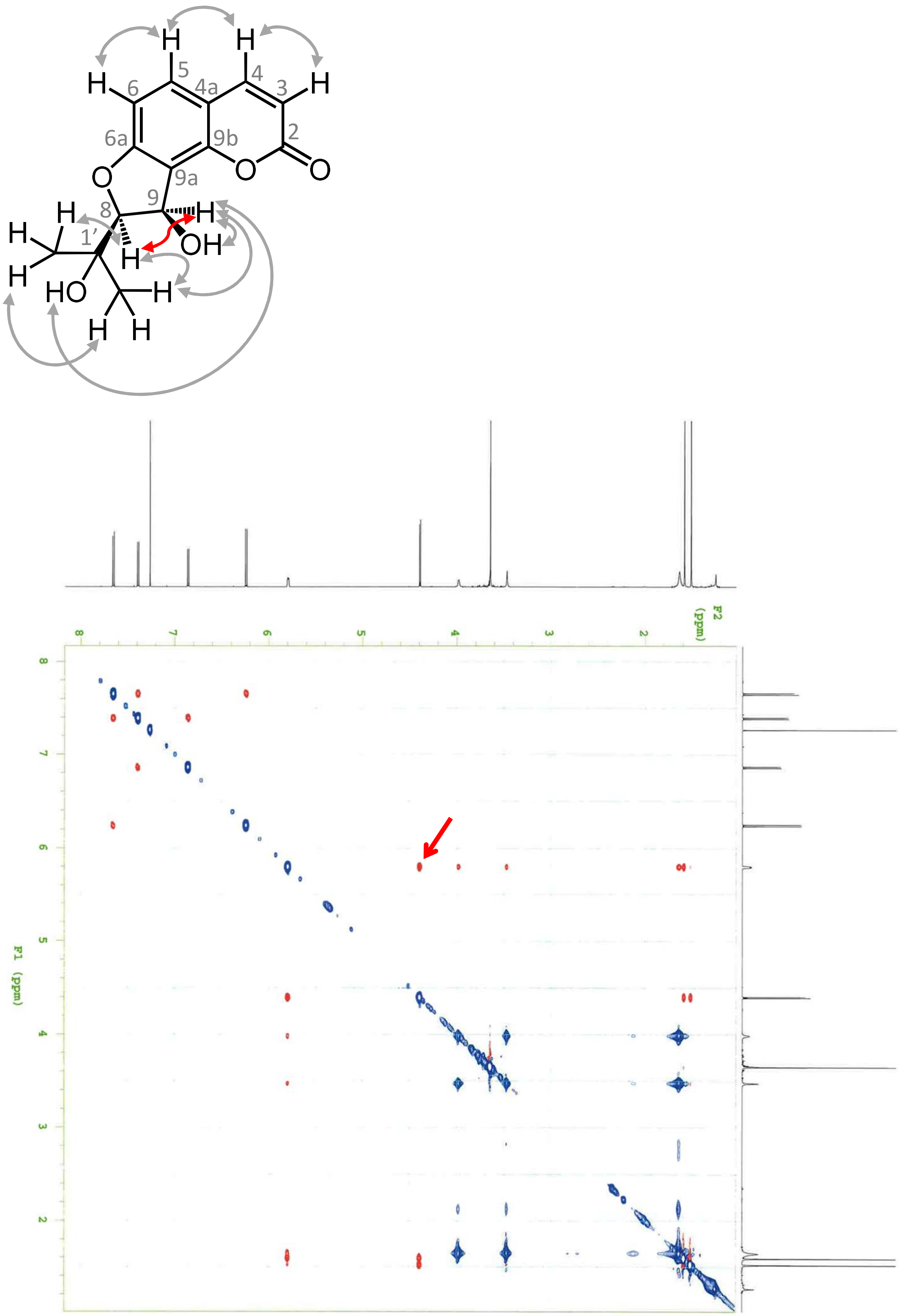


**Figure S15.** NOESY correlations of vaginidiol obtained by hydrolysis. The strong correlation between H-8 and H-9 was observed (shown in red arrows on the structure and NOESY plane), which demonstrates the *cis*-configuration.

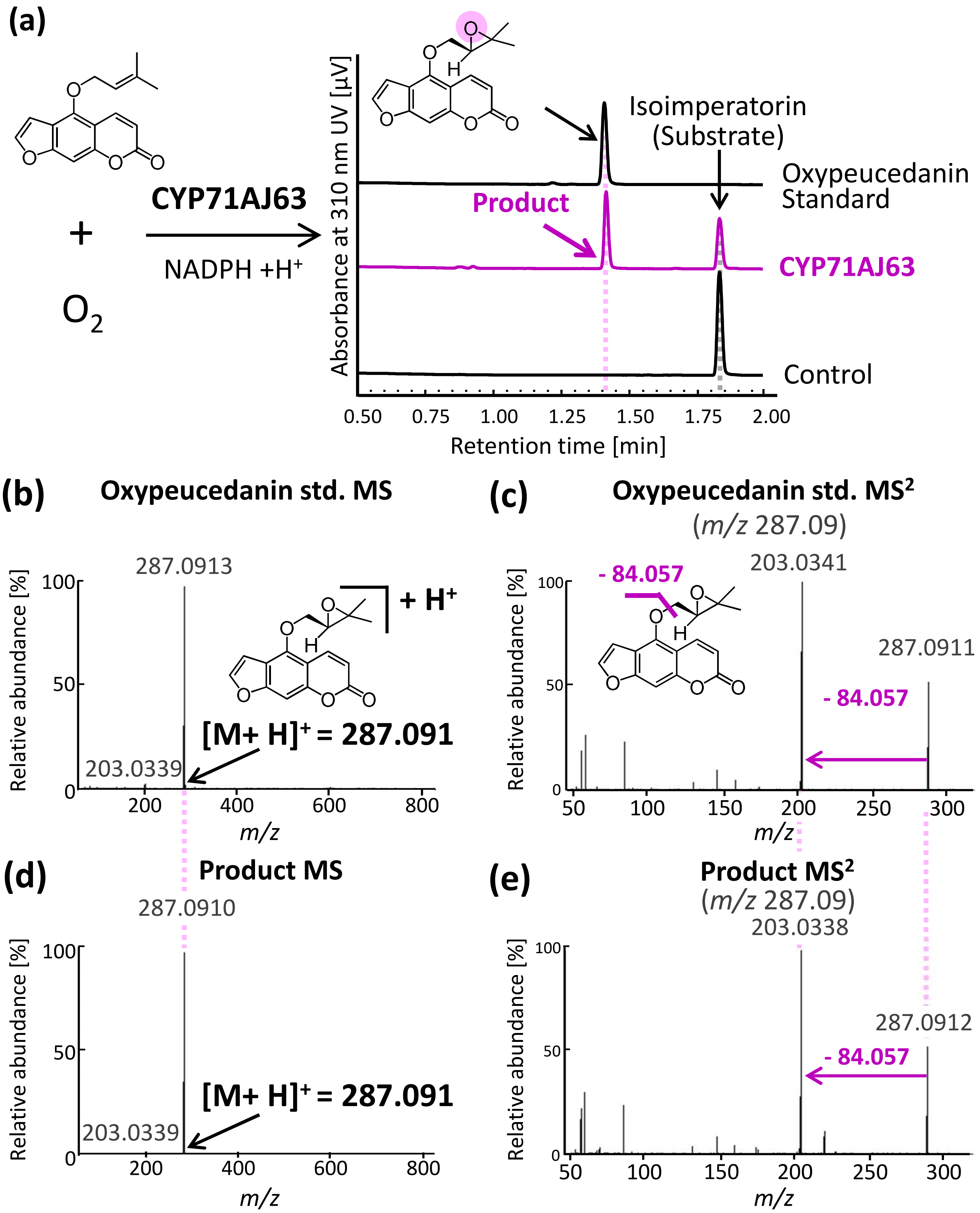


**Figure S16.** Oxypeucedanin synthase activity of CYP71AJ63.
(a) *In vitro* assay of CYP71AJ63 and control (AkCPR) using isoimperatorin as a substrate
(b, d) MS fragmentation of oxypeucedanin standard (b) and CYP71AJ63 product (d).
(c, e) MS2 fragmentation of oxypeucedanin standard (c) and the product (e) at *m/z* 287.09.

Shinya *et al.*

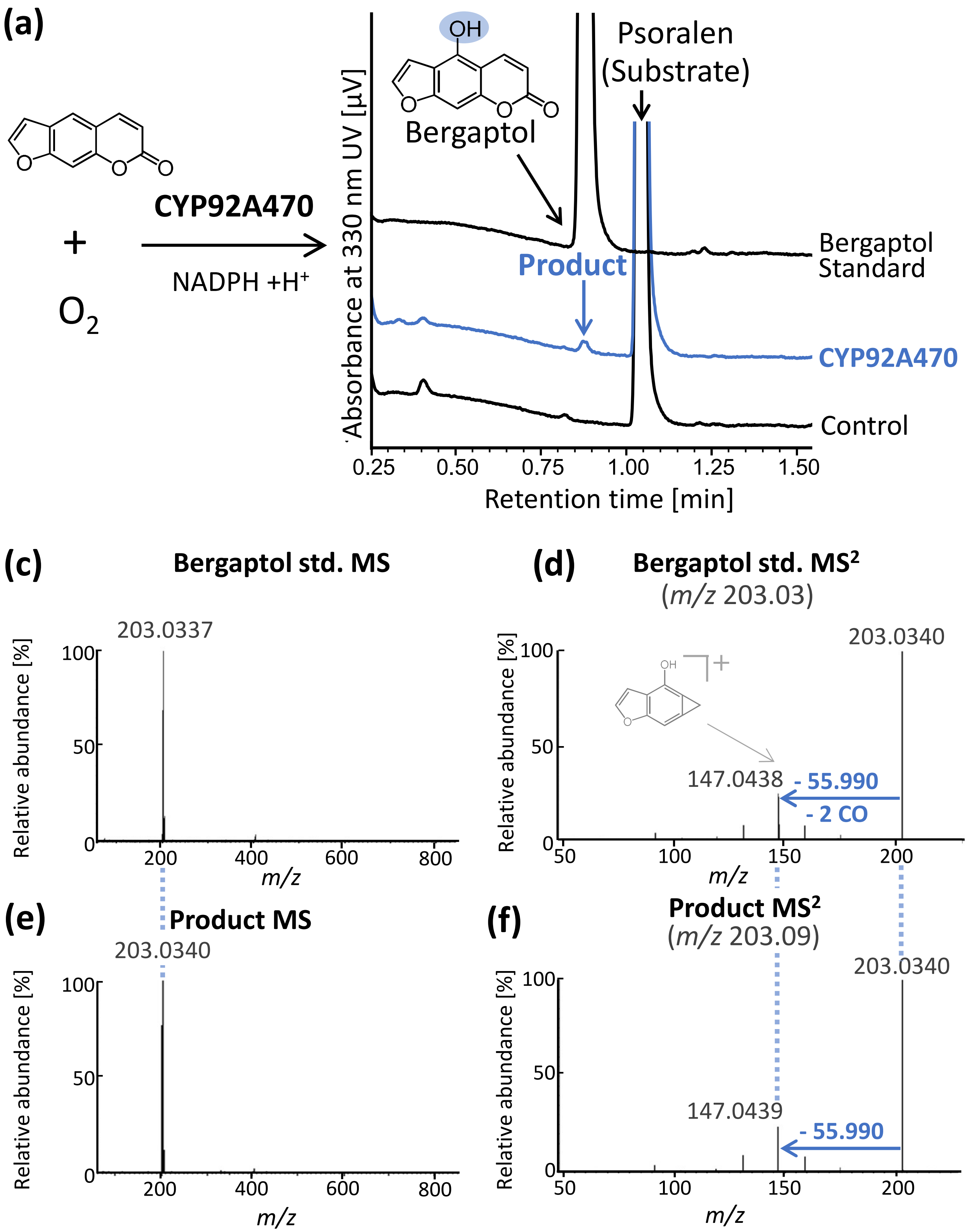


**Figure S17.** Psoralen 5-hydroxy activity of CYP92A470.
(a) UV chromatogram of *in vitro* yeast microsome assay of CYP92A470 using psoralen as a substrate
(b, d) MS fragmentation of bergaptol standard (b) and CYP92A470 product (d).
(c, e) MS$^2$ fragmentation of bergaptol standard (c) and the product (e) at *m/z* 203.03. The chemical structure of the *m/z* 147 fragment was assigned based on Du et al. (2025).

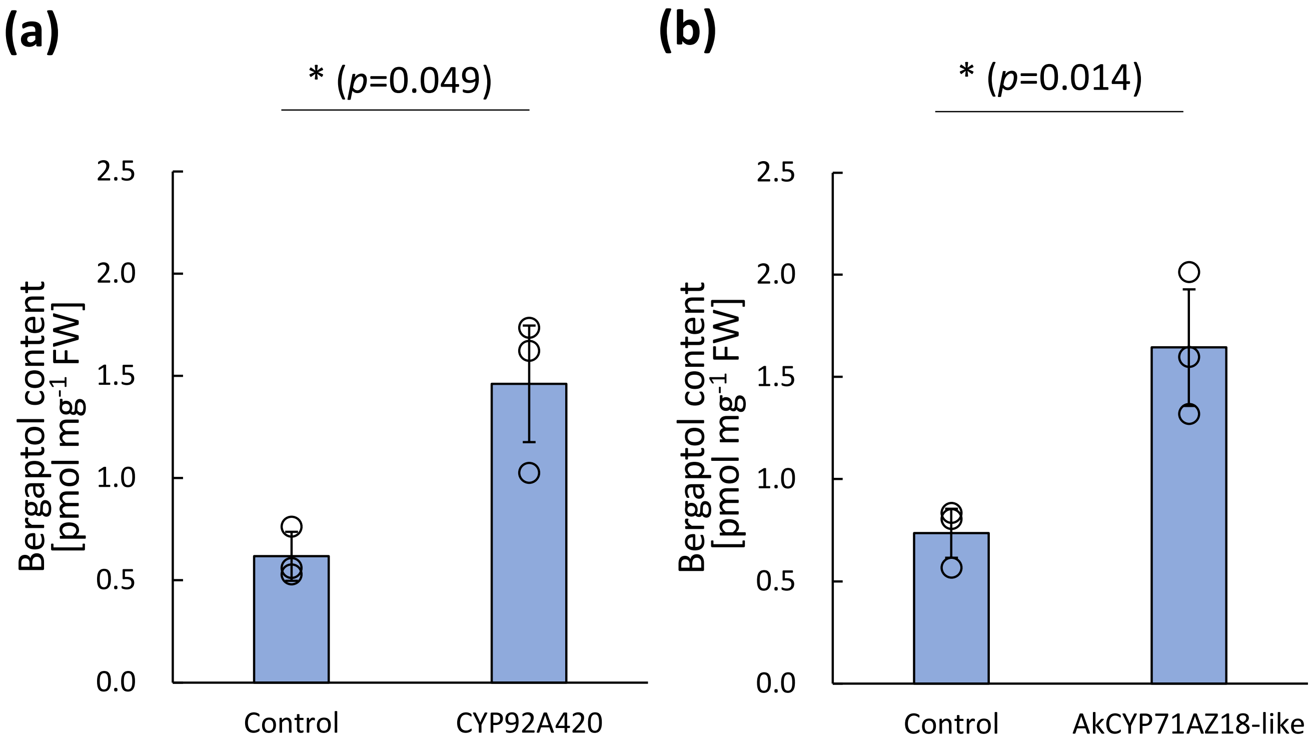


**Figure S18.** Psoralen 5-hydroxy activity of CYP92A470 and AkCYP71AZ18-like in *Nicotiana benthamiana* leaves.
Bergaptol content of *N. benthamiana* leaves expressing EGFP (control), CYP92A470 (a), and AkCYP71AZ18-like (b). Data are shown as mean ± standard deviation with three biological replicates ($n$ = 3). Statistical significance was evaluated using Welch's $t$ test.

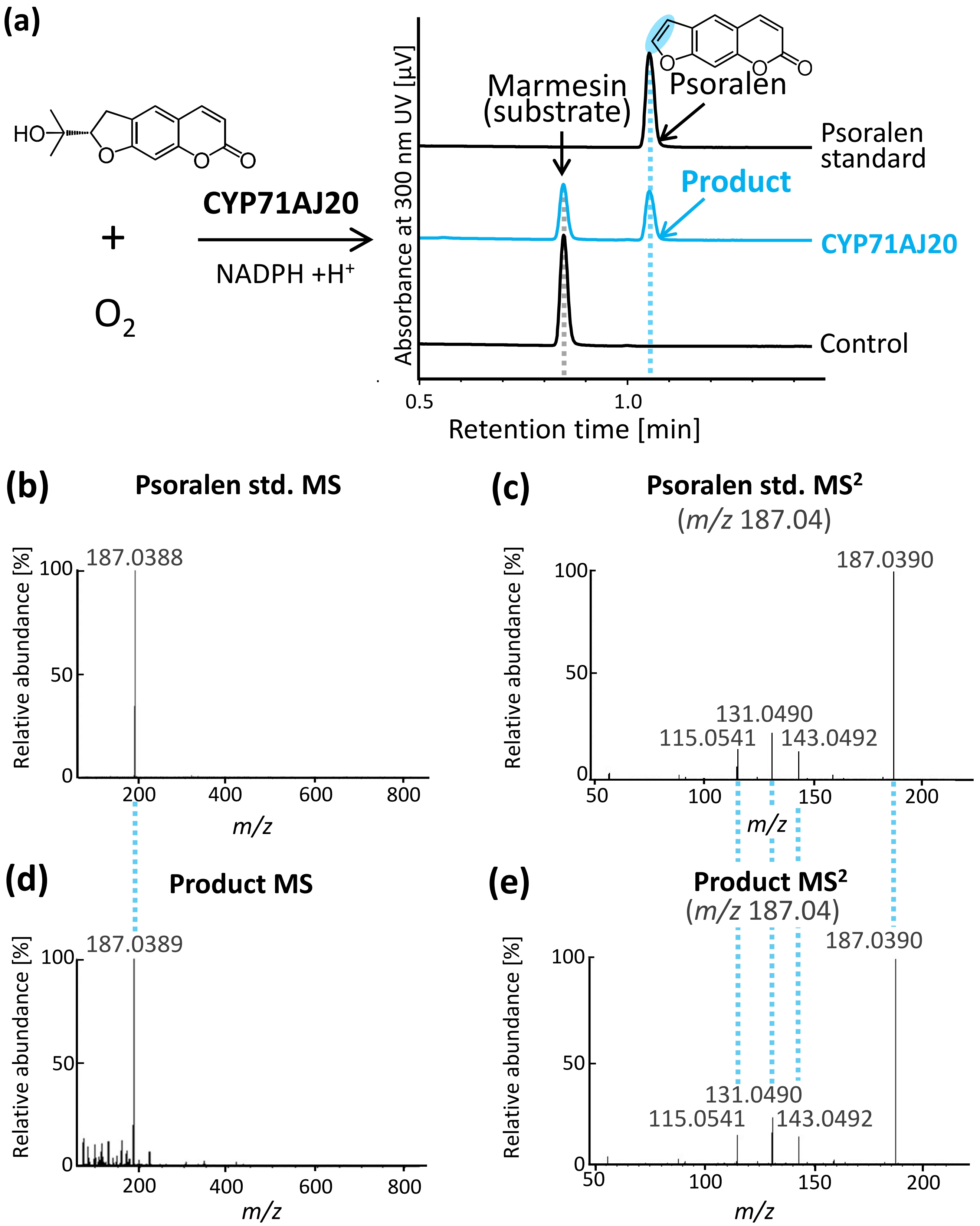


**Figure S19.** Psoralen synthase activity of CYP71AJ20.
(a) UV chromatogram of *in vitro* yeast microsome assay of CYP71AJ20 using marmesin as a substrate.
(b, d) MS fragmentation of psoralen standard (b) and CYP71AJ20 product (d).
(c, e) MS$^2$ fragmentation of psoralen standard (c) and the product (e) at *m/z* 187.04.

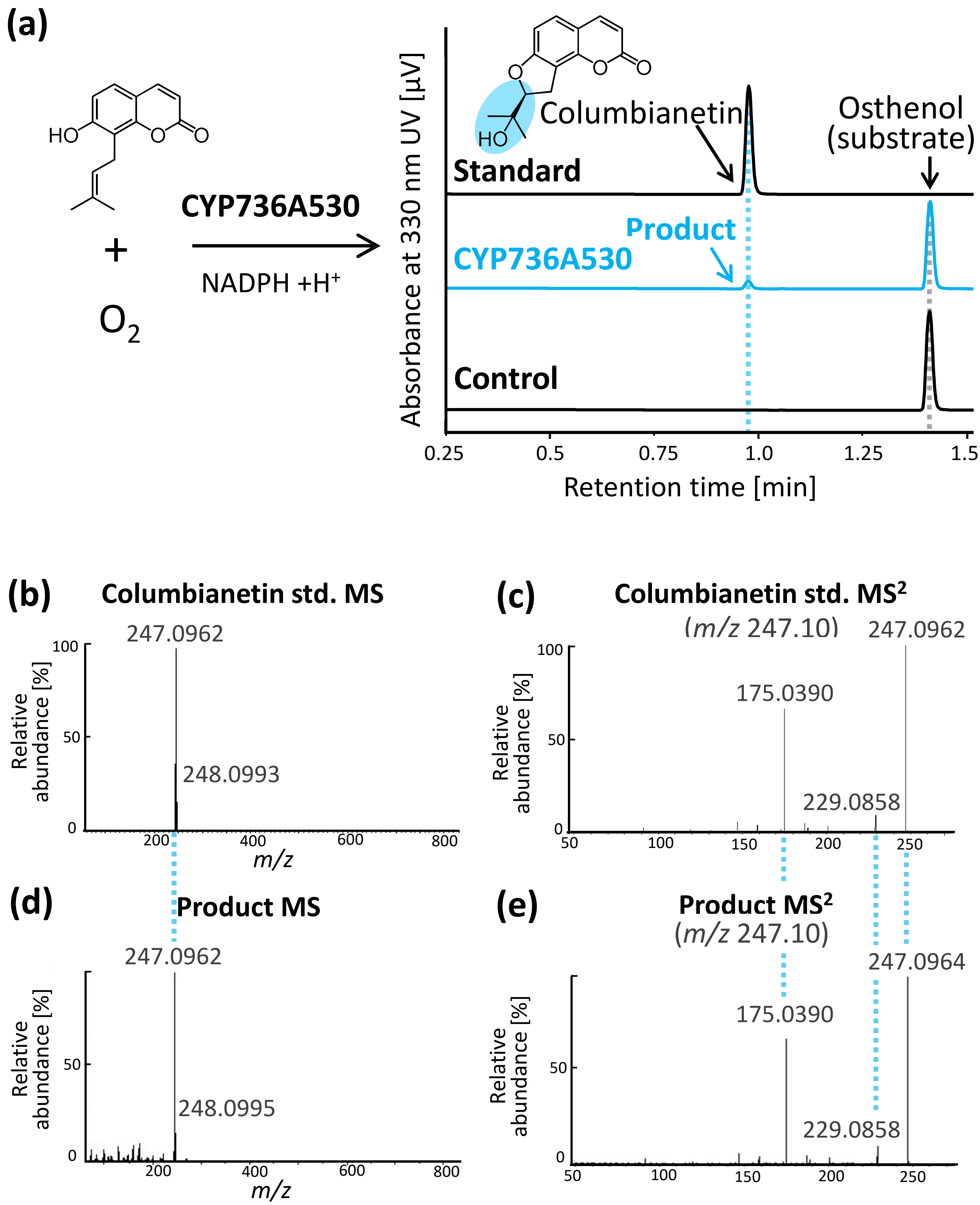


**Figure S20.** Columbianetin synthase activity of CYP736A530.
(a) UV chromatogram of *in vitro* yeast microsome assay of CYP736A530 and control using osthenol as a substrate.
(b, d) MS fragmentation of the columbianetin standard (b) and CYP736A530 product (d).
(c, e) $MS^2$ fragmentation of columbianetin standard (c) and the product (e) at *m/z* 247.10.

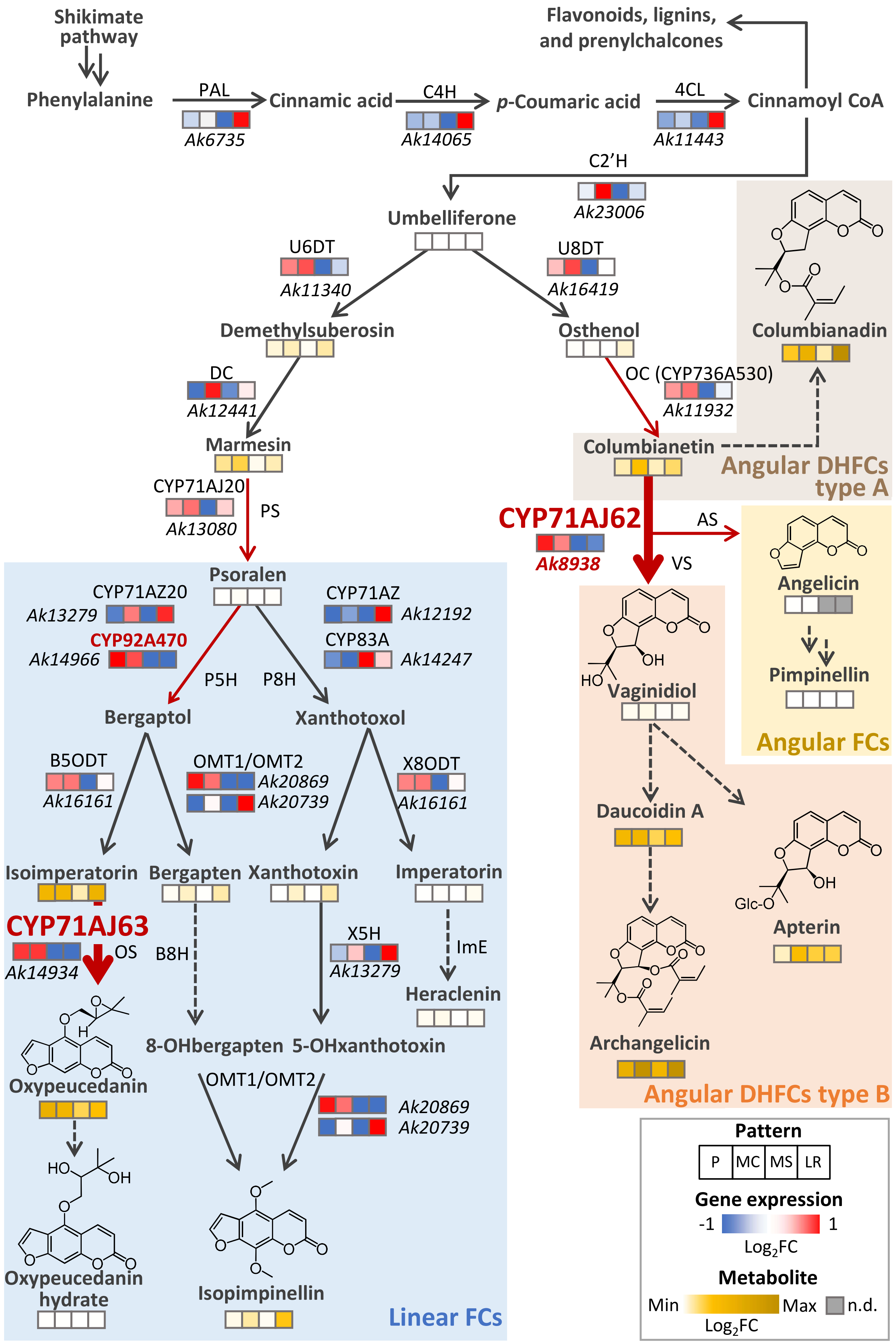


**Figure S21.** Biosynthetic pathway of TCs with transcriptome and metabolite data in *A. keiskei*..

Shinya *et al.*

**Figure S21.** Biosynthetic pathway of TCs with transcriptome and metabolite data in *A. keiskei*. The transcriptome and metabolite data were overlaid onto the TC biosynthetic pathway of Apiaceae. Each arrow represents a single enzymatic reaction. Cytochrome P450 reactions identified in this study are shown in red. Solid lines indicate known genes; and dashed lines indicate unknown genes. Gene expression heatmaps are based on mean-standardized $log_2$ fold changes. Metabolite accumulation levels are shown as mean-standardized $log_2$ values across all samples. Organs and metabolites that were not detected are indicated in gray (n.d.). PAL, phenylalanine ammonia-lyase; C4H, cinnamate 4-hydroxylase; 4CL, 4-coumaroyl: CoA ligase; C2'H, p-coumaroyl CoA 2'-hydroxylase; U6DT, umbelliferone 6-dimethylallyltransferase; U8DT, umbelliferone 8-dimethylallyltransferase; DC, demethylsuberosin cyclase; PS, psoralen synthase; VS, vaginidiol synthase; AS, angelicin synthase; P5H, psoralen 5-hydroxylase; P8H, psoralen 8-hydroxylase; OMT, *O*-methyltransferase; B5ODT, bergaptol 5-*O*-dimethylallyltransferase; X8ODT, xanthotoxol 8-*O*-dimethylallyltransferase; OS, oxypeucedanin synthase; ImE, imperatorin epoxidase; B8H, bergapten 8-hydroxylase; X5H, xanthotoxin 5-hydroxylase.

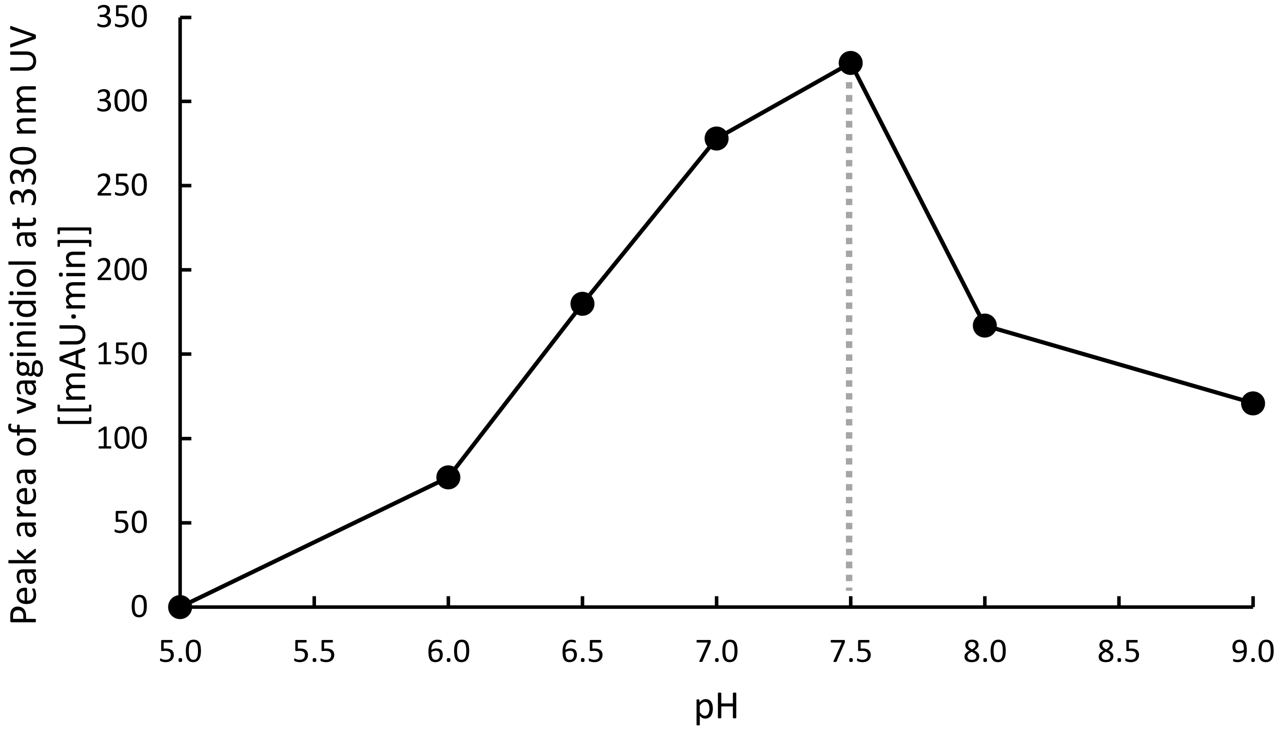


**Figure S22.** A preliminary analysis of the optimal pH for the vaginidiol synthesis activity of CYP71AJ62. Single reactions were performed at pH 5.0−9.0. The peak areas corresponding to vaginidiol formation were measured at 330 nm.

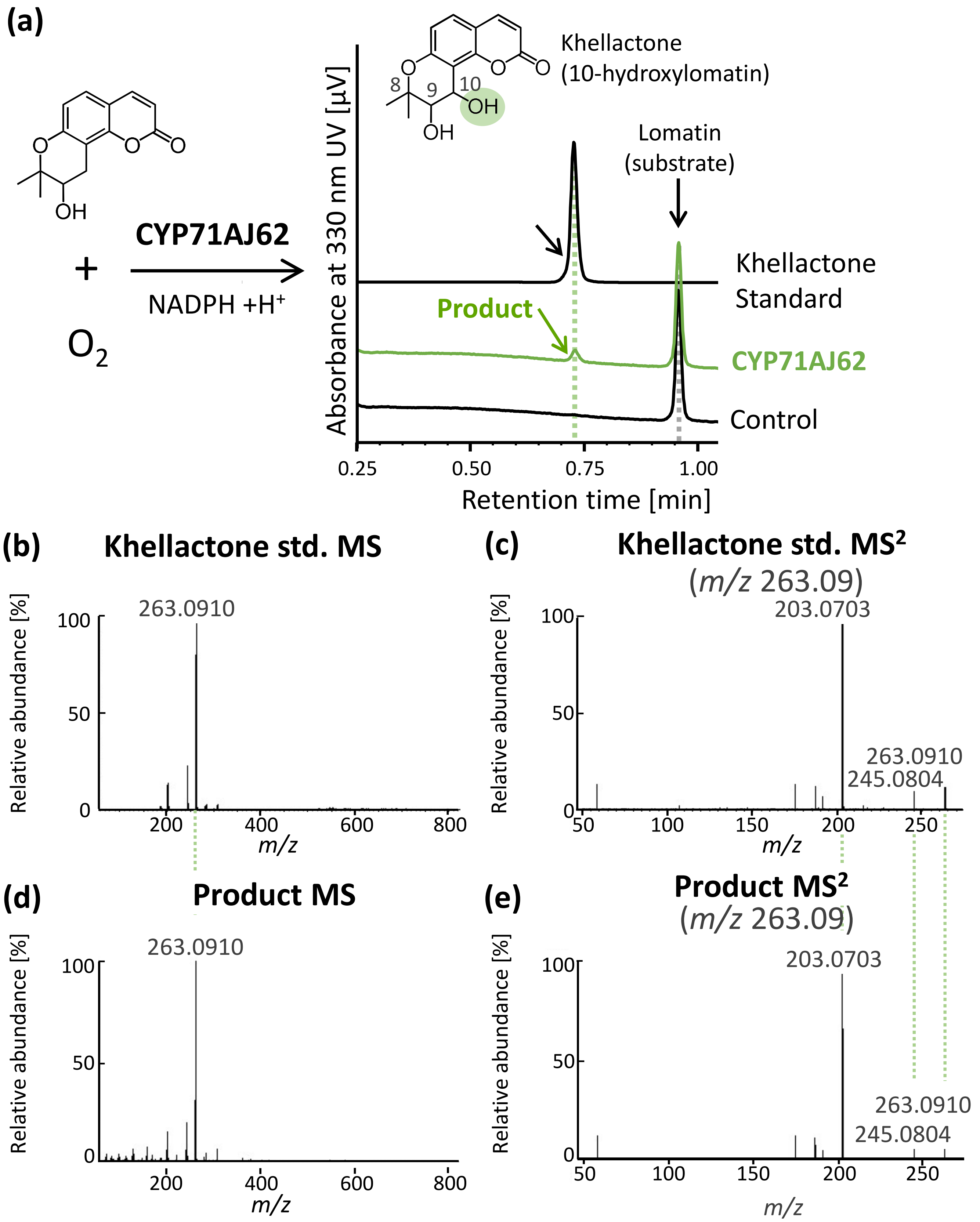


**Figure S23.** Lomatin 10-hydroxylation activity of CYP71AJ62.
(a) UV chromatogram of *in vitro* microsome assay of CYP71AJ62 and control using lomatin as a substrate.
(b, c) MS fragmentation of khellactone standard (b) and CYP71AJ62 product (c).
(d, e) $MS^2$ fragmentation of kellactone standard (d) and the product (e).

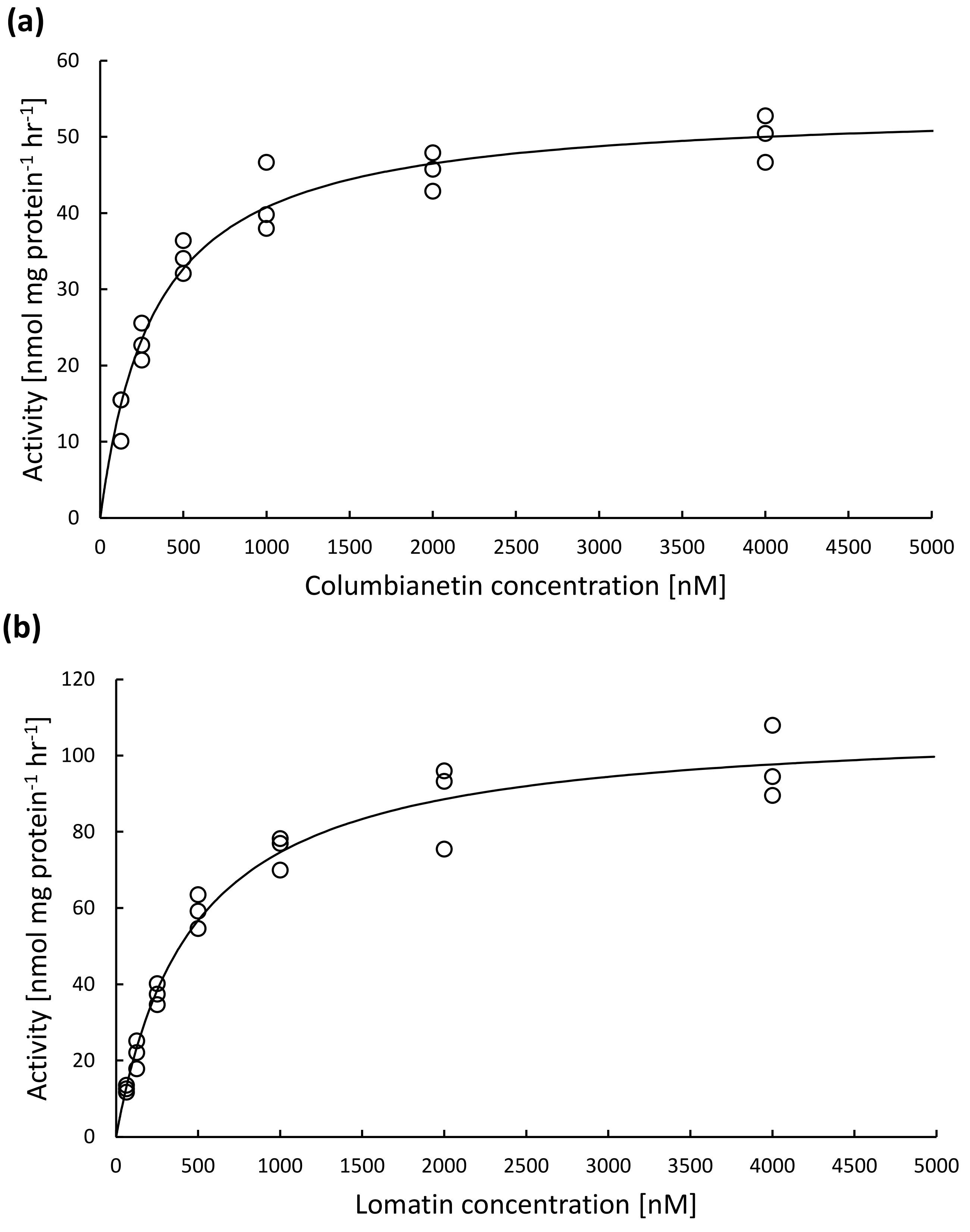


**Figure S24.** Kinetic analysis of CYP71AJ62 with columbianetin or lomatin as substrates.
Apparent $K_m$ values for columbianetin (a) and lomatin (b) were calculated using the nonlinear least squares fitting. Kinetic assays were independently repeated three time.

**(a)**

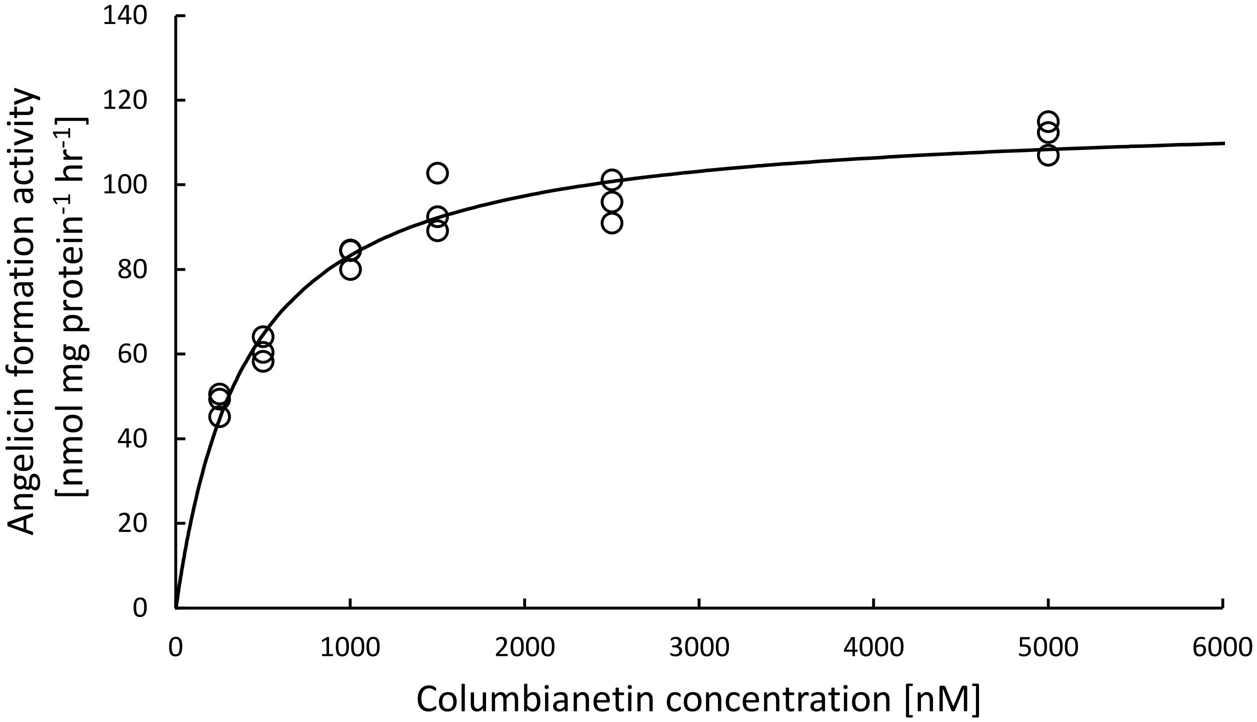


**(b)**

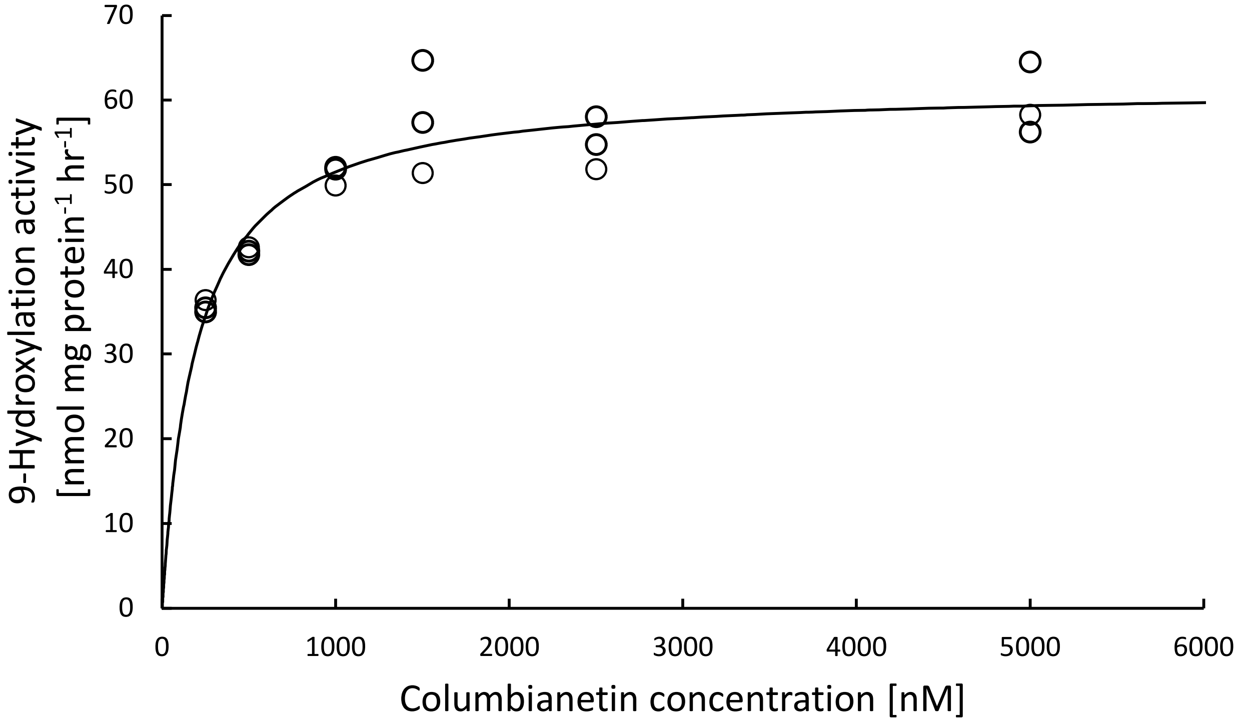


**Figure S25.** Kinetic analysis of CYP71AJ4 with columbianetin as a substrate. Apparent $K_m$ values for angelicin (a) and vaginidiol (b) production were calculated using the nonlinear least squares fitting. Kinetic assays were independently repeated three time.

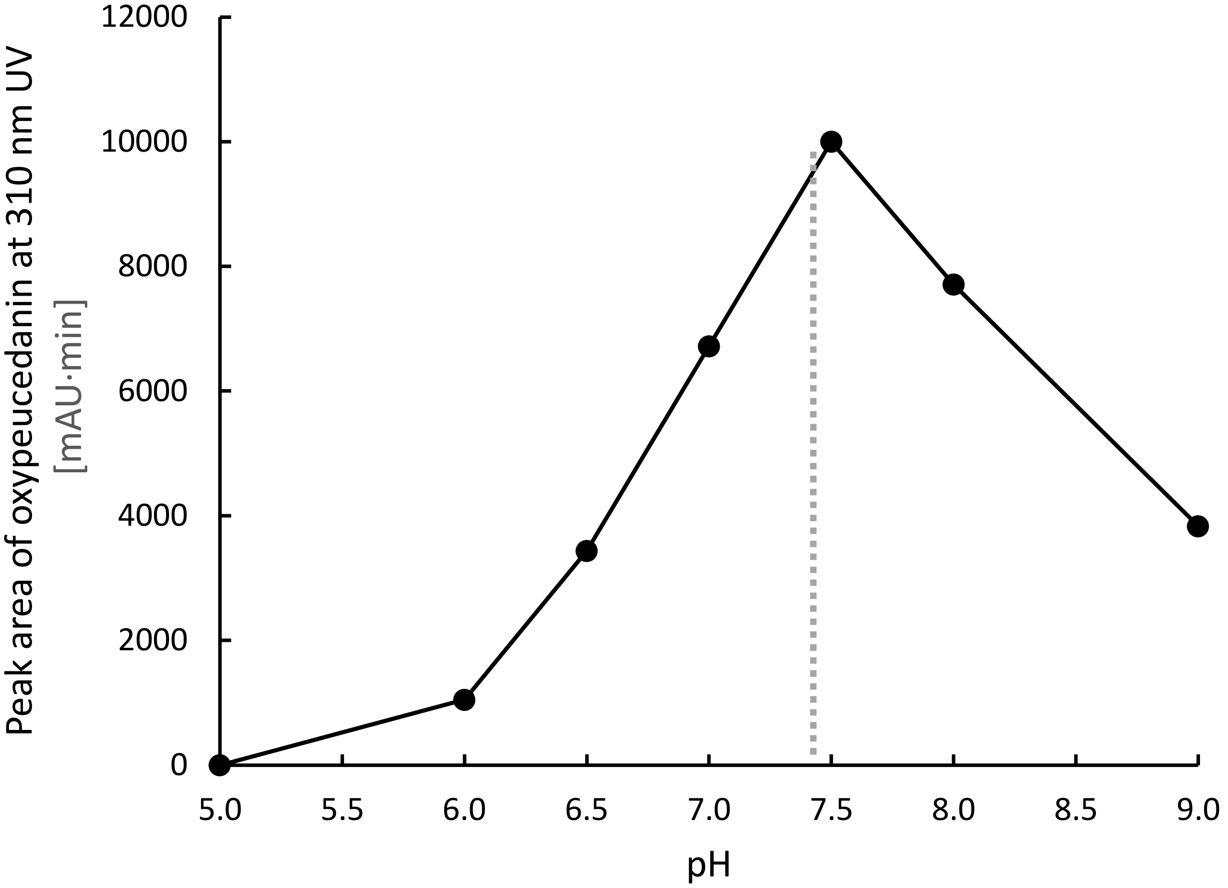


**Figure S26.** A preliminary analysis of the optimal pH for the oxypeucedanin synthesis activity of CYP71AJ63. Single reactions were performed at pH 5.0−9.0. The peak areas corresponding to the formation of oxypeucedanin were measured at 310 nm.

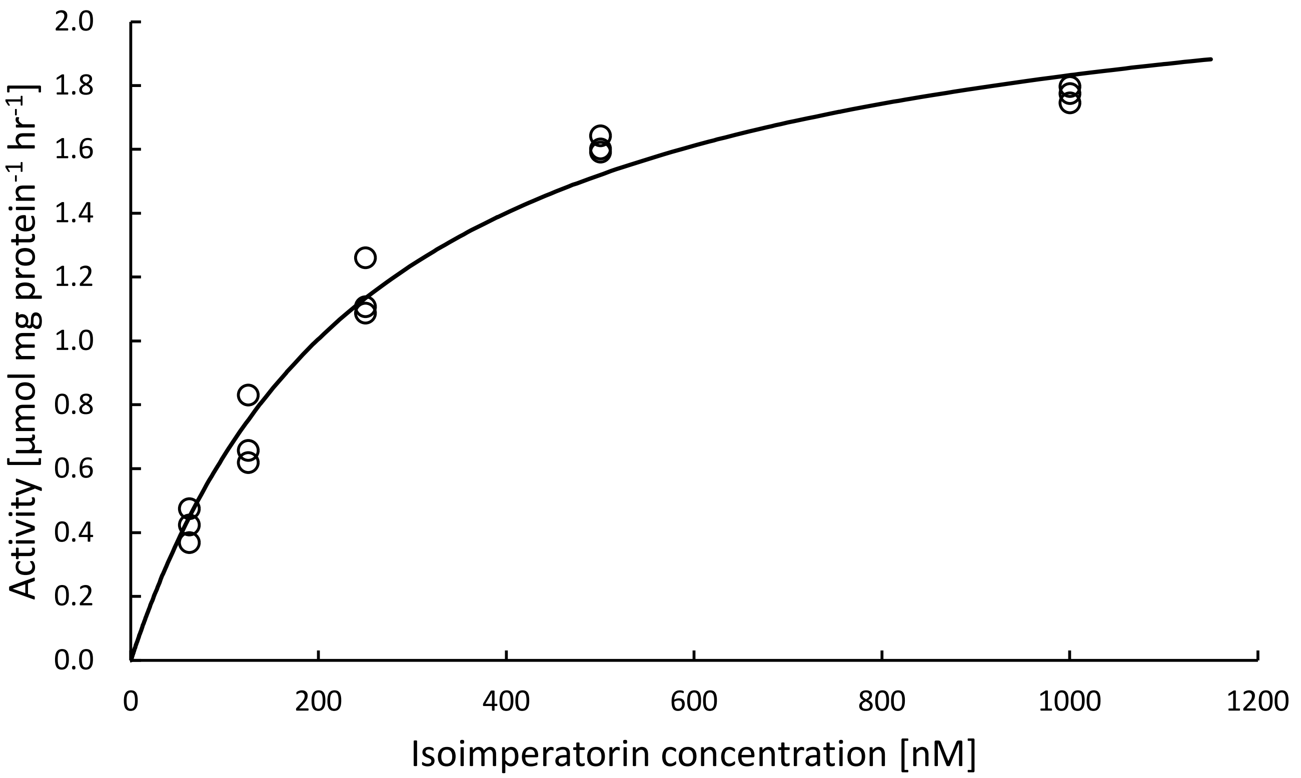


**Figure S27.** Kinetic analysis of CYP71AJ63 with isoimperatorin as a substrate. The apparent $K_m$ value for oxypeucedanin production was calculated using the nonlinear least squares fitting. Kinetic assays were independently repeated three time.

**(g)**

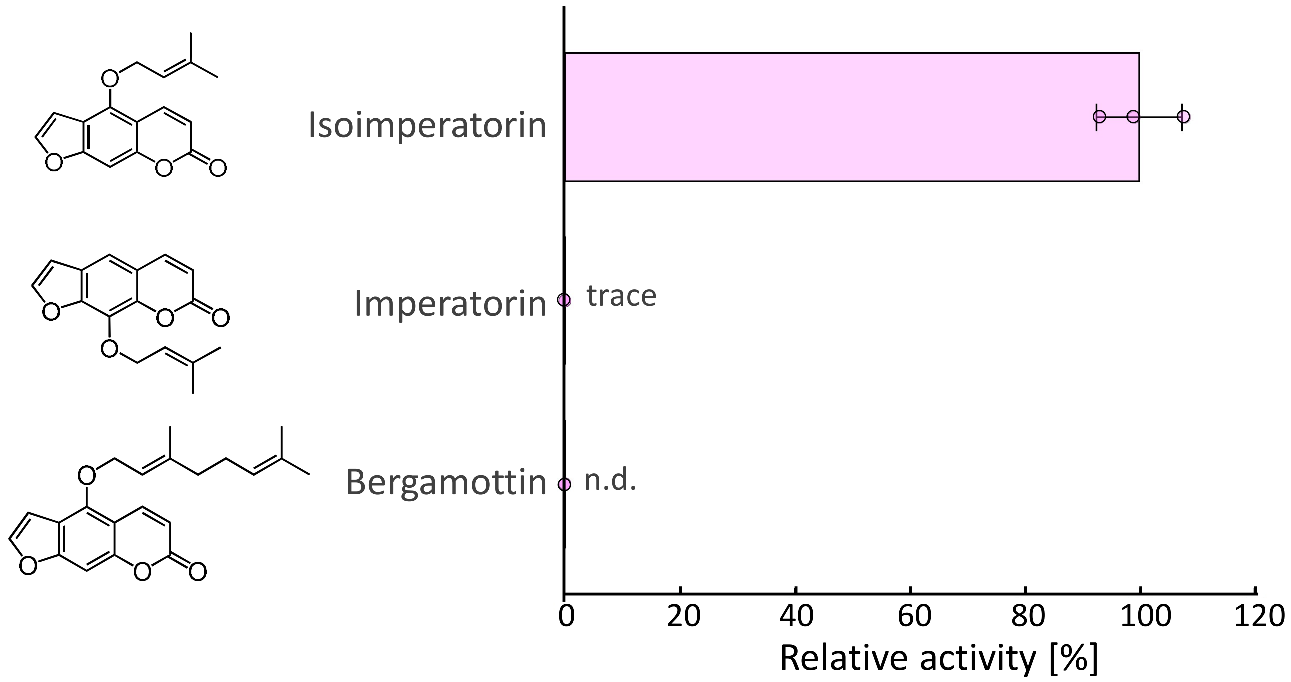


**Figure S28.** Substrate preference of CYP71AJ63.
The isoimperatorin epoxidase activity was set to 100 %. Trace means the consistent detection with control. Data are shown as mean ± standard deviation from three independent experiments. n.d., not detected.

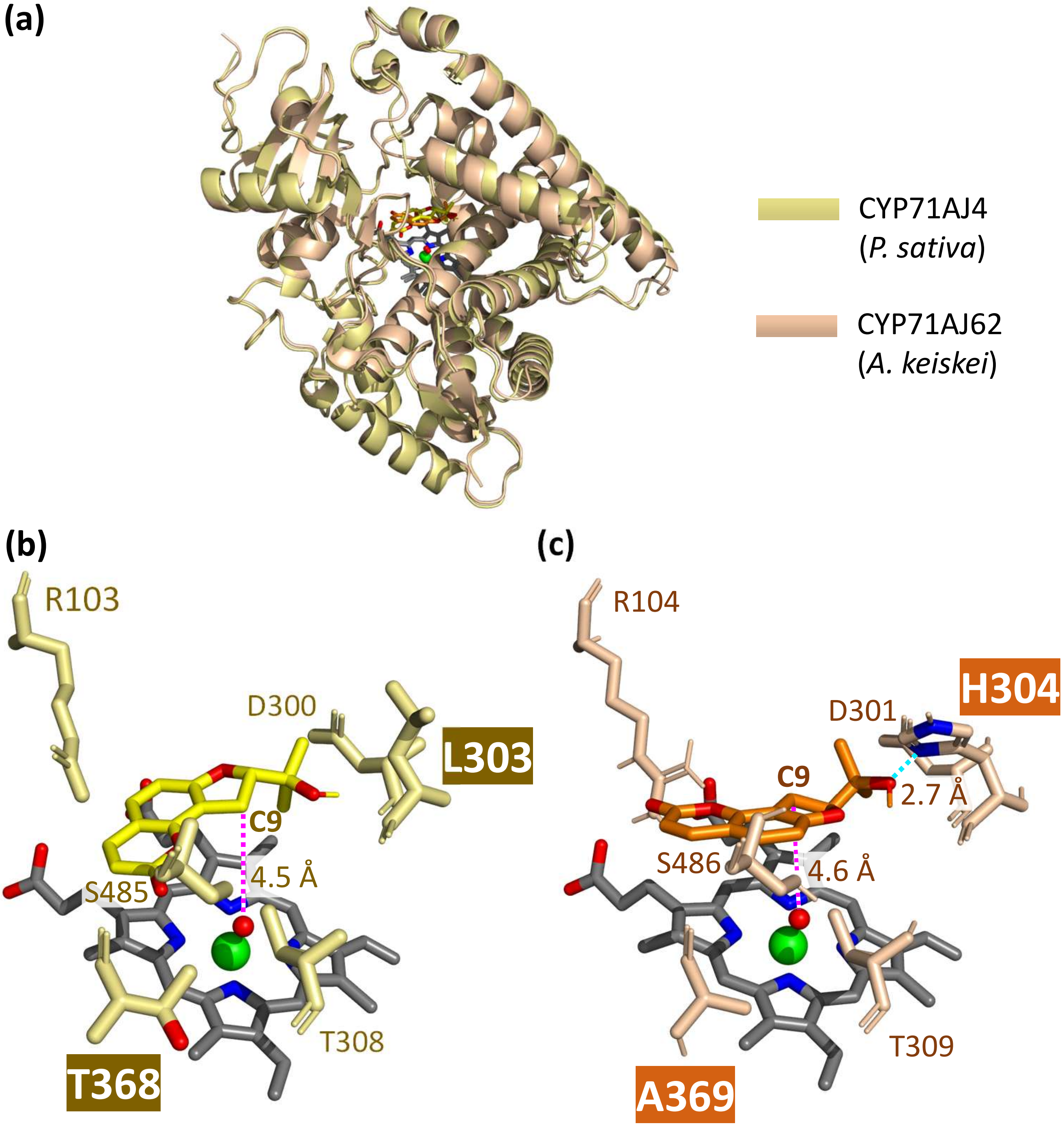


**Figure S29.** Three-dimensional homology models of CYP71AJ4 and CYP71AJ62. Three-dimensional homology models of CYP71AJ4 and CYP71AJ62 are shown in khaki and pale orange, respectively.
(a) Columbianetin docking with CYP71AJ4 and 62 is shown in yellow and orange, respectively. The heme and heme iron are shown in gray and lime, respectively. Oxygen and nitrogen atoms of the heme are shown in red and blue.
(b,c) Orientation of columbianetin toward the heme iron in CYP71AJ4 (b) and 62 (c). The heme-binding oxygen atom (red sphere) is set 1.7 Å from the heme iron (Rupasonghe et al., 2003). Distances from the heme-binding oxygen to C9 are shown.

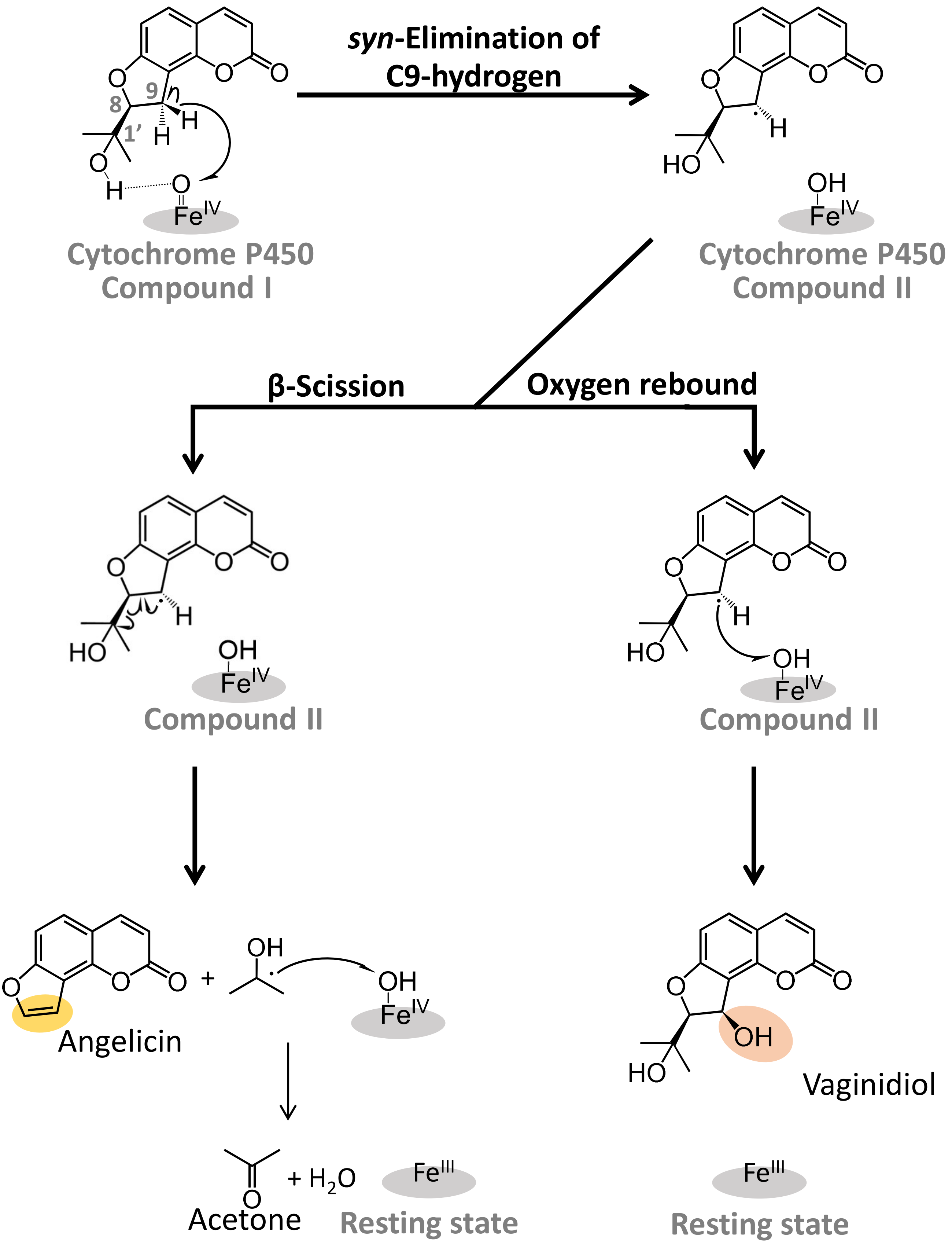


**Figure S30.** Presumed reaction mechanisms catalyzed by CYP71AJ4 (yellow) and CYP71AJ62 (orange). The angelicin biosynthetic pathway was inferred based on Stanjek et al. (1999) and Larbat et al. (2009).

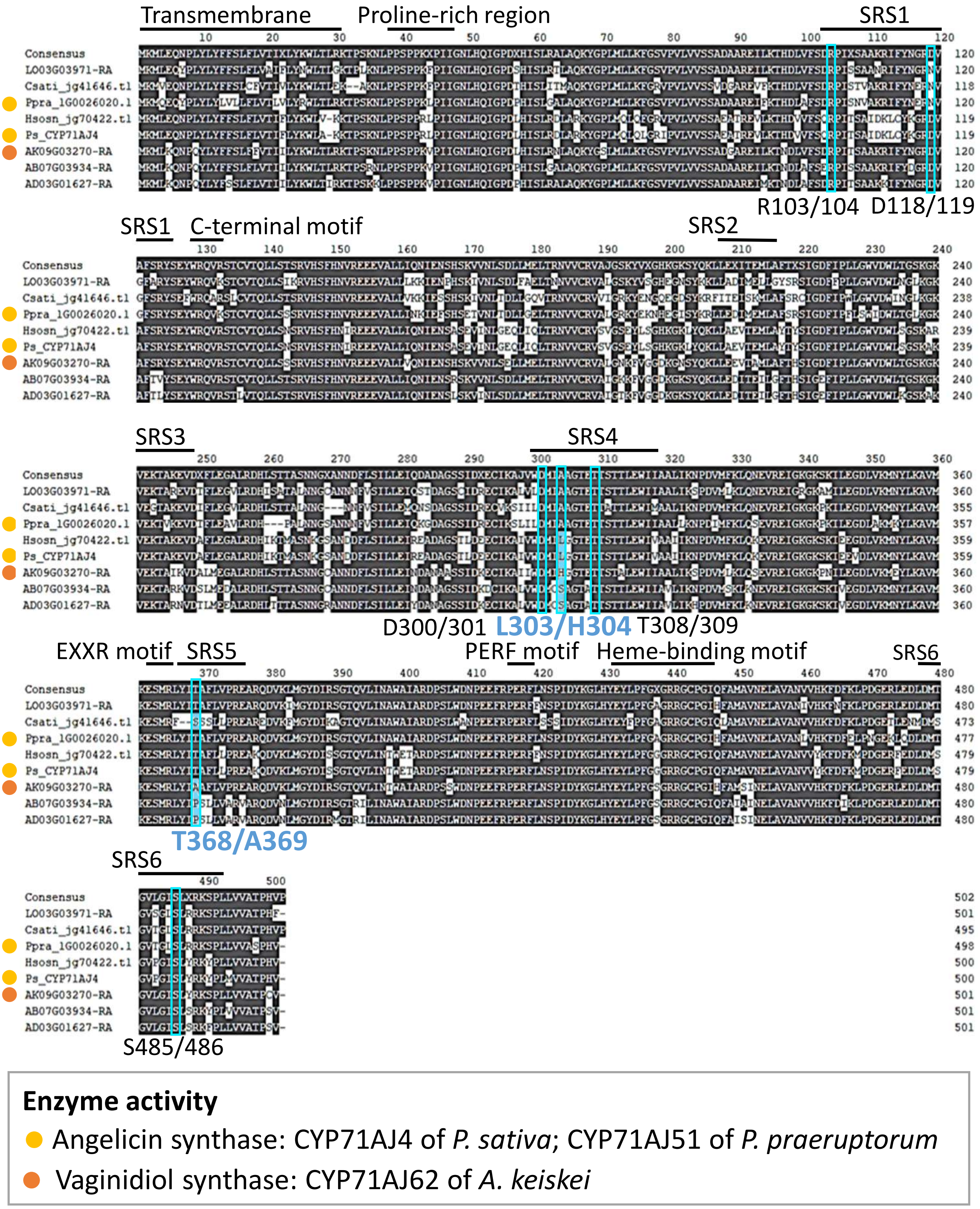


**Figure S31.** Sequence alignment of candidate P450s in the columbianetin clade of Fig. 5a. Amino acid sequences deduced from the 11 candidates were aligned using MAFFT v7.490. All analyses were performed using Geneious Prime® 2024.2.1 (Dotmatics). Known angelicin synthases are indicated by yellow circles, while the vaginidiol synthase of *A. keiskei* is indicated by orange circles. The SRSs and motifs were annotated according to Dueholm et al. (2015) and Fang et al. (2024). The transmembrane region was estimated using TargetP-2.0 (Armenteros et al., 2019).

Residues located within 5 Å of the substrate in the three-dimensional structural analysis are highlighted with cyan boxes.
