## Supplementary Data S1 for "Molecular characterization of a cytochrome P450 involved in the skeletal diversification of tricyclic coumarins in *Angelica keiskei*"

**Supplementary Data S1. Newick data for the 286 sequences selected based on OrthoFinder results.**

(AB01G01156-RA:0.0634944936,(AK10G04270-RA:0.0262107014,AK10G04268-
RA:0.0178215340):0.0114728124[92.2],(((As01G05552:0.0490429861,Ppra_7G0032780.1:0.03837
88503):0.0347190553[94.9],(AB10G00790-RA:0.0236230415,(AD10G00467-
RA:0.0000009998,AD10G00465-
RA:0.0000009998):0.0280005748[99.6]):0.0052786885[74.8]):0.0279457149[90.7],(LO09G04836-
RA:0.0273727730,((XP_017219080.1:0.0917275933,XP_017219081.1:0.0712482869):0.04011088
93[97.4],(((CACVNY010000006.1_RagTag.g83270:0.0465325068,Ppra_4G0014860.1:0.03724074
71):0.0478600077[99.9],(XP_017217982.1:0.1065684499,(XP_017218065.1:0.1067116299,(Csati_j
g22359.t1:0.0827322011,(As01G05549:0.0543378154,(Csati_jg22360.t1:0.0366107974,((AK06G0
0372-RA:0.1822049412,Ppra_7G0032750.1:0.0108379651):0.0072126041[68.1],(AK10G04266-
RA:0.0429674536,(Ppra_7G0032770.1:0.0492159617,AK10G04267-
RA:0.0170574192):0.0548531723[99.9]):0.0364559453[99.6]):0.0090628430[86.9]):0.0071954812[
86]):0.0157449132[83.4]):0.0272559682[20.7]):0.0522639078[99.2]):0.0123749308[68.2]):0.05268
56552[95.5],(Cas06G001983.t1:1.0587872116,(((XP_017254492.1:0.0955686626,((As10G04221:0.
0000009998,(AsUnG01147:0.0176716209,As10G04214:0.0000009998):0.0023071390[79.6]):0.030
5654446[98.9],((CACVNY010000010.1_RagTag.g152879:0.0324245053,As07G03136:0.04329990
90):0.0338060304[98],(Hsosn_jg20492.t1:0.0748469521,(Ppra_2G0008940.1:0.0093748461,(AK04
G01259-RA:0.0068174043,AK04G01251-
RA:0.0141356540):0.0269337085[98.6]):0.0443335340[99.8]):0.0146199153[91.2]):0.0236428568[
88.8]):0.0285394644[93.7]):0.2364922341[100],((pg_19000691:0.0822618195,(pg_19000697:0.008
3360655,pg_3012796:0.0036082508):0.1000010097[100]):0.0473024192[96.2],((XP_017218301.2:
0.1125471291,XP_017218300.1:0.1039970485):0.0226466619[64.1],(((LO09G04845-
RA:0.0227010732,As01G05553:0.0262468995):0.0117710254[94.2],(Csati_jg22349.t1:0.04152397
14,(Ppra_7G0032790.1:0.0393650968,AD10G00463-
RA:0.0453867520):0.0190842935[83.1]):0.0086720919[88.4]):0.0130281128[93.3],((LO09G04847
-RA:0.0127815201,LO06G02729-
RA:0.0236306483):0.0382306693[99.5],(ptg002559l_RagTag.g212921:0.0046338530,(CACVNY0
10000007.1_RagTag.g117445:0.0067643656,(atg027936l.g20569:0.0470524174,CACVNY0100000
06.1_RagTag.g83280:0.0000009998):0.0022440714[80.3]):0.0020832257[76.3]):0.0596746924[100
]):0.0216420665[97.2]):0.0211101284[91.3]):0.1840654912[100]):0.0657046806[96]):0.169094623
6[99.9],(((pg_19000698:0.0450517277,pg_3012795:0.0423154676):0.3108290863[100],((Sc6G333
8:0.2319072251,((Ppra_7G0032740.1:0.0298174592,(AK10G04263-
RA:0.0160436554,AB10G00794-

RA:0.0358136612):0.0130030364[45.6]):0.1083485032[100],((CACVNY010000782.1_RagTag.g18
3689:0.0806475948,((Csati_jg14599.t1:0.0000009998,Csati_jg41199.t1:0.0000009998):0.04596832
87[100],(LO03G03843-
RA:0.0332588754,As04G04540:0.0169067843):0.0045351470[84.5]):0.0058153804[65.3]):0.04627
48783[99.5],(Csati_jg22363.t1:0.0543810476,(Csati_jg22364.t1:0.0744232586,(As01G05548:0.044
9921218,(AB10G05428-RA:0.0591970920,(AK10G04264-RA:0.0151261426,AB10G00793-
RA:0.0268922267):0.0148382569[78.8]):0.0427798298[99.3]):0.0154915495[46]):0.0177271791[9
1.5]):0.0845328747[100]):0.0512085507[96.7]):0.1620451566[100]):0.0251250740[30.3],((Sc2G05
66:0.1810096382,(Csati_jg64783.t1:0.1045805878,(((Ls_CYP71AJ15:0.0284849813,Dc_CYP71AJ
7:0.0529840776):0.0284634275[98.6],(As02G00724:0.0388404360,(Hm_CYP71AJ11:0.024358955
0,((Pc_CYP71AJ9:0.0402295619,Am_CYP71AJ8:0.0373150866):0.0185180720[97.8],(Ppra_3G00
24370.1:0.0175981360,AB08G03833-
RA:0.0502418928):0.0137671164[94.4]):0.0054787814[78.7]):0.0113270242[87.8]):0.0157859037[
95.1]):0.0294092108[98.3],((XP_017241292.1:0.0817309349,(Csati_jg64781.t1:0.0310103494,CA
CVNY010000011.1_RagTag.g177817:0.0693228877):0.0245356631[95.7]):0.0685924721[100],(Cs
ati_jg64776.t1:0.0537129610,((Csati_jg64774.t1:0.0278573189,(CACVNY010000011.1_RagTag.g
177813:0.0315299740,(LO04G01224-
RA:0.0348848717,As02G00727:0.0167912367):0.0161977140[93.8]):0.0049384885[21]):0.003444
1774[46.9],(Ps_CYP71AJ13:0.0258820406,((XP_017240590.1:0.0371951589,XP_017239051.1:0.0
106798984):0.0478393729[99.9],(Ppra_3G0024380.1:0.0147256435,(AB08G03834-
RA:0.0123559861,(AK08G01511-RA:0.0249670181,AD11G03021-
RA:0.0222791998):0.0059585976[7.5]):0.0102655267[85.9]):0.0133408509[93.1]):0.0090683214[8
9.9]):0.0004510681):0.0221209432[97.1]):0.0426736603[99.9]):0.0135230985[90.9]):0.021708571
6[73.2]):0.1269217673[99.9]):0.0663927151[98.8],((XP_017217855.2:0.0671772534,(Csati_jg2845
4.t1:0.0177912048,((LO09G01532-
RA:0.0124093081,As01G02641:0.0202501149):0.0233918497[98.1],(Ppra_3G0004310.1:0.015457
8482,((AD06G01905-RA:0.0180355475,AB08G01524-
RA:0.0166208582):0.0051695134[73.3],(Kp_CYP71AJ50:0.0159768300,(AK07G03618-
RA:0.0076055407,AK07G03615-
RA:0.0050875396):0.0051983711[87]):0.0030165434[65.6]):0.0158533745[96]):0.0368544845[100
]):0.0045842338[43.7]):0.0326877607[97.4]):0.0993116662[100],((XP_017219082.2:0.1908536508
,(LO09G04827-RA:0.0599017800,(Ppra_4G0014870.1:0.3480676018,((AB04G03112-
RA:0.0448424566,AK05G02327-RA:0.0190428156):0.0169573762[95.5],(AK03G06066-
RA:0.0152107718,(AB01G06522-RA:0.0000009998,AB01G06518-
RA:0.0023055769):0.0250087291[99.6]):0.0159300363[95.3]):0.0151158326[71.2]):0.0643323273[
99.9]):0.0411926774[97.4]):0.0803203807[100],((XP_017216421.1:0.1083648487,(Csati_jg22367.t

1:0.0550901514,LO09G04822-
RA:0.0242535573):0.0607689526[99.8]):0.0558217014[98.4],(Dc_CYP71AJ21:0.1405854835,(((at
g013256l.g12305:0.0075208748,atg006713l.g6615:0.0061094896):0.0876946174[100],(AK05G008
18-
RA:0.0569679525,(Ppra_9G0001380.1:0.0413070057,(Ppra_4G0005250.1:0.0381122706,(KpPS_C
YP71AJ49:0.0610859668,Kp_CYP71AJ48:0.0213152725):0.0031807182[74.4]):0.0099405567[20.
1]):0.0063377874[68.3]):0.0325901083[98.7]):0.0215415929[97.2],(((Ppra_1G0025390.1:0.014792
2043,(AD03G01709-RA:0.0069562523,(AK09G03210-RA:0.0139560137,AB07G04022-
RA:0.0285908907):0.0000022299):0.0066147649[84.2]):0.0877402539[100],((LO03G03971-
RA:0.0809296286,(Csati_jg41646.t1:0.1554588056,KpAS_CYP71AJ51:0.0831785294):0.0382844
056[96.7]):0.0236043640[0.9],((PsAS_CYP71AJ4:0.0099990552,Hsosn_jg70422.t1:0.0041132761)
:0.1961411943[100],(AK09G03270-RA:0.0634450343,(AD03G01627-
RA:0.0387995450,AB07G03934-
RA:0.0270800504):0.0799280296[100]):0.0405414664[87.3]):0.0722433309[99.9]):0.0119563752[
26]):0.0281233609[93.6],((LO06G03500-RA:0.0023775844,LO03G03969-
RA:0.0047342506):0.0179174656[97.8],(As04G04982:0.0099354380,((CACVNY010000007.1_Ra
gTag.g109818:0.0257174185,(AgPS_CYP71AJ2:0.0831260360,AmPS_CYP71AJ1:0.0408168668):
0.0157135011[76.9]):0.0769450346[100],((PsPS_CYP71AJ3:0.0046594366,Hsosn_jg70421.t1:0.00
50942553):0.1329201318[100],(AD03G01634-RA:0.0052167847,(AB07G03927-
RA:0.0098580189,(AK09G03269-
RA:0.0073131563,(Ppra_1G0026000.1:0.0000009998,Ppra_1G0025990.1:0.0000009998):0.017072
4067[99.5]):0.0047377740[84.1]):0.0070629257[89.5]):0.0382549240[97.5]):0.0209629133[82.3]):
0.0257232348[97]):0.0045947962[79.3]):0.0953022045[100]):0.0031862640[74.1]):0.0414362408[
97.5]):0.0706747912[98.2]):0.0313005183[94.8]):0.0207470889[83.4]):0.1252196660[100]):0.0278
082227[89]):0.1418254176[99.8]):0.1724333596[99.1],((pg_19000689:0.2348988516,((Sc4G0274:
0.1202951658,((XP_017233626.2:0.1080027697,(AK03G00664-RA:0.0282675065,AD09G04737-
RA:0.0719409862):0.0564489290[99.6]):0.0551869718[98.1],((Ppra_9G0004670.1:0.0553868682,(
XP_017232654.1:0.1669413555,(Csati_jg10331.t1:0.0553186447,Ppra_9G0004690.1:0.039546465
5):0.0157150708[83.4]):0.0226867280[84.5]):0.0434394725[98.1],((XP_017232783.1:0.095843019
5,XP_017232483.1:0.0642291222):0.0825595149[100],(Csati_jg10333.t1:0.2396472674,((Csati_jg
10332.t1:0.0475432901,(ptg017489l_RagTag.g297314:0.0545811633,(LO10G05523-
RA:0.0113668732,As03G05957:0.0259345036):0.0298296977[99.6]):0.0111714893[80.5]):0.03521
98398[99.4],((Ppra_9G0004680.1:0.0044250977,Ppra_9G0004660.1:0.0111827616):0.0213121202[
98],(AK03G00665-RA:0.0241542595,AD09G04738-
RA:0.0526361892):0.0167343662[93.9]):0.0399333591[99.7]):0.0058201067[14.9]):0.0350311413[
97]):0.0382800168[69.6]):0.0241617296[66.5]):0.0970667277[100]):0.0101723892[79.6],(Sc4G02

72:0.2553879105,((Sc6G3334:0.0545706061,(Sc4G0278:0.1042867807,Sc4G0277:0.0347436285):
0.0405431811[93]):0.2126369035[100],(Sc6G3335:0.1819021492,((XP_017254697.1:0.153364448
6,(LO01G03606-
RA:0.0682247973,(Csati_jg71751.t1:0.0801629436,Csati_jg71747.t1:0.0482574914):0.0124270309
[87.8]):0.0173689119[80.2]):0.0369338638[96.5],((XP_017245070.1:0.1619339401,XP_017256944
.1:0.1321934279):0.0258064845[43],((AD04G01741-RA:0.0270157459,(AK11G00296-
RA:0.0197310427,(AB06G05933-RA:0.0026125642,AB06G05932-
RA:0.0046273847):0.0239850630[98.9]):0.0020202646[21.4]):0.1121495851[100],(Csati_jg71749.t
1:0.0582548261,(AB09G01975-
RA:0.0389199134,(Ppra_2G0011840.1:0.0365657595,(AD07G01764-
RA:0.0120581045,(AK04G01627-RA:0.0342267656,AB09G01978-
RA:0.0245846484):0.0122882599[39.2]):0.0023647278[70.1]):0.0361553336[99.3]):0.0247221950[
94.2]):0.0195906732[95.9]):0.0061263574[57.8]):0.0635192469[100]):0.0940872170[100]):0.0400
134674[81]):0.0212293765[72.7]):0.0414761731[97.1]):0.0496018591[90.6]):0.4146132403[100],((
(Sc1G1966:0.1241238832,(Sc8G2325:0.1120621862,Sc8G2316:0.0874395099):0.0229453310[63.8
]):0.0696864974[88.4],(Csati_jg55875.t1:0.1493557589,((((XP_063941415.1:0.0305333133,XP_01
7229157.1:0.0553508561):0.0600442915[99.8],((AK06G01613-RA:0.0000009998,AK06G01612-
RA:0.0000009998):0.0226905997[98.2],(Ppra_11G0011870.1:0.0204893264,As10G01387:0.03086
79298):0.0081326379[38.3]):0.0279140506[95.5]):0.0684181613[99.9],((Csati_jg55883.t1:0.04130
62299,As10G01392:0.0256120862):0.0065089936[8.9],(Ppra_11G0011890.1:0.0236262307,(AK06
G01626-RA:0.0417152145,(AD08G01768-RA:0.0243208358,AB05G04314-
RA:0.0070745926):0.0106749676[78.9]):0.0093557125[91.6]):0.0180711579[97.3]):0.0740810342[
100]):0.0368854754[98.5],((Csati_jg55882.t1:0.0655836379,(LO05G03523-
RA:0.0208005347,As10G01391:0.0207786828):0.0462836316[99.3]):0.0686576043[99.9],(Csati_j
g55881.t1:0.0558342828,((Hsosn_jg61969.t1:0.0433623159,(Hsosn_jg62005.t1:0.0000025510,Hsos
n_jg62006.t1:0.0139545099):0.0025022301[11.7]):0.0272940129[99.5],(As10G01390:0.041486976
6,(Ppra_11G0011880.1:0.0148979937,(AK06G01625-RA:0.0455648555,(AD08G01769-
RA:0.0301831563,(AK06G01624-RA:0.0186643920,(AB05G04307-
RA:0.0189311822,(AB05G04405-RA:0.0023226527,AB05G04299-
RA:0.0000026760):0.0160828468[94.4]):0.0098078227[92.5]):0.0015002870[69.1]):0.0022838595[
76.2]):0.0035003515[76.5]):0.0199495366[97.6]):0.0038274796[39.3]):0.0079967872[74.2]):0.040
3451820[99.5]):0.0108102003[10.9]):0.0211761991[61.1]):0.1007924686[98.8]):0.6875902468[100
],((XP_017250493.1:0.0593729698,(LO06G00715-
RA:0.0189800823,(Ppra_7G0005530.1:0.0162324806,(Csati_jg78838.t1:0.0283247547,Hsosn_jg56
968.t1:0.0234548256):0.0045277465[82.3]):0.0022440991[36.3]):0.0190269470[83.3]):0.51582514
96[100],((((pg_24007881:0.0092659961,pg_23000926:0.0019265223):0.0665963072[99.3],((pg_24

007878:0.0000025510,pg_23000896:0.0162289775):0.0475425940[99.5],(pg_24007904:0.0148842
192,pg_23000295:0.0087082389):0.0385271852[98.7]):0.0257223718[44.7]):0.4707678503[100],(p
g_20006105:0.1275831835,(Sc8G0807:0.0792410844,(XP_017248439.1:0.0615923016,((LO08G00
243-
RA:0.0182899526,As08G00251:0.0135196682):0.0076028754[88],((Ppra_10G0034540.1:0.037398
4874,AK01G02894-RA:0.0165590180):0.0022333505[39.2],(AD03G05488-
RA:0.0000009998,AD03G05486-
RA:0.0022819750):0.0094848322[94.8]):0.0130985638[94.9]):0.0178453314[91.5]):0.0841131983[
100]):0.0587355148[97.8]):0.2102855265[100]):0.0420222401[32.5],((Sc2G1661:0.4988780118,pg
_11011005:0.3710953197):0.0780716833[90.9],((pg_12009828:0.1280777190,(XP_017248319.1:0.
0942761627,(Ppra_10G0029050.1:0.0348071911,(Csati_jg52156.t1:0.0045832153,Csati_jg52147.t1
:0.0000009998):0.0387731560[99.4]):0.0537969287[99.1]):0.1645143412[100]):0.2363779999[100
],(((Csati_jg38585.t1:0.0702491620,XP_017251851.1:0.1124217082):0.1601234498[100],(Sc8G05
42:0.2516947001,(XP_017248478.1:0.1947243990,((XP_063947372.1:0.0000009998,XP_0172483
18.1:0.0000009998):0.1419745959[100],(LO08G00850-
RA:0.0315449034,(Csati_jg52157.t1:0.0593643596,(Ppra_10G0029060.1:0.0048077159,AK01G03
511-
RA:0.0073966664):0.0117992957[90.5]):0.0106944362[88.4]):0.0684904267[99.7]):0.1092204299[
100]):0.0359866479[93.7]):0.0187723709[81.6]):0.0668400768[89.9],((Csati_jg52161.t1:0.0935211
825,(As08G00803:0.0747062860,(LO08G00849-
RA:0.0139930723,As08G00806:0.0071380378):0.0365043320[99.9]):0.0064681959[81.3]):0.01161
20932[88.2],((XP_017248491.2:0.0000009998,(XP_063947761.1:0.0000009998,(XP_063947762.1:
0.0000000000,XP_063947760.1:0.0000000000):0.0000009998):0.0000009998):0.0352505128[99.9
],(Csati_jg52160.t1:0.0076045471,(Hsosn_jg49290.t1:0.4051056130,(As08G00802:0.0134085268,(
AD03G04764-RA:0.0090383649,AB08G00004-
RA:0.0145442911):0.0098255778[92.1]):0.0000024335):0.0026851208[68.3]):0.0147424615[96]):0
.0076537060[74.4]):0.2461231852[100]):0.2188500645[100]):0.1115444021[97.3]):0.0776473367[
85.7]):0.4005333873[100]):0.0699868902[62.4]):0.0687663123[36]):0.3563076527[100]):0.137865
3994[98.4]):0.1742341915[98.6]):0.0724367493[89.9]):0.0772754363[99.6]):0.0154135431[86.8]):
0.0120915038[75.7]):0.0131394559[92.9]);
